# An integrative ENCODE resource for cancer genomics

**DOI:** 10.1101/706424

**Authors:** Jing Zhang, Donghoon Lee, Vineet Dhiman, Peng Jiang, Jie Xu, Patrick McGillivray, Hongbo Yang, Jason Liu, William Meyerson, Declan Clarke, Mengting Gu, Shantao Li, Shaoke Lou, Jinrui Xu, Lucas Lochovsky, Matthew Ung, Lijia Ma, Shan Yu, Qin Cao, Arif Harmanci, Koon-Kiu Yan, Anurag Sethi, Gamze Gursoy, Michael Rutenberg Schoenberg, Joel Rozowsky, Jonathan Warrell, Prashant Emani, Yucheng T. Yang, Timur Galeev, Xiangmeng Kong, Shuang Liu, Xiaotong Li, Jayanth Krishnan, Yanlin Feng, Juan Carlos Rivera-Mulia, Jessica Adrian, James R Broach, Michael Bolt, Jennifer Moran, Dominic Fitzgerald, Vishnu Dileep, Tingting Liu, Shenglin Mei, Takayo Sasaki, Claudia Trevilla-Garcia, Su Wang, Yanli Wang, Chongzhi Zang, Daifeng Wang, Robert Klein, Michael Snyder, David M. Gilbert, Kevin Yip, Chao Cheng, Feng Yue, X. Shirley Liu, Kevin White, Mark Gerstein

## Abstract

ENCODE comprises thousands of functional genomics datasets, and the encyclopedia covers hundreds of cell types, providing a universal annotation for genome interpretation. However, for particular applications, it may be advantageous to use a customized annotation. Here, we develop such a custom annotation by leveraging advanced assays, such as eCLIP, Hi-C, and whole-genome STARR-seq on a number of data-rich ENCODE cell types. A key aspect of this annotation is comprehensive and experimentally derived networks of both transcription factors and RNA-binding proteins (TFs and RBPs). Cancer, a disease of system-wide dysregulation, is an ideal application for such a network-based annotation. Specifically, for cancer-associated cell types, we put regulators into hierarchies and measure their network change (rewiring) during oncogenesis. We also extensively survey TF-RBP crosstalk, highlighting how SUB1, a previously uncharacterized RBP, drives aberrant tumor expression and amplifies the effect of MYC, a well-known oncogenic TF. Furthermore, we show how our annotation allows us to place oncogenic transformations in the context of a broad cell space; here, many normal-to-tumor transitions move towards a stem-like state, while oncogene knockdowns show an opposing trend. Finally, we organize the resource into a coherent workflow to prioritize key elements and variants, in addition to regulators. We showcase the application of this prioritization to somatic burdening, cancer differential expression and GWAS. Targeted validations of the prioritized regulators, elements and variants using siRNA knockdowns, CRISPR-based editing, and luciferase assays demonstrate the value of the ENCODE resource.

## Introduction

The 2012 ENCODE release provided comprehensive functional genomics data, such as RNA-seq, histone modification and transcription factor (TF) ChIP-seq, and DNase-seq, to annotate the noncoding regions in the human genome^1^. After the release, the cancer genomics community embraced the ENCODE data, together with other functional genomic data, to study the mutational landscape and regulatory networks in cancer^2–8^.

The current release broadens the number of cell lines and considerably expands the available tissue data. It also greatly increases the depth by adding advanced assays, such as eCLIP, RAMPAGE, ChIA-PET, Hi-C, and whole-genome STARR-seq. The ENCODE encyclopedia takes advantage of the breadth of ENCODE data to provide a “universal” annotation across hundreds of cell types. It uniformly constructs regulatory elements using assays common to all the cell types to provide an easy-to-use annotation for a wide variety of circumstances. However, a number of particular applications may require specialized annotations tailored to specific data contexts and questions (e.g., investigation of nuclear architecture or systems biology). The current ENCODE release, in fact, provides a data-rich context for a subset of cell types. Deep integration over many advanced assays allows us to connect many regulators and non-coding elements into multi-modal networks, including proximal and distal ones, such as TF and RNA-binding proteins (RBP) to gene, enhancers to gene, and TF-to-enhancer-to-gene. Here, focusing on these data-rich cell types, we developed an integrative and network-associated annotation, which may serve as a valuable resource for cancer genomics.

Cancer genomics is, in fact, one of the best applications to illustrate many key aspects of ENCODE. Unlike many other diseases, cancer is very much a disease of whole-genome alteration and dysregulation^9–12^. Moreover, cancer cells usually display aberrant behavior of key regulators, extensive epigenetic remodeling, and apparent transitions between cell states^13–17^. Finally, the systems aspect of cancer has been extensively studied, providing a need to connect linear genome annotation with pathways and networks^18–24^.

In the following sections, we first introduce the resource. We then demonstrate its utility through several applications such as evaluation of regulator activity, regulatory network rewiring, investigation of tumor-to-normal cell-state trajectories, and interpretation of expression and mutation profiles using extended genes. Synthesizing these, we propose a framework to prioritize regulators, elements, and nucleotides and then perform targeted experimental validations using different techniques.

## The ENCODEC resource

ENCODEC is a specialized *<u>ENCODE</u>* companion resource for *<u>C</u>*ancer genomics. First, using the ENCODE data, for each cancer, we try to find the best tumor-normal pairing available. To achieve this, we often constructed a “composite normal” by reconciling multiple related cell types (see suppl. sect. 1.4). Although the pairings are only approximate, many of them have been widely used in prior studies (see suppl. sect. 1.3). Then we build a derived resource. Overall, this consists of (1) comprehensive networks that allow us to see global alterations in network rewiring and regulatory hierarchy; (2) an annotated catalogue of cell types that allows us to place oncogenic changes relative to normal and stem cells; and (3) compact noncoding annotations and extended gene definitions that can potentially increase statistical power to interpret genome variation (both germline and somatic) and gene expression changes. Practically, the resource consists of a set of annotation files and computer codes available online (ENCODEC.encodeproject.org).

Figure 1 illustrates two key dimensions of the resource and the ENCODE data: breadth across cell types and depth across assays. From the depth of the ENCODE experiments in data-rich cell types, we constructed a deep, integrated annotation with two key characteristics: 1) noncoding elements are compactly defined to more precisely locate functional sites, and 2) these discontinuous regulatory regions are linked to genes to form extended-gene definitions. Extended genes are highly dynamic and may change considerably across cell types (similar in fashion to cell-type specific isoforms for conventional gene structures).

**Figure 1.**
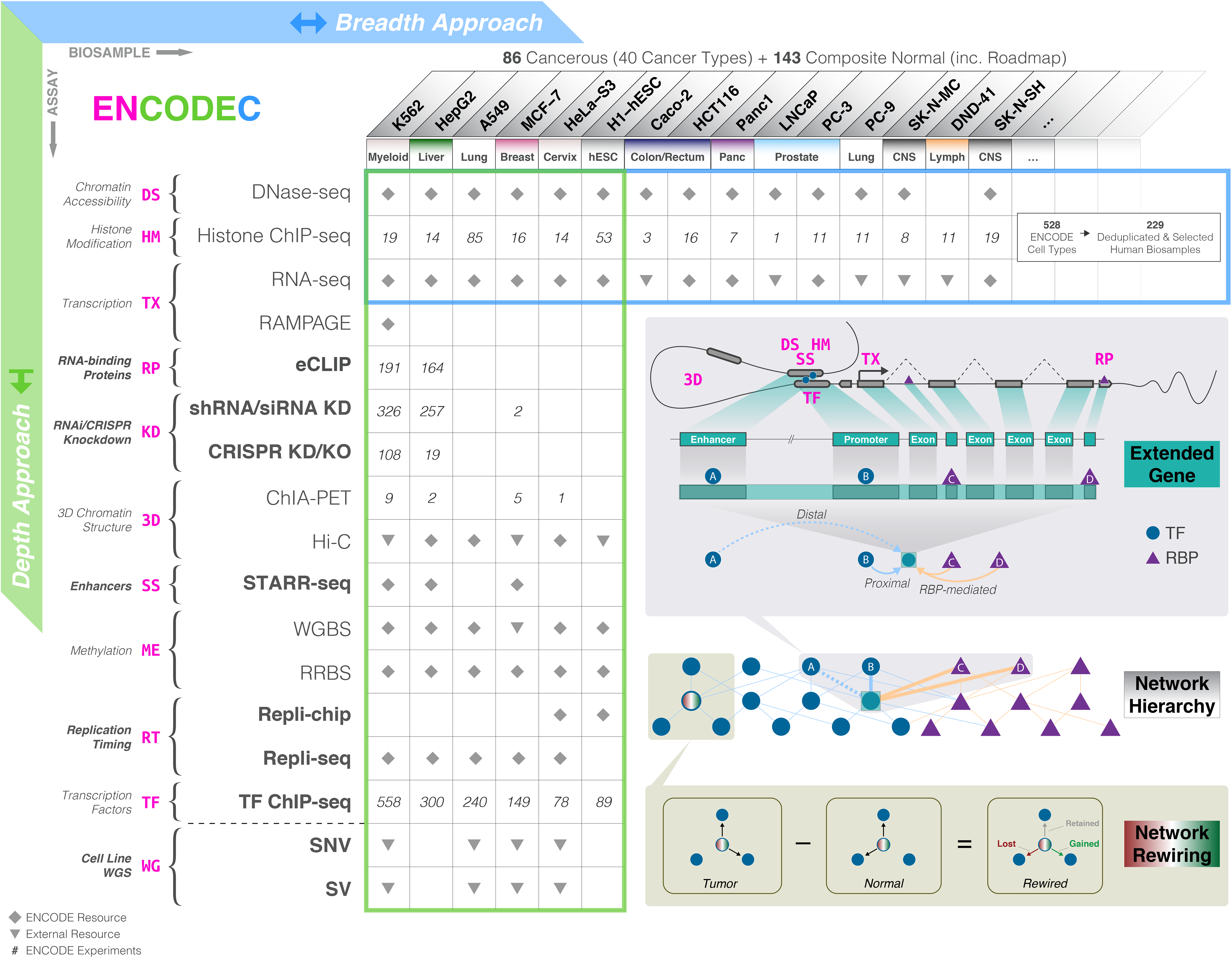
Overview of the ENCODEC resource.

Table columns list cell types and rows list assays. **Blue table boundary**: Cell types with assays in the ENCODE Encyclopedia highlight the breadth of the resource. The large number of cell types allows for comparative analyses between cell-types, as well as cell-type specific analyses. **Green table boundary**: Cell-type specific analyses based on deep annotations of cell lines. The integration of assays allows for high-resolution investigation of genomic biology. **Inset:**We use annotations from cell-type specific ENCODE assays to build extended gene definitions - coding and non-coding elements that are linked according to their interaction and associated function (top). We relate transcription factors (TFs) and RNA binding proteins (RBPs) in a joint network hierarchy that describes their regulatory potential (middle). By comparing regulatory networks in tumor and normal ENCODE samples, we develop rewiring networks that may relate to regulatory changes that occur in the context of normal-to-tumor transition (bottom).

In particular, to define distal regulatory elements (e.g. putative enhancers), we integrated up to ten histone modification ChIP-seq experiments per cell type using a support vector machine approach^25^. This procedure uses a shape-matching filter to predict enhancers based on element-associated meta-profiles of epigenetic features^26^. It has been extensively validated, giving an overall error rate of ~20% at 80% sensitivity (see suppl. sect. 2.1.2.1). Next, where possible, we intersected these regions with positives called from STARR-seq experiments (see suppl. sect. 2.1.2.2). This resulted in a substantially shorter list of distal elements than one gets with conventional approaches. Further, we restricted individual annotated elements down to a core definition enriched for functional sites by pruning based on binding motifs and using novel advanced assays such as eCLIP^27^. As a result, our annotations are short in length but have a high degree of conservation (see suppl. sect. 2.4.1).

Thus, overall, our annotation is compact in two respects: it contains fewer total elements (because the deep integration across many assays removes many potential false positives) and each individual element tends to be shorter in length yet is more enriched in functionally relevant nucleotides. In principle, both these facts benefit statistical power through decreasing multiple testing burden or more sharply defining core regions by removing nonfunctional nucleotides in each element.

We also linked together the above compact annotation elements to define extended gene structures, which may also increase power in many circumstances (see suppl. sect. 2.6). Diagramed in Fig. 1, the extended gene links the non-coding promoters and enhancers to genes. To define enhancer-gene linkages, we first used physically based linkages from Hi-C. These are accurate but often with fairly low-resolution, potentially spuriously connecting genes within the same topologically associating domain (TAD). Therefore, we pruned this with activity correlations: we correlated the chromatin marks on enhancers and gene expression on potential targets (both within the same TAD) using a machine learning approach^28^, to generate a high-confidence subset (see suppl. sect. 2.2). The extended gene annotation potentially enriches the number of functional sites being tested, thus increasing power. Second, it helps with the interpretation of noncoding elements by linking them to genes. Third, it allows us to subset non-coding annotations by the many well-known gene categories, for instance, cancer-associated and metabolic genes.

Building on the extended gene annotation, we constructed detailed networks linking regulators to genomic elements to target genes. Specifically, we built both distal and proximal networks linking TFs to genes. This was accomplished by directly inferring from ChIP-seq experiments either by TF-promoter binding or indirectly via TF-enhancer-gene interactions in each cell type (see details in suppl. sect. 2.2). We then pruned the full networks to just the strongest interactions using a signal shape algorithm that keeps the most-relevant peaks by weighting their location by the expected binding profile of each TF ^29^ (details in suppl. sect. 2.3.3). Similarly, we also defined an RBP network from eCLIP experiments. For the data-rich cell types with numerous TF ChIP-seq experiments, we further built cell-type specific regulatory networks and then compared these between matched tumor and normal cell types, enabling measurement of the change in connections during oncogenesis (i.e., network rewiring). Compared to other network definitions (e.g. via imputation based on motifs^30^), our ENCODE TF and RBP networks are based on direct experimental evidence and can capture more literature-supported regulations and correlate better with knockdown experiments (see suppl. sect. 2.4.4).

## Leveraging ENCODE networks to prioritize regulators

After constructing the multi-modal TF-RBP network, we systematically arranged it into a hierarchy (Fig. 2A-B). Here, regulators are placed at different levels such that those in the middle tend to regulate those below them and, in turn, are more regulated by regulators above them (see suppl. sect. 3.1). In the hierarchy, we find that top-layer TFs and RBPs more significantly drive differential expression (p-value<2.2e-16, one-sided Wilcoxon Test). The joint TF-RBP networks also enable investigation of cross-regulation between TFs and RBPs. Interestingly, we find that there are fewer TF-RBP interactions on the bottom level, as compared to top and middle-level ones (p-value=3.4e-16 and 1.2e-09, one-sided Wilcoxon Test, see suppl. sect. 3.7). Furthermore, we notice a well-known oncogene MYC is one of the master TFs that sits on the top-level of the hierarchy. Interestingly, MYC not only directly regulates the expression of other TFs but also targets many RBPs.

**Figure 2.**
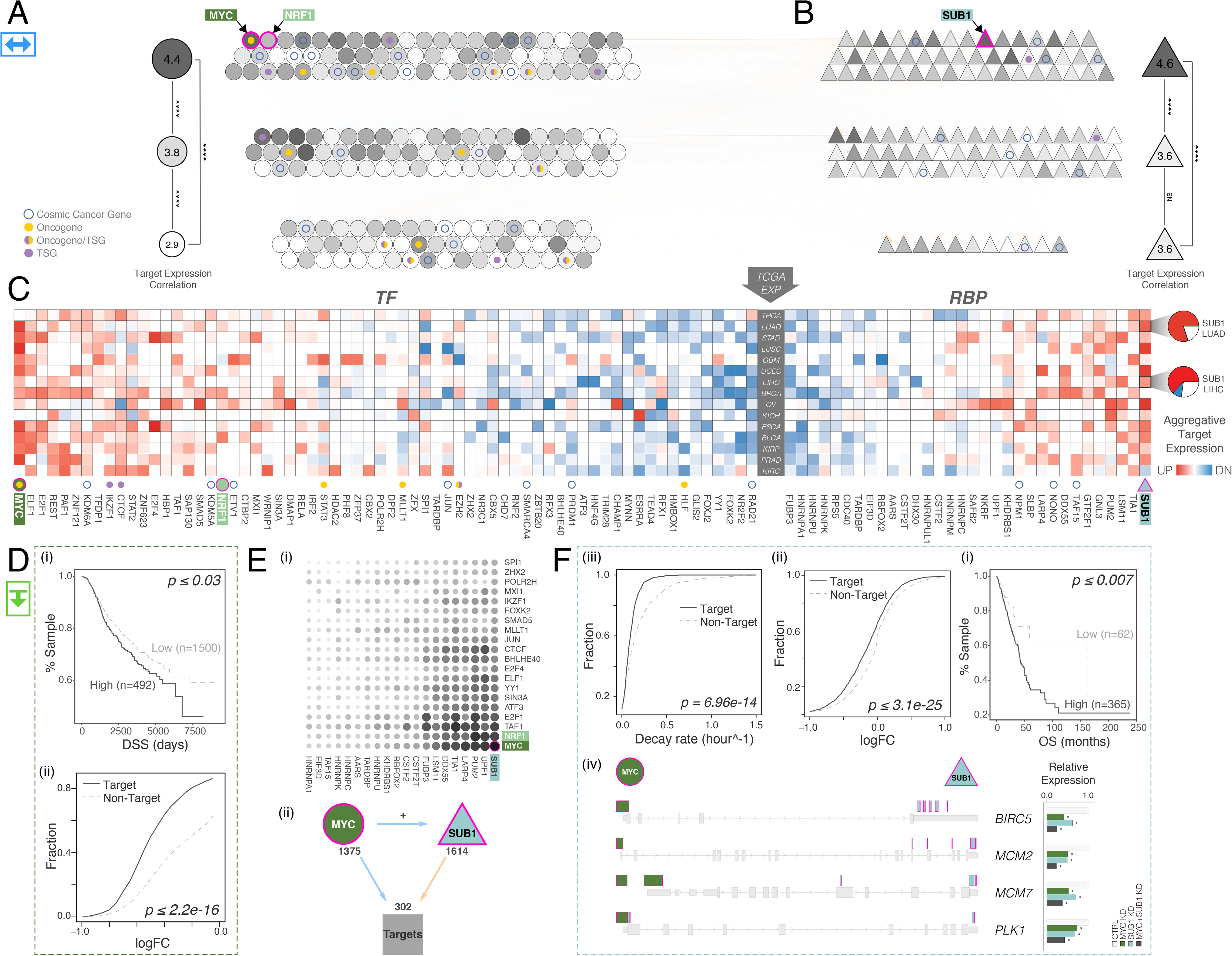
Regulatory network hierarchies.

**(A)**TFs and **(B)**RBPs are systematically organized into a hierarchy, forming a joint TF-RBP regulatory network. Higher layer elements tend to regulate lower layer elements. **(C)**The regulatory potentials of TFs/RBPs to drive tumor-to-normal expression changes are shown as a heatmap; red and blue indicate up- and down- regulation respectively. **(D)**Elevated MYC regulatory activity is associated with reduced disease-specific survival (DSS) in breast cancer (i); MYC knockdown in MCF-7 leads to significantly larger expression reduction in MYC target genes (ii). **(E)**MYC expression is more positively correlated with its target genes as compared to other TFs (top); MYC frequently forms FFLs with NRF1. These are mostly coherent FFLs and OR-gate logic predominates (bottom). (**F**) Elevated SUB1 regulation activity is associated with reduced overall survival (OS) in lung cancer (i); SUB1 knockdown in HepG2 leads to reduced target gene expression (ii); Targets of SUB1 show slower mRNA decay rate (iii); for cancer-associated target genes of MYC and SUB1, gene expression is decreased with both MYC and SUB1 knockdown (KD), compared with knockdown of either MYC or SUB1 individually, and compared to control (iv).

Our networks also enable gene-expression analyses in tumor samples. We used a regression-based approach to systematically search for the TFs and RBPs most strongly driving tumor-normal differential expression across different cancers (see suppl. sect. 3.4). For each patient, we tested the degree to which a regulator’s activity correlates with its target’s tumor-to-normal expression changes. We then calculated the percentage of patients with these relationships in each cancer type and presented the overall trends for TFs and RBPs in Fig. 2C. As expected, we find that the target genes of MYC are significantly up-regulated in numerous cancer types -- in fact, it has the most up-regulated targets of any TF -- consistent with its well-known role as a key oncogenic TF^31,32^. We further validated MYC’s regulatory effects using knockdowns (Fig. 2D). Consistent with our predictions, the expression of MYC targets is significantly reduced after MYC knockdown in MCF-7 (Fig. 2D).

We analyzed the RBP network in a manner similar to the TF network, finding regulators associated with each cancer. For example, the ENCODE eCLIP profile for the RBP SUB1 has binding peaks enriched on the 3’UTR regions of genes, and the predicted targets of SUB1 were significantly up- regulated in many cancer types (Fig. 3F, left). As an RBP, SUB1 has not been associated with cancer previously, so we sought to investigate its role. Knocking down SUB1 in HepG2 cells significantly down-regulated its targets, and the decay rate of SUB1 targets is lower than those of non-targets (Fig. 3F, right). Moreover, we find that up-regulation of SUB1 targets may lead to decreased patient survival in some cancer types.

**Figure 3.**
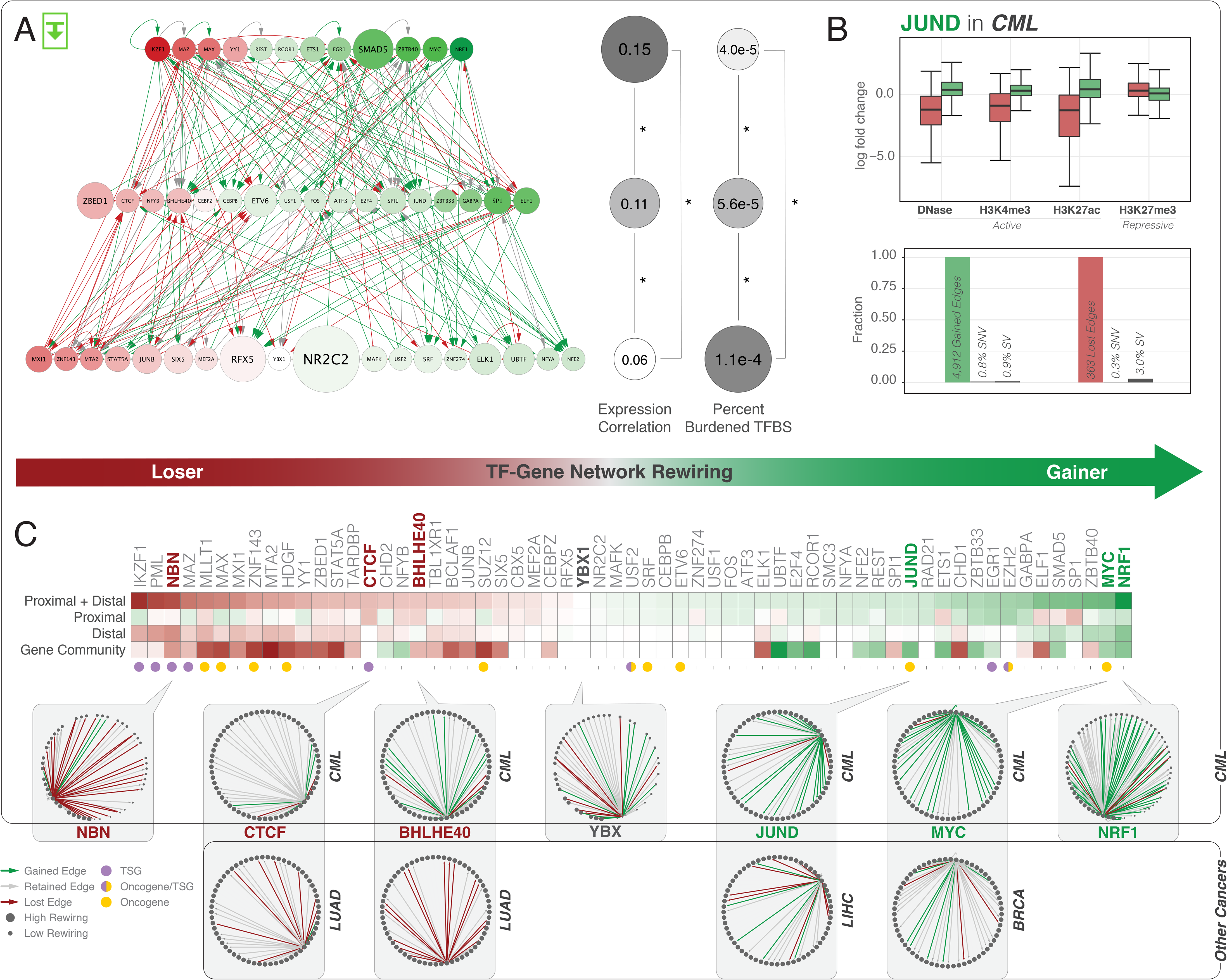
TF-Gene network rewiring.

Green and red arrows designate edge gain and loss, respectively. **(A)**Cell-type specific network using K562 and GM12878: top layer TFs significantly drive tumor-normal differential expression; bottom layer TFs are more often associated with burdened binding sites. **(B)**JUND is a top edge-gainer in CML, and its targets demonstrate increased gene expression. However, few of its binding sites are affected by SVs or SNVs. **(C)**Rewiring index in CML by direct edge counts using both proximal and distal networks (top) and by gene community analysis (bottom). Comparisons to TF-gene rewiring networks in other cancers are also shown.

We then used the regulatory network to investigate how prioritized regulators interact with each other and other genes. For TFs, we first looked at how MYC’s target genes are co-regulated by a second TF. An accounting of all the possible three-way co-regulatory relationships is shown in Fig. 2E. We find that the most common pattern is the well-characterized feed-forward loop (FFL). In this case, MYC regulates both another TF and a common target of both MYC and that TF. Many of the FFLs involve well-known MYC partners such as MAX and MXL1. However, we also discovered many involving NRF1. Upon further examination, we find that that the MYC-NRF1 FFL relationships were mostly coherent, i.e., “amplifying” in nature (see suppl. sect. 3.7). We further studied the FFLs by organizing them into logic gates, in which two TFs act as inputs and the target gene expression represents the output^33^. We find that most of these gates follow either an OR or MYC-always-dominant logic, very much in consonance with MYC’s role in driving oncogenesis.

Similarly, with respect to RBPs, we find that the top co-regulatory partner of SUB1 is, in fact, MYC. SUB1 is a direct target of MYC in many cell types (see suppl. sect. 3.7) and also forms many FFLs with MYC in the regulatory network. We hypothesized that MYC binds to the promoter regions of key oncogenes to initiate their transcription, whereas SUB1 binds to their 3’ UTRs to stabilize their RNA transcripts. Such collaboration between MYC and SUB1 potentially could result in the overexpression of several key oncogenes (see suppl. sect. 3.7). To validate this hypothesis, we knocked down MYC and SUB1 in HepG2 and used qPCR to quantify changes in gene expression. As expected, the expression of oncogenes (such as MCM2, MCM7, BIRC5, and PLK1) is significantly reduced (Fig. 2F and see suppl. sect. 3.5).

## Measuring network rewiring

In addition to the TF regulatory activity change through expression analysis above, we also directly measured the fractional number of regulatory edge changes for “tumor-normal pairs”, to study how TF targets change in oncogenesis. We call this the “rewiring index” and ranked TFs according to it (Fig. 3C). In leukemia, well-known oncogenes (such as MYC and NRF1) were among the top edge gainers, while the well-known tumor suppressor IKZF1 is the most significant edge loser (Fig. 3C). Mutations in IKZF1, in fact, serve as a hallmark of various forms of high-risk leukemia^34,35^. We observed a similar rewiring trend using distal, proximal, and combined networks (Fig. 3C). This trend was also consistent across a number of cancers: in particular, highly rewired TFs such as BHLHE40, JUND, and MYC behaved similarly in lung, liver, and breast cancers (Fig. 3C).

In addition to direct TF-to-gene connections, we also measured rewiring using a gene-community model. Here, the targets within the regulatory network were characterized in terms of self-consistent modules of related genes (so-called “gene communities”). Instead of directly measuring the changes in a TF’s targets between tumor and normal cells, we determined the changes in regulated gene communities (via a mixed-membership model, see suppl. sect. 4.3.3). Similar patterns to direct rewiring were observed (Fig. 3C).

Overall, we find that the majority of rewiring events were associated with notable gene-expression and chromatin-status changes, but not necessarily with direct variant-induced motif loss or gain events (Fig. 3B). For example, JUND is a top edge gainer in K562. Most of its gained targets in tumor cells demonstrate higher levels of gene expression, stronger active and weaker repressive histone-modification signals, yet few of its binding sites are mutated, either by SNVs or SVs. This is consistent with previous work^36^, and with a few notable exceptions, we find a similar trend for the rewiring events associated with JUND in liver cancer and, largely, for other factors in a variety of cancers (see suppl. sect. 4.4).

We also organized the cell-type specific networks into hierarchies, as shown in Fig. 3A (similar to the “universal,” cross-cell-type hierarchies described earlier in Fig. 2A-B). We find that the strongest edge gainers and losers, driving the rewiring of the regulatory network, sit at the top level of these hierarchies in blood cancer. In addition, we find the TFs more associated with driving cancer gene expression changes also tend to be at the top. MYC is a most prominent example of both a highly-rewired TF and one driving expression. In contrast, the more mutationally affected TFs sit at the bottom of the hierarchy. To some degree, this is consistent with our results in Fig. 3B showing that binding site mutations do not drive the regulatory change.

## Placing cancer cells in the context of many ENCODE cell types

ENCODE data provides an additional way of studying the oncogenic transformation beyond network rewiring: via placing various cancer cells in a context of many cell types (in “cell space”). This is possible because of the wide variety of cell types profiled in the new ENCODE release, which includes many stem cells, especially the data-rich H1 cell line. We are particularly interested in comparisons to stem cells since a decades-old paradigm has held that at least a subpopulation of tumor cells can self-renew, differentiate, and regenerate in a manner similar to stem cells^37–42^. For such comparison, we first projected the RNA-seq data from 299 ENCODE cell types into a low-dimensional space (using the procedure described in Li et al^43^, see suppl. sect. 5.1). We find that various types of stem cells form a tight cluster (Fig. 4). Moreover, there is a trend where the trajectory from normal to tumor cells involves moving toward stem cells, along a single “stem-like component.” This is true for a variety of different cancers. This observation is consistent with previous efforts using expression and methylation analysis^44^. Notably, we observed a consistent (or even stronger) pattern from proximal and distal chromatin data, which can be viewed as the underlying cause of the observed gene expression changes.

**Figure 4.**
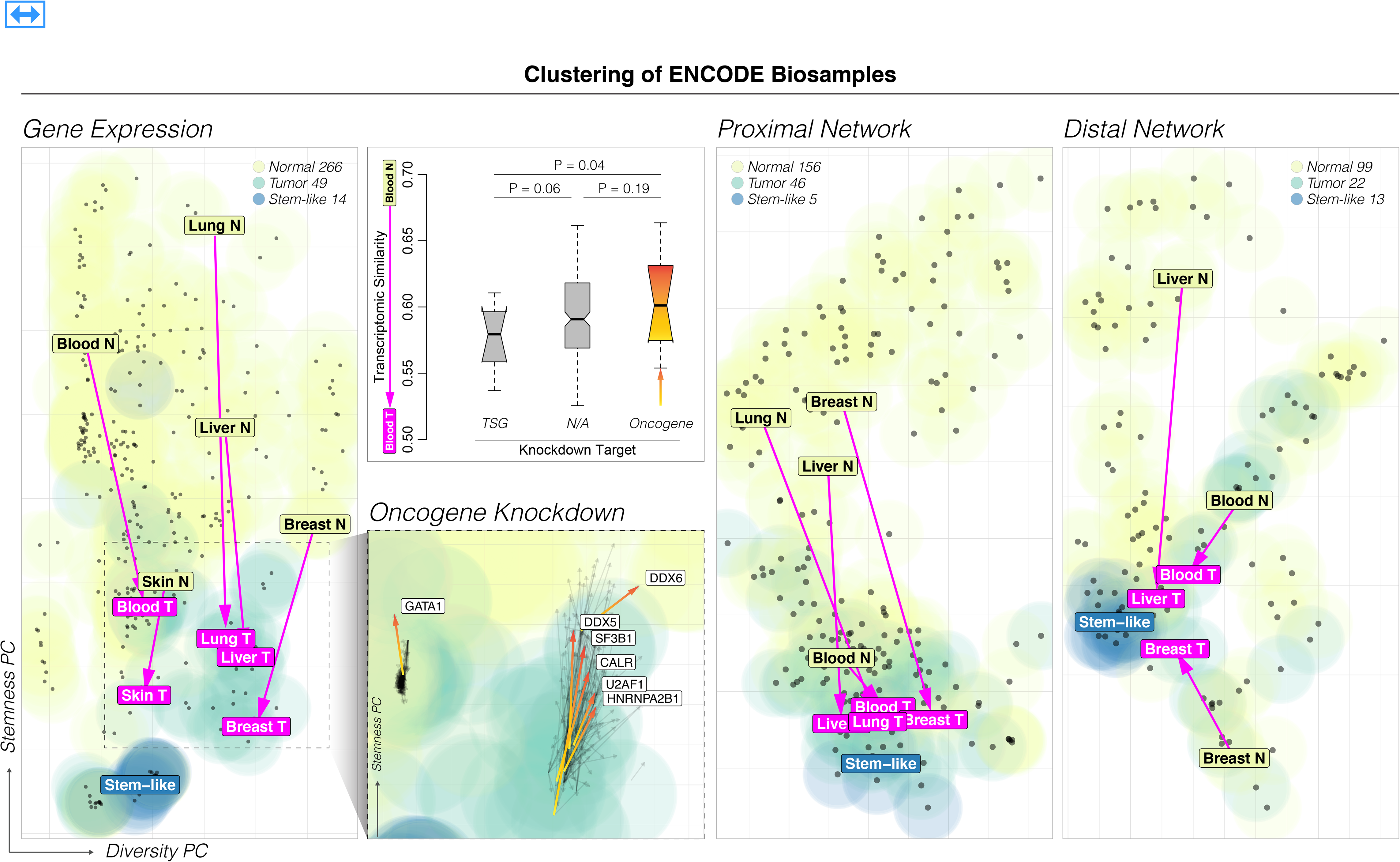
Oncogenic transformation and cell state.

We project the expression profiles (left, poly-A long RNA-seq), proximal network (second from right, CTCF ChIP-seq), and distal network (right, candidate cis-regulatory elements) of the ENCODE cell types to a lower dimension space. Stem-like cell types formed a cluster, suggesting stem-like cell types have a distinct profile from normal and cancerous cell types. Further, we find that cancerous cell types tend to locate closer to stem-like clusters. Oncogene knockdown in K562 led to more transcriptomic similarity to a normal cell-type, and tumor suppressor gene (TSG) knockdown led to greater similarity to a tumor cell-type (second from left, top, in comparison to GM12878). In general, we find that oncogene knockdown leads to a slight reversion towards normal state along the stem-like component (second from left, bottom).

It is well-known that dysregulation of oncogene TFs is a hallmark of tumor progression^11,45-48^. Key genes, such as MYC, initiate overexpression of other oncogenes in tumor cells^32,49^. We can use the cell-space diagram to see the degree to which these TFs contribute to the state of cell differentiation: in particular, we measured the perturbations induced by oncogenic TFs through expression comparisons before and after TF knockdowns. Interestingly, the expression profiles usually reverted slightly back towards normal state upon oncogene knockdown, along the stem-like component. One can see this difference more precisely and test it statistically if one restricts just to the single transition between GM12878 and K562 (Fig. 4).

## The extended gene representation

After identifying key regulators, we next aimed to prioritize their associated genomic elements. To do this, we combined the extended gene annotation with expression and mutation data from patients. We show three examples where this is useful.

First, our extended gene definitions can be used for associating differential expression with mutational status. For example, we combined the mutation and expression profiles from large cohorts, such as those in TCGA, and found that mutation status in extended genes can better explain the tumor expression than other annotations, such as just canonical coding sequences (CDS). That is, one can much better predict tumor-normal differential expression from mutations in the extended gene as compared to just in CDS or in individual promoters or enhancers (see suppl. sect. 6.1). One example of the explanatory potential of the extended gene is seen for the splicing factor SRSF2, which has been shown to affect liver cancer progression and for which differential expression in HepG2 can be well predicted using mutations in the extended gene (Fig. 5A, p-value=0.002, one-sided Wilcoxon test).

**Figure 5.**
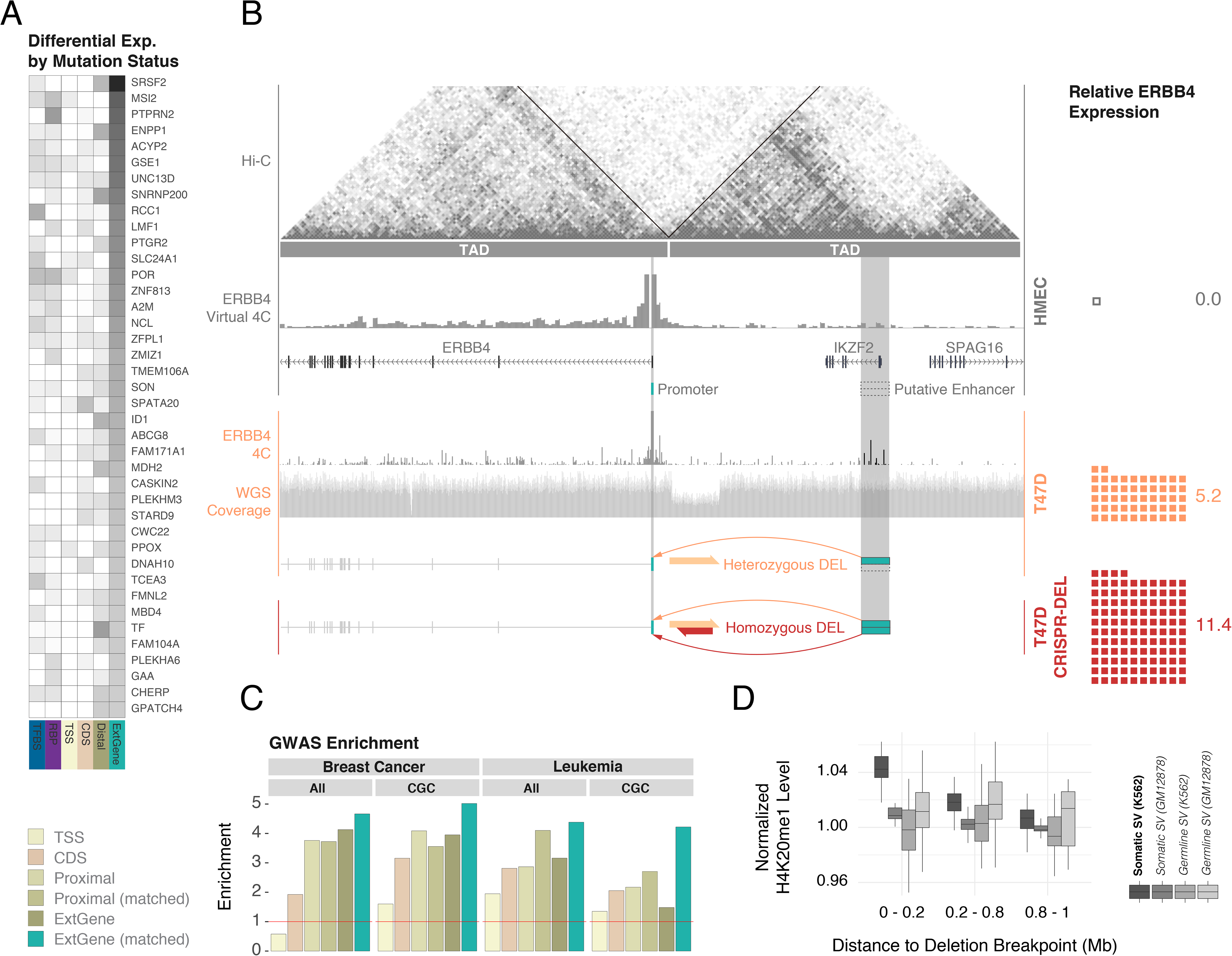
Extended genes and mutation burden analysis.

**(A)**Mutation status in extended genes can explain expression differences for a larger number of genes than other annotations, such as annotations of coding sequences (CDS). **(B)**A 130kbp deletion in the breast cancer cell line T47D potentially links a distal enhancer to the promoter of ERBB4, leading to its activation. This change does not affect coding sequences, highlighting the value of an extended gene annotation. **(C)**Cancer-associated GWAS SNVs display greater enrichment with the inclusion of proximal and distal annotations in extended gene definitions. **(D)**Somatic structural variant breakpoints in K562 tend to be associated with the activating histone mark H4K20me1, but not in GM12878.

The second example is cancer genome-wide association study (GWAS) variant enrichment. That is, the enrichment of cancer-associated GWAS germline SNPs in particular genome regions. The enrichment significantly increases in going from CDS to extended genes for both breast cancer and leukemia (Fig. 5C). This trend is much more pronounced when the newly added non-coding annotations are from matched cell types. One may further subset the genes according to different subcategories associated with cancer and identify enrichment. For instance, we observed a significant enrichment in genes from the Cancer Gene Consensus (CGC) in breast cancer based on the extended gene annotation. This sub-setting by well-known gene categories is not possible using conventional non-coding annotations.

One can get a physical sense of the importance of the extended gene by looking at a situation where a genomic variant rearranges the extended gene structure without affecting the coding regions. We find such an example in the breast cancer cell line T47D, where a 130kbp heterozygous deletion links a distal enhancer to the ERBB4 promoter and results in the activation of this well-known oncogene^64,65^ (Fig. 5B). The enhancer is not connected to ERBB4 in normal breast tissue; however, in T47D, the deletion, located around 45kbp downstream from the ERBB4 promoter, merges two Hi-C TADs in an allele-specific way. We tested this through CRISPR editing, by excising an 86bp sequence within the wild-type allele of the heterozygous deletion containing the CTCF binding sites at the boundary of the two TADs. This CRISPR excision confirmed the elevated ERBB4 expression (see suppl. sect. 6.4).

Another perspective on the effect of SVs changing chromatin structure is provided from broadly surveying SVs in a number of the data-rich ENCODE cells types. (Note, ENCODE provides SV call sets based on integration of assays including Hi-C for a number of these cell lines, see suppl. sect. 6.5.3). In particular, in Fig. 5D, we surveyed regions around somatic SV breakpoints in K562. We find that the activating histone mark H4K20me1 occurs preferentially around these breakpoints. This enrichment was not observed using GM12878 histone mark data at these exact same locations. We further examined the GM12878 H4K20me1 levels proximal to germline breakpoints (for common variants as determined from the 1000 Genomes Project^66^) and also find no enrichment (see suppl. sect. 6.5). One potential implication is that the somatic SVs in tumor cells may be associated with creating active regions of chromatin.

## Step-wise prioritization framework

Collectively, as described in Fig. 6, ENCODEC enables a step-wise prioritization that allows us to pinpoint key regulators, noncoding elements, and variants associated with oncogenesis. Specifically, we first highlighted regulators that are either greatly rewired, located in hubs, sit at the top of the hierarchy, or significantly drive expression changes in cancer. We then prioritize functional elements associated with these regulators that are either highly burdened by mutations, undergo large chromatin changes, or change in extended gene linkages. Finally, on a nucleotide level, we prioritize SNVs by estimating their ability to disrupt or introduce specific binding sites and assessing to what degree they lie in a prioritized element.

**Figure 6.**
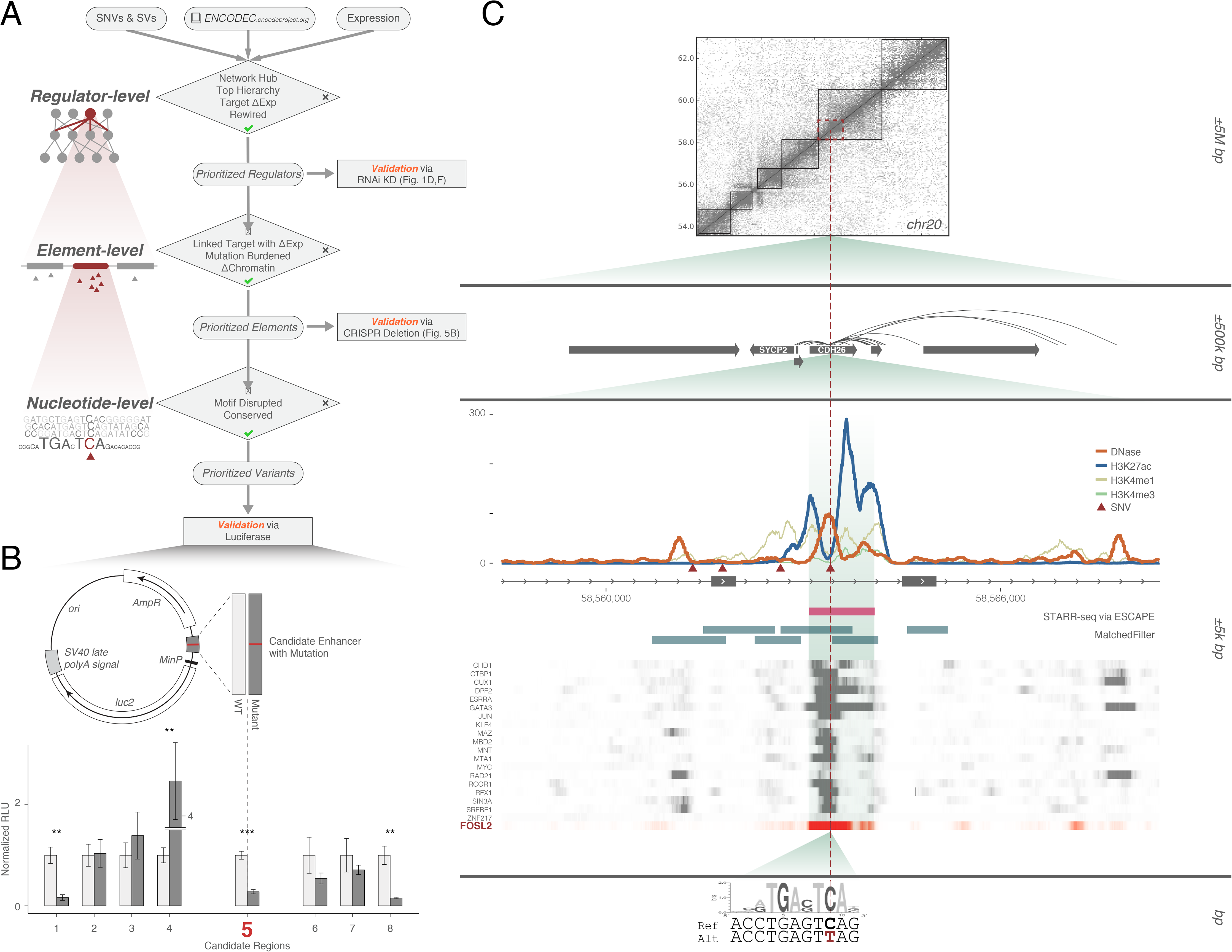
Variant prioritization and validation.

**(A)**A stepwise prioritization scheme for genomic regulators, elements, and variants, using the ENCODEC resources. At each step of prioritization, we indicate criteria for prioritization, as well as the applicable validation assay. **(B)**Small-scale validation of prioritized variants using a luciferase reporter assay. Candidate region 5 showed the most significant degree of differential expression and was selected for follow-up analysis. **(C)**Multiscale integrative analysis of candidate region 5 with assorted functional genomics data. The affected region is observed in the context of large-scale Hi-C linkages (top), as well as element-level signal tracks of histone modification marks and DNase hypersensitivity together with various TF binding events (middle), and nucleotide level disruption of the FOSL2 motif (bottom).

We instantiated our prioritization workflow in a few select cancers and experimentally validated the results. In particular, as described above, we subjected some key regulators, such as MYC and SUB1, to knockdown experiments (Fig. 2D and Fig. 2F) and we measured the effect of SVs on element linkages via CRISPR engineered deletions (Fig. 5B). Finally, we selected key SNVs based on their disruption of enhancers with a strong influence on gene expression. These SNVs were prioritized based on element-level mutation recurrence in breast-cancer cohorts, as well as motif disruption scores. Of the eight motif-disrupting SNVs that we tested, six exhibited consistent up- or down-regulation relative to the wild-type in multiple biological replicates (see suppl. sect. 7.2 and 7.3).

One particularly interesting example occurs in an intronic region of CDH26 in chromosome 20 (Fig. 6C). The signal shapes for both histone modification and chromatin accessibility (DNase-seq) data indicate its active regulatory role as an enhancer in MCF-7. This was further confirmed by STARR-seq (Fig. 6C). Hi-C and ChIA-PET linkages indicated that the region is within a TAD and validated a regulatory connection to the cancer-associated gene SYCP2^67^. We further observed strong binding of many TFs in this region in MCF-7. Motif analysis predicts that a common mutation in breast cancer affects this region, and significantly disrupts the local binding affinity of several TFs, such as FOSL2 (Fig. 6C). Luciferase assays demonstrated that this mutation introduces a 3.6-fold reduction in expression relative to the wild-type, indicating a strong repressive effect on enhancer functionality.

## Discussion

In this paper, we describe a customized ENCODE annotation: a companion resource providing an integrative network annotation including extended gene. Cancer genomics is an ideal application to highlight the value of the resource, and we show how it can help describe oncogenic transformations in terms of cell-space trajectories and network rewiring. We also use the specialized annotation to prioritize key regulators, element, and variants.

There remain several caveats associated with our resource. First and most obviously, proper somatic variant annotation and, especially driver discovery, is a multiple-step process that requires coordinated, large-scale effort. Extensive follow-up validations are required, in addition to the careful calibration required for statistical identification of mutation recurrence and the many biases in sequencing (e.g. taking into account the elevated mutation rate associated with TF binding sites^2,6^, sequence coverage and mutational signatures^50,57^). While we hope that ENCODE data and annotation can be useful in this context, they are not sufficient. Second, our resource associates cancer types with ENCODE cell lines and then secondarily pairs them with a composite normal. Both types of pairings are, by nature, approximate. Tumor cells from a given patient show distinct molecular, morphological, and genetic profiles^68–71^. Moreover, linking cancer to one specific cell-type may not even fully capture the heterogeneity seen in actual tumors^72^. In the future, technological advances, such as single-cell sequencing, may allow cell-type or tissue-type comparisons at a higher resolution^73–77^. Nevertheless, we feel that our annotation and networks currently provide the best available view of the regulatory changes in oncogenesis.

Finally, we argue here that, somewhat counter-intuitively, a comprehensive non-coding annotation that, in the extreme, attempts to assign functional impact to every base in the genome may not always be best suited to specific disease-oriented studies. Rather, the most useful annotation often has several characteristics. First, it is useful to be as compact as possible, both in terms of the extent of individual annotation blocks and in the number of elements. Second, since the currently discovered high impact variants tend to be tightly associated with genes, an optimum non-coding annotation is best “invisible,” folding itself into gene annotation for better variant interpretation. Third, the network aspect is often needed to allow larger-scale systems perspective. This is particularly valuable for appreciating the overall cellular dysregulation in cancer. With the depth and breadth of the ENCODE assays across thousands of cell types, we endeavored here to provide such a customized annotation resource for cancer and demonstrated its value through several showcase applications. We anticipate that the rapid accumulation of functional genomic data will make possible further, potentially even more specialized, annotation resources for future disease studies.

## Notes

http://encodec.encodeproject.org

## Reference

1 Consortium, E. P. An integrated encyclopedia of DNA elements in the human genome. Nature 489, 57-74, doi:10.1038/nature11247 (2012).,

2 Frigola, J. et al. Reduced mutation rate in exons due to differential mismatch repair. Nat Genet 49, 1684-1692, doi:10.1038/ng.3991 (2017).

3 Martincorena, I. et al. Universal Patterns of Selection in Cancer and Somatic Tissues. Cell 171, 1029-1041 e1021, doi:10.1016/j.cell.2017.09.042 (2017).

4 Imielinski, M., Guo, G. & Meyerson, M. Insertions and Deletions Target Lineage-Defining Genes in Human Cancers. Cell 168, 460-472 e414, doi:10.1016/j.cell.2016.12.025 (2017).

5 Nik-Zainal, S. et al. Landscape of somatic mutations in 560 breast cancer whole-genome sequences. Nature 534, 47-54, doi:10.1038/nature17676 (2016).

6 Sabarinathan, R., Mularoni, L., Deu-Pons, J., Gonzalez-Perez, A. & Lopez-Bigas, N. Nucleotide excision repair is impaired by binding of transcription factors to DNA. Nature 532, 264-267, doi:10.1038/nature17661 (2016).

7 Supek, F. & Lehner, B. Differential DNA mismatch repair underlies mutation rate variation across the human genome. Nature 521, 81-84, doi:10.1038/nature14173 (2015).

8 Polak, P. et al. Cell-of-origin chromatin organization shapes the mutational landscape of cancer. Nature 518, 360-364, doi:10.1038/nature14221 (2015).

9 Ntziachristos, P., Abdel-Wahab, O. & Aifantis, I. Emerging concepts of epigenetic dysregulation in hematological malignancies. Nat Immunol 17, 1016-1024, doi:10.1038/ni.3517 (2016).

10 Liu, F., Wang, L., Perna, F. & Nimer, S. D. Beyond transcription factors: how oncogenic signalling reshapes the epigenetic landscape. Nat Rev Cancer 16, 359-372, doi:10.1038/nrc.2016.41 (2016).

11 Gonda, T. J. & Ramsay, R. G. Directly targeting transcriptional dysregulation in cancer. Nat Rev Cancer 15, 686-694, doi:10.1038/nrc4018 (2015).

12 Timp, W. & Feinberg, A. P. Cancer as a dysregulated epigenome allowing cellular growth advantage at the expense of the host. Nat Rev Cancer 13, 497-510, doi:10.1038/nrc3486 (2013).

13 Baylin, S. B. & Jones, P. A. A decade of exploring the cancer epigenome - biological and translational implications. Nat Rev Cancer 11, 726-734, doi:10.1038/nrc3130 (2011).

14 Polyak, K. & Weinberg, R. A. Transitions between epithelial and mesenchymal states: acquisition of malignant and stem cell traits. Nat Rev Cancer 9, 265-273, doi:10.1038/nrc2620 (2009).

15 Yu, H. & Jove, R. The STATs of cancer--new molecular targets come of age. Nat Rev Cancer 4, 97-105, doi:10.1038/nrc1275 (2004).

16 Darnell, J. E. Jr., Transcription factors as targets for cancer therapy. Nat Rev Cancer 2, 740-749, doi:10.1038/nrc906 (2002).

17 Jones, P. A. & Baylin, S. B. The fundamental role of epigenetic events in cancer. Nat Rev Genet 3, 415-428, doi:10.1038/nrg816 (2002).

18 Sanchez-Vega, F. et al. Oncogenic Signaling Pathways in The Cancer Genome Atlas. Cell 173, 321-337 e310, doi:10.1016/j.cell.2018.03.035 (2018).

19 Garraway, L. A. & Lander, E. S. Lessons from the cancer genome. Cell 153, 17-37, doi:10.1016/j.cell.2013.03.002 (2013).

20 Vogelstein, B. et al. Cancer genome landscapes. Science 339, 1546-1558, doi:10.1126/science.1235122 (2013).

21 Horn, H. et al. NetSig: network-based discovery from cancer genomes. Nat Methods 15, 61-66, doi:10.1038/nmeth.4514 (2018).

22 Creixell, P. et al. Pathway and network analysis of cancer genomes. Nat Methods 12, 615-621, doi:10.1038/nmeth.3440 (2015).

23 Leiserson, M. D. et al. Pan-cancer network analysis identifies combinations of rare somatic mutations across pathways and protein complexes. Nat Genet 47, 106-114, doi:10.1038/ng.3168 (2015).

24 Hofree, M., Shen, J. P., Carter, H., Gross, A. & Ideker, T. Network-based stratification of tumor mutations. Nat Methods 10, 1108-1115, doi:10.1038/nmeth.2651 (2013).

25 Sethi, A. et al. A cross-organism framework for supervised enhancer prediction with epigenetic pattern recognition and targeted validation. bioRxiv, doi:10.1101/385237 (2018).

26 Kundaje, A. et al. Ubiquitous heterogeneity and asymmetry of the chromatin environment at regulatory elements. Genome Res 22, 1735-1747, doi:10.1101/gr.136366.111 (2012).

27 Van Nostrand, E. L. et al. Robust transcriptome-wide discovery of RNA-binding protein binding sites with enhanced CLIP (eCLIP). Nat Methods 13, 508-514, doi:10.1038/nmeth.3810 (2016).

28 Cao, Q. et al. Reconstruction of enhancer-target networks in 935 samples of human primary cells, tissues and cell lines. Nat Genet 49, 1428-1436, doi:10.1038/ng.3950 (2017).

29 Cheng, C., Min, R. & Gerstein, M. TIP: a probabilistic method for identifying transcription factor target genes from ChIP-seq binding profiles. Bioinformatics 27, 3221-3227, doi:10.1093/bioinformatics/btr552 (2011).

30 Neph, S. et al. Circuitry and dynamics of human transcription factor regulatory networks. Cell 150, 1274-1286, doi:10.1016/j.cell.2012.04.040 (2012).

31 McKeown, M. R. & Bradner, J. E. Therapeutic strategies to inhibit MYC. Cold Spring Harb Perspect Med 4, doi:10.1101/cshperspect.a014266 (2014).

32 Dang, C. V. MYC on the path to cancer. Cell 149, 22-35, doi:10.1016/j.cell.2012.03.003 (2012).

33 Wang, D. et al. Loregic: a method to characterize the cooperative logic of regulatory factors. PLoS Comput Biol 11, e1004132, doi:10.1371/journal.pcbi.1004132 (2015).

34 Boer, J. M. et al. Prognostic value of rare IKZF1 deletion in childhood B-cell precursor acute lymphoblastic leukemia: an international collaborative study. Leukemia 30, 32-38, doi:10.1038/leu.2015.199 (2016).

35 de Rooij, J. D. et al. Recurrent deletions of IKZF1 in pediatric acute myeloid leukemia. Haematologica 100, 1151-1159, doi:10.3324/haematol.2015.124321 (2015).

36 Farh, K. K. et al. Genetic and epigenetic fine mapping of causal autoimmune disease variants. Nature 518, 337-343, doi:10.1038/nature13835 (2015).

37 O’Connor, M. L. et al. Cancer stem cells: A contentious hypothesis now moving forward. Cancer Lett 344, 180-187, doi:10.1016/j.canlet.2013.11.012 (2014).

38 Ge, Y. et al. Stem Cell Lineage Infidelity Drives Wound Repair and Cancer. Cell 169, 636-650 e614, doi:10.1016/j.cell.2017.03.042 (2017).

39 Fabregat, I., Malfettone, A. & Soukupova, J. New Insights into the Crossroads between EMT and Stemness in the Context of Cancer. J Clin Med 5, doi:10.3390/jcm5030037 (2016).

40 Friedmann-Morvinski, D. & Verma, I. M. Dedifferentiation and reprogramming: origins of cancer stem cells. EMBO Rep 15, 244-253, doi:10.1002/embr.201338254 (2014).

41 Eppert, K. et al. Stem cell gene expression programs influence clinical outcome in human leukemia. Nat Med 17, 1086-1093, doi:10.1038/nm.2415 (2011).

42 Gentles, A. J., Plevritis, S. K., Majeti, R. & Alizadeh, A. A. Association of a leukemic stem cell gene expression signature with clinical outcomes in acute myeloid leukemia. JAMA 304, 2706-2715, doi:10.1001/jama.2010.1862 (2010).

43 Li, H. et al. Reference component analysis of single-cell transcriptomes elucidates cellular heterogeneity in human colorectal tumors. Nat Genet 49, 708-718, doi:10.1038/ng.3818 (2017).

44 Malta, T. M. et al. Machine Learning Identifies Stemness Features Associated with Oncogenic Dedifferentiation. Cell 173, 338-354 e315, doi:10.1016/j.cell.2018.03.034 (2018).

45 Hanahan, D. & Weinberg, R. A. Hallmarks of cancer: the next generation. Cell 144, 646-674, doi:10.1016/j.cell.2011.02.013 (2011).

46 Vicente-Duenas, C., Romero-Camarero, I., Cobaleda, C. & Sanchez-Garcia, I. Function of oncogenes in cancer development: a changing paradigm. EMBO J 32, 1502-1513, doi:10.1038/emboj.2013.97 (2013).

47 Santhekadur, P. K. et al. The transcription factor LSF: a novel oncogene for hepatocellular carcinoma. Am J Cancer Res 2, 269-285 (2012).

48 Perkins, N. D. The diverse and complex roles of NF-kappaB subunits in cancer. Nat Rev Cancer 12, 121-132, doi:10.1038/nrc3204 (2012).

49 Lin, C. Y. et al. Transcriptional amplification in tumor cells with elevated c-Myc. Cell 151, 56-67, doi:10.1016/j.cell.2012.08.026 (2012).

50 Bailey, M. H. et al. Comprehensive Characterization of Cancer Driver Genes and Mutations. Cell 173, 371-385 e318, doi:10.1016/j.cell.2018.02.060 (2018).

51 Kandoth, C. et al. Mutational landscape and significance across 12 major cancer types. Nature 502, 333-339, doi:10.1038/nature12634 (2013).

52 Lawrence, M. S. et al. Mutational heterogeneity in cancer and the search for new cancer-associated genes. Nature 499, 214-218, doi:10.1038/nature12213 (2013).

53 Lochovsky, L., Zhang, J., Fu, Y., Khurana, E. & Gerstein, M. LARVA: an integrative framework for large-scale analysis of recurrent variants in noncoding annotations. Nucleic Acids Res 43, 8123-8134, doi:10.1093/nar/gkv803 (2015).

54 Melton, C., Reuter, J. A., Spacek, D. V. & Snyder, M. Recurrent somatic mutations in regulatory regions of human cancer genomes. Nat Genet 47, 710-716, doi:10.1038/ng.3332 (2015).

55 Wang, K. et al. Whole-genome sequencing and comprehensive molecular profiling identify new driver mutations in gastric cancer. Nat Genet 46, 573-582, doi:10.1038/ng.2983 (2014).

56 Weinhold, N., Jacobsen, A., Schultz, N., Sander, C. & Lee, W. Genome-wide analysis of noncoding regulatory mutations in cancer. Nat Genet 46, 1160-1165, doi:10.1038/ng.3101 (2014).

57 Rheinbay, E. et al. Discovery and characterization of coding and non-coding driver mutations in more than 2,500 whole cancer genomes. bioRxiv, doi:10.1101/237313 (2017).

58 Morganella, S. et al. The topography of mutational processes in breast cancer genomes. Nat Commun 7, 11383, doi:10.1038/ncomms11383 (2016).

59 Schuster-Bockler, B. & Lehner, B. Chromatin organization is a major influence on regional mutation rates in human cancer cells. Nature 488, 504-507, doi:10.1038/nature11273 (2012).

60 Tomkova, M., Tomek, J., Kriaucionis, S. & Schuster-Boeckler, B. Widespread impact of DNA replication on mutational mechanisms in cancer. bioRxiv, doi:10.1101/111302 (2017).

61 Cattoretti, G. et al. Deregulated BCL6 expression recapitulates the pathogenesis of human diffuse large B cell lymphomas in mice. Cancer Cell 7, 445-455, doi:10.1016/j.ccr.2005.03.037 (2005).

62 Niu, H. The proto-oncogene BCL-6 in normal and malignant B cell development. Hematol Oncol 20, 155-166, doi:10.1002/hon.689 (2002).

63 Shvarts, A. et al. A senescence rescue screen identifies BCL6 as an inhibitor of anti-proliferative p19(ARF)-p53 signaling. Genes Dev 16, 681-686, doi:10.1101/gad.929302 (2002).

64 Yu, T. et al. MicroRNA-193a-3p and -5p suppress the metastasis of human non-small-cell lung cancer by downregulating the ERBB4/PIK3R3/mTOR/S6K2 signaling pathway. Oncogene 34, 413-423, doi:10.1038/onc.2013.574 (2015).

65 Sundvall, M. et al. Role of ErbB4 in breast cancer. J Mammary Gland Biol Neoplasia 13, 259-268, doi:10.1007/s10911-008-9079-3 (2008).

66 Genomes Project, C. et al. A global reference for human genetic variation. Nature 526, 68-74, doi:10.1038/nature15393 (2015).

67 Masterson, L. et al. Deregulation of SYCP2 predicts early stage human papillomavirus-positive oropharyngeal carcinoma: A prospective whole transcriptome analysis. Cancer Sci 106, 1568-1575, doi:10.1111/cas.12809 (2015).

68 Patel, A. P. et al. Single-cell RNA-seq highlights intratumoral heterogeneity in primary glioblastoma. Science 344, 1396-1401, doi:10.1126/science.1254257 (2014).

69 Bedard, P. L., Hansen, A. R., Ratain, M. J. & Siu, L. L. Tumour heterogeneity in the clinic. Nature 501, 355-364, doi:10.1038/nature12627 (2013).

70 Meacham, C. E. & Morrison, S. J. Tumour heterogeneity and cancer cell plasticity. Nature 501, 328-337, doi:10.1038/nature12624 (2013).

71 Gerlinger, M. et al. Intratumor heterogeneity and branched evolution revealed by multiregion sequencing. N Engl J Med 366, 883-892, doi:10.1056/NEJMoa1113205 (2012).

72 Visvader, J. E. Cells of origin in cancer. Nature 469, 314-322, doi:10.1038/nature09781 (2011).

73 Tirosh, I. et al. Dissecting the multicellular ecosystem of metastatic melanoma by single-cell RNA-seq. Science 352, 189-196, doi:10.1126/science.aad0501 (2016).

74 Gawad, C., Koh, W. & Quake, S. R. Single-cell genome sequencing: current state of the science. Nat Rev Genet 17, 175-188, doi:10.1038/nrg.2015.16 (2016).

75 Rotem, A. et al. Single-cell ChIP-seq reveals cell subpopulations defined by chromatin state. Nat Biotechnol 33, 1165-1172, doi:10.1038/nbt.3383 (2015).

76 Eirew, P. et al. Dynamics of genomic clones in breast cancer patient xenografts at single-cell resolution. Nature 518, 422-426, doi:10.1038/nature13952 (2015).

77 Wang, Y. et al. Clonal evolution in breast cancer revealed by single nucleus genome sequencing. Nature 512, 155-160, doi:10.1038/nature13600 (2014).

