## Supplementary Information for "An integrative ENCODE resource for cancer genomics"

**SUPPLEMENTARY INFORMATION TO**

### Integrative noncoding annotation and regulatory network for cancer genomics

Figure S 5-6. projection of ENCODE knockdown data onto the RCA space. Start of an arrow represents the placement of the cell before knockdown (control experiment) and the tip of arrow represents the placement of the cell after knocking down respective protein target...84

### Preamble

The recent ENCODE data release provides a rich source of information for investigating questions both in basic biology and human disease. In large part, this wealth of information derives from the multiple genomic annotations provided across multiple cell types. An overarching objective of our study is to leverage ENCODE data to provide novel insights and resources for cancer research. We performed large-scale integration of various assays to construct a companion resource to the general encyclopedia focused on top-tier cell lines and their relevance to cancer biology. In particular, we aim to integrate ENCODE data to gain a more comprehensive understanding of the non-coding regulations involved in oncogenesis, their associated linkages to protein-coding genes, and the global regulatory nature of regulators in networks.

In addition to providing new opportunities, however, the very richness of this data introduces considerable challenges with respect to data integration and organization. Our analyses rely on an array methodology, the details for which are difficult to include within the main text of this paper. As such, the purpose of this Supplement is to provide a clear and organized reference to support and explain the datasets, pipelines, and analyses associated with this study. In addition to supplementary text, the supplementary figures and tables provide additional information not included in the main figures. As reflected in the main text, our study is broadly organized into several main parts:

- A description of the breadth and depth of ENCODE data
- Application #1: Construction of compact individual regulatory elements to form its extended genes
- Application #2: Generalized network analysis to prioritize key regulators and investigate their crosstalk
- Application #3: Tissue-specific network analysis and its application in network rewiring
- Application #4: Cell state transition analysis using expression and network information
- Application #5: The extended gene definitions in expression, germline, and somatic data interpretation
- Workflow for regulators, elements, and nucleotides prioritization

This supplement is presented in roughly a parallel fashion to the main text. It is also connected to the main text through the major results presented in the form of main text figures – captions associated with main text figures point to relevant sub-sections within this supplement. In cases where the related supplementary section wasn't obvious, we said "(see supp. sect xxx) to refer to a specific section. With the aim of presenting data and results (including software packages) in an organized way, we have written about this study in roughly a hierarchical fashion. The main text lies at the top of this hierarchy and synthesizes everything in a broad manner. It refers to more detailed descriptions of our methods and datasets, as provided in this supplement. Raw data files, which lie at the bottom of the hierarchy (and which are hosted as online resources) form the bedrock from which our results are built.

We note that, in preparing this supplement, we adopt the conventions prescribed in the recent opinion piece by Greenbaum et al[1]. As such, labels that correspond to sections, sub-sections, figures, tables, and data files are labeled to indicate whether these items directly parallel (‖) or do not parallel (∦) the main text, as well as whether these items are high-level or technical in nature (designated by "HL" and "TL", respectively).

### 1 More details about "breadth and depth of ENCODE3 data"

#### 1.1 (HL, ‖) Summary of the cancer-related encyclopedia companion resource

Unlike many other diseases, cancer is very much a disease of whole-genome dysregulation. Cancer cells may display aberrant behaviors of key regulators, extensive remodeling of epigenetics, and abnormal transitions between cell states. The wealth of ENCODE functional characterization data allows direct measurement of chromatin status, regulatory changes, and expression perturbations for individual genes. It may also be used to construct comprehensive, high-quality networks, to capture the tumor-to-normal alterations from a more global perspective.

*Figure S 1-1. Summary of ENCODEC*

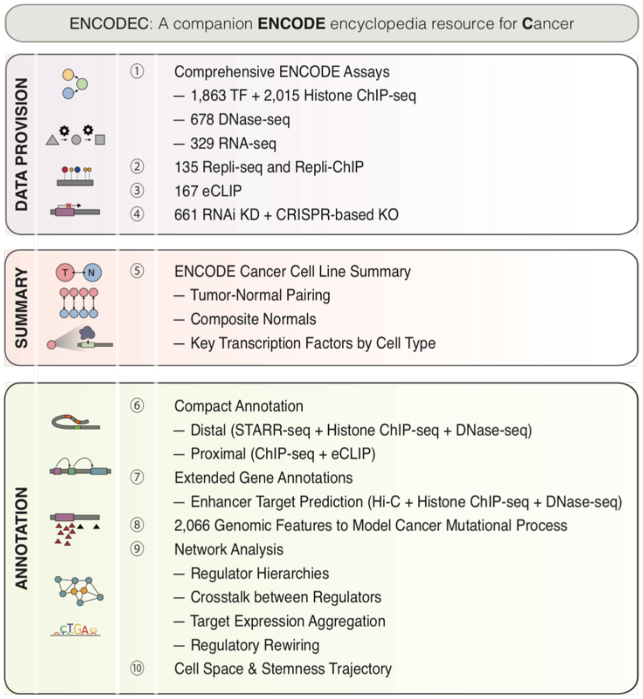

Hence, we try to provide a resource to the main ENCODE encyclopedia by building an "**ENCODE C**ancer-related encyclopedia companion" resource (ENCODEC). The main encyclopedia is oriented toward the breath of the annotations to describe elements over thousands of cell types. In contrast, we focus on top cell types with a wide variety of profiles available. Most of these cell types are associated with cancers of the blood, liver, lung, cervix, and breast. We show that these cell types can be used to provide a better understanding of cancer progression, and we try to provide a resource for interpreting the wealth of mutational and transcriptional profiles produced by the cancer community. We have summarized our efforts in Figure S 1-1. This encyclopedia companion mainly provides three layers of resources: 1) Data provision: carefully collected and de-duplicated signal tracks from various experimental assays both within and outside ENCODE; 2) pairing cell types and datasets to cancer types; 3) Detailed Annotations: enhancers and their gene linkages, cell-type specific and generalized networks, network hierarchies, rewiring status, gene expression regulating potentials. Together, we proposed a step-wise prioritization method to pinpoint key regulators, elements, and SNVs and subject them to experimental validations by several targeted assays.

#### 1.2 (TL, ||) Detailed annotation of TFs and RBPs

In this study, we collected 1,863 TF ChIP-seq experiments in human biosamples. Among them, we selected 344 transcription-related factors (abbreviated them as TFs) and further classified them into four major classes: 282 sequence-specific TFs, which bind DNA at particular motifs to regulate gene expression; 16 general TFs, which comprise that segment of the cell's transcriptional machinery that complexes with DNA; 19 chromatin-associated TFs, which comprise complexes that bind to and remodel chromatin; and 27 co-factors, which support the function of other TFs, do not directly bind DNA, and do not belong to another class. A detailed classification was given in the supplementary datasheet, which is also available from our main data portal (http://encodec.encodeproject.org/).

Of all 103 common TF targets between K562 and GM12878, we further extracted 68 common TFs that have literature support and annotated the TFs in Table S 1-1. We searched the COSMIC Cancer Gene census[2] and an authoritative list of cancer genes by Vogelstein *et al*. to identify TFs associated with cancer[3]. We further listed whether a TF has been reported to regulate the ABL gene or BCR-ABL transcript, or the BCR-ABL KEGG pathway, because of the dominant role this fusion gene plays in CML and K562.

*Table S 1-1. (Shadow table for data in Fig. 1) Detailed Annotation of common TFs in K562 and GM12878*

| TF | Class | FAMILY | COSMIC | Vogelstein | ABL* | BCR-ABL Pathway* | Vogelstein* |
|---|---|---|---|---|---|---|---|
| ATF3 | TFSS | bZIP | 0 | 0 | 1 | 1 | 1 |
| BCLAF1 | TFSS | bZIP | 0 | 0 | 0 | 1 | 1 |
| BHLHE4 0 | TFSS | HLH | 0 | 0 | 0 | 0 | 0 |
| CBX5 | chromatin | | 0 | 0 | 0 | 0 | 0 |
| CEBPB | TFSS | bZIP | 0 | 0 | 0 | 1 | 1 |
| CEBPZ | TFSS | bZIP | 0 | 0 | 0 | 0 | 0 |
| CHD1 | chromatin | Homeodomain | 0 | 0 | 0 | 0 | 0 |
| CHD2 | chromatin | Homeodomain | 0 | 0 | 0 | 0 | 0 |
| CTCF | TFSS | ZNF | 1 | 0 | 0 | 1 | 1 |
| E2F4 | TFSS | wHTH | 0 | 0 | 0 | 1 | 1 |

| | | | | | | | |
|---|---|---|---|---|---|---|---|
| EGR1 | TFSS | ZNF | 0 | 0 | 0 | 1 | 1 |
| ELF1 | TFSS | ETS | 0 | 0 | 0 | 1 | 1 |
| ELK1 | TFSS | ETS | 0 | 0 | 0 | 0 | 0 |
| EP300 | general | | 1 | 1 | 1 | 1 | 1 |
| ETS1 | TFSS | ETS | 0 | 0 | 0 | 1 | 1 |
| ETV6 | TFSS | ETS | 1 | 0 | 0 | 0 | 0 |
| EZH2 | chromatin | | 1 | 1 | 0 | 0 | 0 |
| FOS | TFSS | bZIP | 0 | 0 | 0 | 1 | 1 |
| GABPA | TFSS | ETS | 0 | 0 | 0 | 0 | 0 |
| HDGF | TFSS | | 0 | 0 | 0 | 0 | 0 |
| IKZF1 | TFSS | ZF-C2H2 | 1 | 0 | 0 | 0 | 0 |
| JUNB | TFSS | bZIP | 0 | 0 | 0 | 0 | 0 |
| JUND | TFSS | bZIP | 0 | 0 | 0 | 1 | 1 |
| MAFK | TFSS | bZIP | 0 | 0 | 1 | 1 | 1 |
| MAX | TFSS | HLH | 1 | 0 | 0 | 1 | 1 |
| MAZ | TFSS | HLH | 0 | 0 | 0 | 0 | 0 |
| MEF2A | TFSS | MADs-box | 0 | 0 | 0 | 0 | 0 |
| MLLT1 | TFSS | | 1 | 0 | 0 | 0 | 0 |
| MTA2 | TFSS | ZF-GATA | 0 | 0 | 0 | 0 | 0 |
| MXI1 | TFSS | HLH | 0 | 0 | 0 | 1 | 0 |
| MYC | TFSS | HLH | 1 | 0 | 0 | 1 | 1 |
| NBN | TFSS | | 1 | 0 | 0 | 0 | 0 |

| | | | | | | | |
|---|---|---|---|---|---|---|---|
| NFE2 | TFSS | bZIP | 0 | 0 | 0 | 1 | 0 |
| NFYA | TFSS | CBF-NFY | 0 | 0 | 0 | 1 | 1 |
| NFYB | TFSS | CBF-NFY | 0 | 0 | 0 | 1 | 1 |
| NR2C2 | TFSS | NR | 0 | 0 | 1 | 1 | 1 |
| NRF1 | TFSS | bZIP | 0 | 0 | 1 | 1 | 1 |
| PML | cofactor | | 1 | 0 | 0 | 0 | 0 |
| POLR2A | general | | 0 | 0 | 0 | 0 | 0 |
| POLR3G | general | | 0 | 0 | 0 | 0 | 0 |
| RAD21 | chromatin | | 1 | 0 | 1 | 1 | 1 |
| RCOR1 | TFSS | MYB | 0 | 0 | 0 | 0 | 0 |
| REST | TFSS | ZNF | 0 | 0 | 0 | 1 | 1 |
| RFX5 | TFSS | wHTH | 0 | 0 | 0 | 0 | 1 |
| SIN3A | general | | 0 | 0 | 0 | 1 | 1 |
| SIX5 | TFSS | Homeodomain | 0 | 0 | 0 | 1 | 1 |
| SMAD5 | TFSS | MH1 | 0 | 0 | 0 | 0 | 0 |
| SMC3 | chromatin | | 0 | 0 | 1 | 1 | 1 |
| SP1 | TFSS | ZNF | 0 | 0 | 0 | 1 | 1 |
| SPI1 | TFSS | ETS | 0 | 0 | 0 | 1 | 1 |
| SRF | TFSS | MADs-box | 0 | 0 | 0 | 1 | 1 |
| STAT5A | TFSS | STAT | 0 | 0 | 0 | 0 | 0 |
| SUZ12 | chromatin | ZNF | 1 | 0 | 0 | 1 | 1 |
| TAF1 | general | | 0 | 0 | 0 | 1 | 1 |

| | | | | | | | |
|---|---|---|---|---|---|---|---|
| TARDBP | TFSS | | 0 | 0 | 0 | 0 | 0 |
| TBL1XR1 | cofactor | | 1 | 0 | 0 | 0 | 0 |
| TBP | general | | 0 | 0 | 0 | 0 | 1 |
| UBTF | TFSS | HMG | 0 | 0 | 0 | 0 | 0 |
| USF1 | TFSS | HLH | 0 | 0 | 0 | 1 | 1 |
| USF2 | TFSS | HLH | 0 | 0 | 1 | 1 | 1 |
| YBX1 | TFSS | CSD | 0 | 0 | 0 | 0 | 0 |
| YY1 | TFSS | ZNF | 0 | 0 | 0 | 1 | 1 |
| ZBED1 | TFSS | ZNF | 0 | 0 | 0 | 0 | 0 |
| ZBTB33 | TFSS | ZNF | 0 | 0 | 1 | 1 | 1 |
| ZBTB40 | TFSS | ZNF | 0 | 0 | 0 | 0 | 0 |
| ZNF143 | TFSS | ZNF | 0 | 0 | 0 | 0 | 0 |
| ZNF274 | TFSS | ZNF | 0 | 0 | 1 | 1 | 1 |

To provide functional annotation of the RBPs included in our study, we used gene ontology (GO) categorizations from the Gene Ontology Consortium. Using the Amigo2 webserver (amigo.geneontology.org/goose), we selected 17 GO categories, each representing a major functional category of RBP function, and overlapped these with the list of RBPs generated by the ENCODE project. Our chosen functional categories correspond to RNA binding (GO:0003723), tRNA binding and splicing (GO:0000049 and GO:0006388), RNA splicing (GO:0043484 and GO:0000398), RNA polyadenylation (GO:0043631, GO:0006378, and GO:1900363), regulation of RNA stability (GO:0043488 and GO:0061157), rRNA processing/ribosome (GO:0006364 and GO:0003735), RNA editing (GO:0009451), and snoRNA binding (GO:0030515). Of the ~1000 ENCODE annotated RBPs, 553 are listed as "RNA binding" by GO, and 327 have at least one specific functional annotation that involves RNA binding. A summary of the RBP annotations have been listed in Figure S 1-2.

*Figure S 1-2. Summary of RBPs used in this paper*

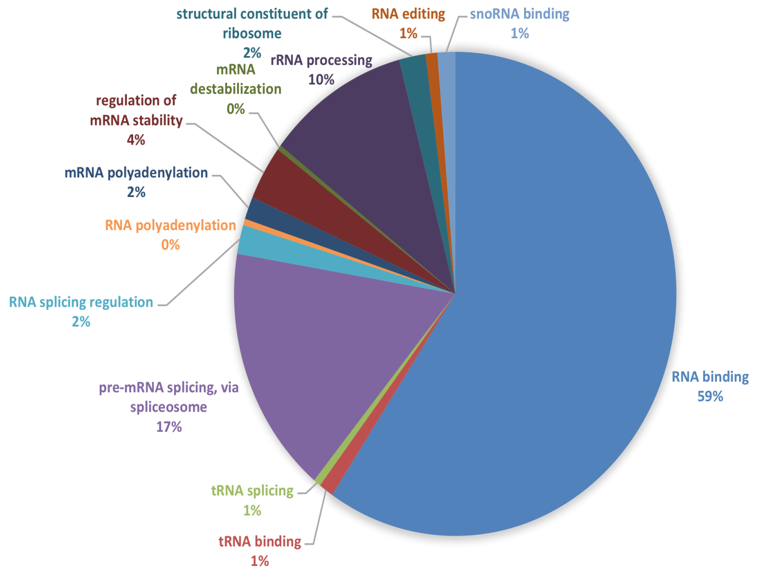

#### 1.3 (HL, ‖) Matching of ENCODE cell types to major cancer types

##### 1.3.1 (HL, ‖) ENCODE data is suitable for cancer analysis

Of all the cell lines in ENCODE, we found around 70 percent of them to be cancerous (shown in Figure S 1-3). Many of the cell types are quite enriched with various experimental assays for functional characterization, leaving the ENCODE data quite suitable for cancer research (Figure S 1-4).

*Figure S 1-3. Summary of the ENCODE cell lines*

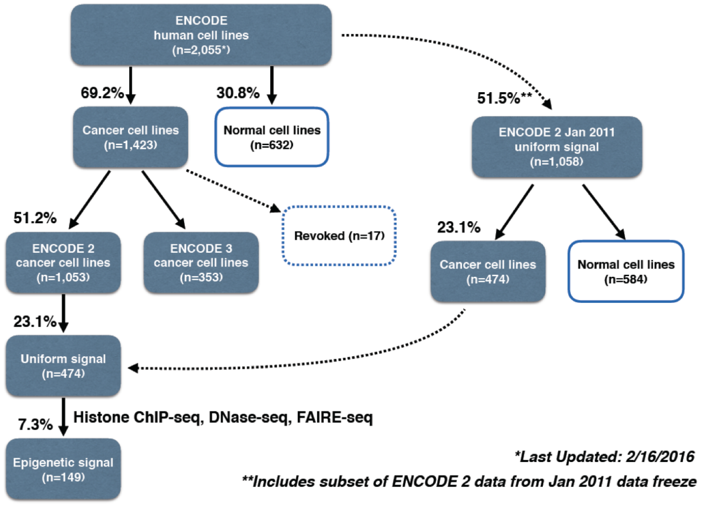

*Figure S 1-4. Broad spectrum of ENCODE cell types*

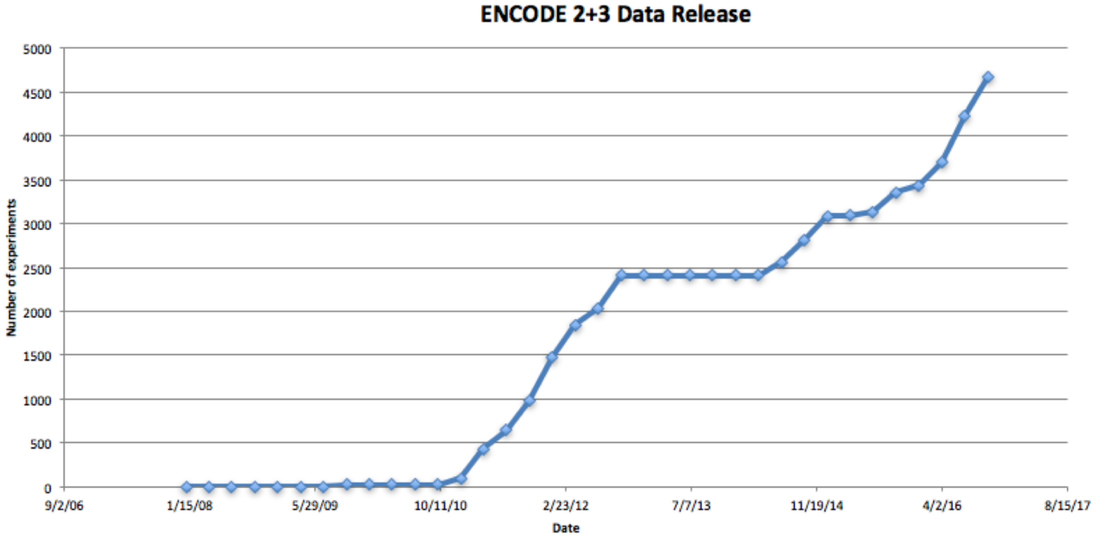

#### 1.4 (HL, ∦) Rationale for matching cell types

Although wide-ranging functional characterization assays are available through ENCODE, deep integration of this to develop an annotation resource for disease studies remains challenging. For example, in cancer, biological heterogeneity among cancer types requires that cell types studied in cancer research be optimally matched to a cancer of interest. However, well-matched tumor-normal pairs are only available for certain cancer types from among ENCODE cell types, and most cell types lack data from one or more key experimental assays (as shown in Fig. 1). Therefore, it is necessary to create biologically relevant tumor-normal pairs and to develop algorithms to learn from sub-optimally matched data. Another challenge arises from the heterogeneous raw data collected from experimental assays. These raw data must undergo de-duplication, unified processing, and proper normalization before accurate large-scale integration can be achieved. Here we endeavor to provide ENCODE data matched according to cancer type, and with appropriate tumor-normal pairs, for several high-incidence cancer types. A detailed matching summary is provided in Table S 1-2.

To utilize the extensive collection of advanced, ENCODE assays on several “key cell types”, one can perform an approximate matching of cancerous cells to normal cells. For the normal cells, we could simply combine a collection of normal proxies to form a composite normal. In this way, we can maximize the functional coverages of approximately matched normal while minimizing artifacts by diluting unwanted cell type-specific noises.

A key feature of the ENCODE annotation is that it includes data from a great diversity of functional assays. Because it is not currently possible to perform such a wide variety of assays on tissue from individual cancer patients, we believe that this matching provides a unique opportunity to refine our understanding of the cancer genome, using large-scale data integration.

*Table S 1-2. Summary of cell line and cancer type matching*

| Cancer | Abbreviation | ENCODE cell line | |
|---|---|---|---|
| Breast | BRCA | Tumor | MCF-7 |

| | | | |
|---|---|---|---|
| | | Normal | HMEC, MCF-10A |
| Liver | LIHC | Tumor | HepG2 |
| | | Normal | Fetal + Adult liver tissue |
| Lung | LUAD [SARC] | Adenocarcinoma | A549 |
| | | Sarcoma | SK-N-MC |
| | | Normal | Lung tissue, IMR-90 |
| Blood | CML [CLL/LAML] | Tumor | K562, DND-41 |
| | | Normal | CD34+ common myeloid progenitor cell, GM12878 |
| Cervix | CESC | Tumor | HeLa-S3 |
| | | Normal | N/A |
| Colorectal | COAD+READ | Adenocarcinoma | Caco-2, HCT116 |
| Prostate | PRAD | Adenocarcinoma | LNCaP, PC-3 |
| Pancreas | PAAD | Adenocarcinoma | Panc1 |

##### 1.4.1 (TL, ∦) Blood cancer cell line matching

Wherever possible, we have matched each ENCODE cancer cell type with a composite normal, which was derived from multiple of normal samples with the same cell type of origin. Exact matching was not possible with K562: this cancer cell-line derives from a myeloid lineage, but there is no data-rich noncancerous myeloid cell included in ENCODE. GM12878 is a data-rich ENCODE cell-line derived from the closely related lymphoid lineage. Supporting this choice, we determined that among all non-cancerous cell-lines provided by Roadmap Epigenome and GTEx, GM12878 has the highest Spearman correlation with K562, as shown in Figure S 1-5. Hence, we used GM12878 as a ***rough*** pair for K562.

*Figure S 1-5.* Expression Matching with K562

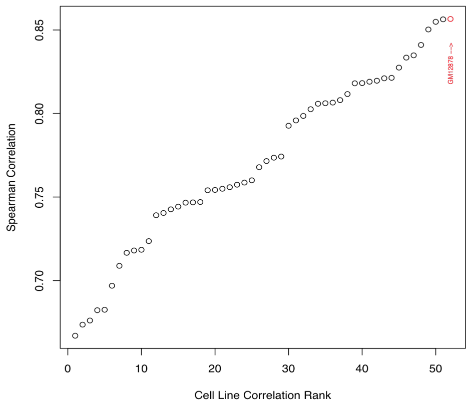

In addition, we found CD34+ common myeloid progenitor cells to be another close normal proxy to K562. Common myeloid progenitor cells are a direct ancestor of many differentiated myeloid cells including granulocytes and monocytes, and therefore it can directly be related to chronic myeloid leukemia (CML). However, there is a limited range of data assays with CD34+

common myeloid progenitor cell, and therefore we merged them with data-rich GM12878 assays to build a composite normal.

##### 1.4.2 (TL, ∦) Breast cancer cell line matching

MCF-7 is the most studied human breast cancer cell line, with nearly 25,000 scientific publications reporting results from studies of MCF-7[4]. It is a human cell line derived from a malignant pleural effusion due to breast carcinoma[5]. MCF-7 is one of few cell lines that express substantial levels of estrogen receptor (ER), and so is widely used to mimic ER-positive invasive human breast cancers. It is also used to study intracellular binding constants, transport mechanisms, and DNA binding sites among ER target genes[4]. T47D is also an ER-positive breast cancer cell line that has been widely used to study breast cancer and is also derived from a malignant pleural effusion[6]. Unlike MCF-7, T47D is a mutant for the tumor suppressor gene TP53[7].

MCF-10A is the human breast epithelial cell line most commonly used as an *in vitro* model for studying normal breast cell function and transformation[8]. It was derived from spontaneously immortalized benign fibrocystic mammary tissue, which is non-tumorigenic and does not express ER[8,9]. Numerous studies have utilized both MCF-7 and MCF-10A cell lines to facilitate the development of breast cancer treatment and therapy, by comparing the differential response of these two cell lines under multiple experimental settings[10-12].

A recent study challenges MCF-10A as a representative model for normal mammary cells, with the study authors claiming that this cell line exhibits phenotypes and expression profiles that have not been observed in mammary gland tissues[8]. However, these authors expressed a need for further investigation into the appropriateness of MCF-10A cells as a model for normal human mammary epithelial cells. Though we cannot exclude differences between MCF-7 and MCF-10A from causes unrelated to malignant transformation, given the wealth of ENCODE data on MCF-7, and the high incidence of breast cancer, we consider the pairing of MCF-7 and MCF-10A worthwhile.

Human mammary epithelial cells (HMEC) is another representative model for normal mammary cells and numerous studies have used them for matching normal against MCF-7[13-15]. HMEC is also widely used outside ENCODE and we could obtain external data such as Hi-C, from a literature search (GSM1551613).

##### 1.4.3 (TL, ∦) Lung cancer cell line matching

A549 is a carcinomic lung epithelial cell line[16] and IMR-90 is a normal lung fibroblast cell line[17]. Lung fibroblasts and lung epithelial cells are closely related cell types, and conversion between these cell types is common and meaningful in tumor cells and normal cells[18,19]. Lung fibroblasts, like IMR-90, are mesenchymal cells that arise in embryologic development subsequent to epithelial to mesenchymal transition (EMT). The dedifferentiation of mesenchymal cells into secondary epithelial tissue following mesenchymal to epithelial transition (MET) is also observed and is best characterized in kidney development[20]. It has been postulated that the dedifferentiation and metastasis of epithelial lung cancer cells, may occur through EMT and/or MET[21-24]. Such a process has been observed in other cancers[18]. Indeed, exposure of A549 epithelial cells to chemotherapeutic agents or TGF-B, causes differentiation to a mesenchymal phenotype, and EMT is thought to play a role in chemotherapeutic resistance of lung adenocarcinoma[25,26]. These cellular relationships support the benefit of using a tumor normal comparison between A549 cancer cells and IMR-90 normal cells. Indeed, IMR-90 is frequently used as a normal control for A549 in experiment[27-33].

ENCODE contains a wide range of data-rich assays on both fetal and adult lung tissues, including but not limited to DNase-seq, ChIP-seq, and RNA-seq. Not a single biosample can be used as a perfect match for A549, and therefore, we merged data-rich assays on normal lung biosamples from anatomical origin (IMR-90, fetal and adult lung tissue samples) or cellular origin (any normal epithelial cells) to form a composite normal.

##### 1.4.4 (TL, ∦) Normal to tumor cell line matching using replication timing data

It is well known that replication timing significantly affects the mutational landscape in both germline and normal cells[3437]. We also made a genome-wide correlation of replication timing data (excluding ChrX and ChrY to avoid gender differences) between the cancer cell lines and several candidate normal cell types. Results are listed in Figure S 1-6.

As expected, the best matching normal data for K562 and HepG2 cell lines are Erythroid progenitors and Hepatocytes. However, we also noticed that replication timing data in A549 and MCF-7 shows the highest correlation with those in Mesenchymal Stem cells and Splanchnic mesoderm. However, our proposed matching normal cell lines, such as like IMR-90 for A549, still showed a decent correlation regarding their replication timing profiles.

*Figure S 1-6. Comparison of tumor with normal cell lines by replication timing data*

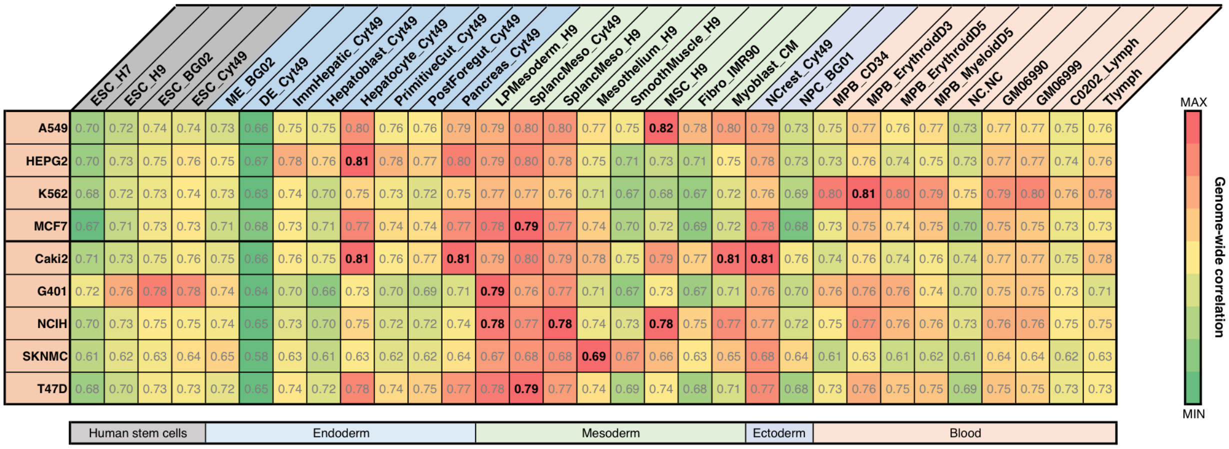

| | ESC_H7 | ESC_H9 | ESC_BG02 | ESC_Cyt49 | ME_BG02 | DE_Cyt49 | ImmHepatic_Cyt49 | Hepatoblast_Cyt49 | Hepatocyte_Cyt49 | PrimitiveGut_Cyt49 | PostForegut_Cyt49 | Pancreas_Cyt49 | LPMesoderm_H9 | SplancMeso_Cyt49 | SplancMeso_H9 | Mesothelium_H9 |
|---|---|---|---|---|---|---|---|---|---|---|---|---|---|---|---|---|
| A549 | 0.70 | 0.72 | 0.74 | 0.74 | 0.73 | 0.66 | 0.75 | 0.75 | 0.80 | 0.76 | 0.76 | 0.79 | 0.79 | 0.80 | 0.80 | 0.77 |
| HEPG2 | 0.70 | 0.73 | 0.75 | 0.76 | 0.75 | 0.67 | 0.78 | 0.76 | **0.81** | 0.78 | 0.77 | 0.80 | 0.79 | 0.80 | 0.78 | 0.75 |
| K562 | 0.68 | 0.72 | 0.73 | 0.74 | 0.73 | 0.63 | 0.74 | 0.70 | 0.75 | 0.73 | 0.72 | 0.75 | 0.77 | 0.77 | 0.76 | 0.71 |
| MCF7 | 0.67 | 0.71 | 0.73 | 0.73 | 0.71 | 0.68 | 0.73 | 0.71 | 0.77 | 0.74 | 0.74 | 0.77 | 0.78 | **0.79** | 0.77 | 0.74 |
| Caki2 | 0.71 | 0.73 | 0.75 | 0.76 | 0.75 | 0.66 | 0.76 | 0.75 | **0.81** | 0.76 | 0.77 | **0.81** | 0.79 | 0.80 | 0.79 | 0.78 |
| G401 | 0.72 | 0.76 | 0.78 | 0.78 | 0.74 | 0.64 | 0.70 | 0.66 | 0.73 | 0.70 | 0.69 | 0.71 | **0.79** | 0.76 | 0.77 | 0.71 |
| NCIH | 0.70 | 0.73 | 0.75 | 0.75 | 0.74 | 0.65 | 0.73 | 0.70 | 0.75 | 0.72 | 0.72 | 0.74 | **0.78** | 0.77 | **0.78** | 0.74 |
| SKNMC | 0.61 | 0.62 | 0.63 | 0.64 | 0.65 | 0.58 | 0.63 | 0.61 | 0.63 | 0.62 | 0.62 | 0.64 | 0.67 | 0.68 | 0.68 | **0.69** |
| T47D | 0.68 | 0.70 | 0.73 | 0.73 | 0.72 | 0.65 | 0.74 | 0.72 | 0.78 | 0.74 | 0.75 | 0.77 | 0.78 | **0.79** | 0.77 | 0.74 |

| | SmoothMuscle_H9 | MSC_H9 | Fibro_IMR90 | Myoblast_CM | NCrest_Cyt49 | NPC_BG01 | MPB_CD34 | MPB_ErythroidD3 | MPB_ErythroidD5 | MPB_MyeloidD5 | NC.NC | GM06990 | GM06999 | C0202_Lymph | Tlymph |
|---|---|---|---|---|---|---|---|---|---|---|---|---|---|---|---|
| A549 | 0.75 | **0.82** | 0.78 | 0.80 | 0.79 | 0.73 | 0.75 | 0.77 | 0.76 | 0.77 | 0.73 | 0.77 | 0.77 | 0.75 | 0.76 |
| HEPG2 | 0.71 | 0.73 | 0.71 | 0.75 | 0.78 | 0.73 | 0.73 | 0.76 | 0.74 | 0.75 | 0.73 | 0.77 | 0.77 | 0.74 | 0.76 |
| K562 | 0.67 | 0.68 | 0.67 | 0.72 | 0.76 | 0.69 | 0.80 | **0.81** | 0.80 | 0.79 | 0.75 | 0.79 | 0.80 | 0.76 | 0.78 |
| MCF7 | 0.70 | 0.72 | 0.69 | 0.72 | 0.78 | 0.68 | 0.73 | 0.75 | 0.74 | 0.75 | 0.70 | 0.75 | 0.75 | 0.73 | 0.73 |
| Caki2 | 0.75 | 0.79 | 0.77 | **0.81** | **0.81** | 0.76 | 0.74 | 0.76 | 0.74 | 0.76 | 0.74 | 0.77 | 0.77 | 0.75 | 0.78 |
| G401 | 0.67 | 0.73 | 0.67 | 0.71 | 0.76 | 0.70 | 0.76 | 0.76 | 0.76 | 0.74 | 0.70 | 0.75 | 0.75 | 0.73 | 0.71 |
| NCIH | 0.73 | **0.78** | 0.75 | 0.77 | 0.77 | 0.72 | 0.75 | 0.77 | 0.76 | 0.76 | 0.73 | 0.76 | 0.76 | 0.75 | 0.75 |
| SKNMC | 0.67 | 0.66 | 0.63 | 0.65 | 0.68 | 0.64 | 0.61 | 0.63 | 0.61 | 0.62 | 0.61 | 0.64 | 0.64 | 0.62 | 0.63 |
| T47D | 0.69 | 0.74 | 0.68 | 0.71 | 0.77 | 0.68 | 0.73 | 0.76 | 0.75 | 0.75 | 0.69 | 0.75 | 0.75 | 0.73 | 0.73 |

#### 1.5 (TL, ‖) Summary of data from each experimental assay from ENCODE

We have integrated data sets from ENCODE and Roadmap Epigenomics Mapping Consortium (REMC), after performing quality control and uniform processing, to build one of the most comprehensive representations of how functional regulatory elements interplay in the human genome. All datasets used in the analysis were mapped to a standardized version of the GRCh37 (hg19) reference human genome. We used ENCODE data that was submitted and released up to October 31st, 2016 (Oct 2016 freeze). We summarized the ENCODE data with matched tumors in Figure S 1-7.

*Figure S 1-7. (Shadow figure) Summary of ENCODE data for multiple cancer types*

| | K562 | GM12878 | HepG2 | liver | A549 | lung | MCF-7 | HMEC | HeLa-S3 | H1-hESC | Caco-2 | HCT116 | Panc1 | LNCaP | PC-3 | PC-9 | SK-N-MC | DND-41 | SK-N-SH |
|---|---|---|---|---|---|---|---|---|---|---|---|---|---|---|---|---|---|---|---|
| DNase-seq | 10 | 3 | 3 | 2 | 14 | 13 | 8 | 3 | 4 | 3 | 1 | 1 | 1 | 3 | 1 | 1 | 1 | | 2 |
| Histone | 19 | 14 | 14 | 25 | 85 | 25 | 16 | 26 | 14 | 53 | 3 | 16 | 7 | 1 | 11 | 11 | 8 | 11 | 19 |
| polyA RNA-seq | 19 | 14 | 11 | 5 | 27 | 4 | 5 | 3 | 5 | 10 | | 1 | 1 | | | | | | 5 |
| RAMPAGE | 1 | 1 | | 2 | | 1 | | | | | | | | | 1 | | | | |
| eCLIP | 191 | | 164 | | | | | | | | | | | | | | | | |
| RNAi KD | 326 | | 257 | | | | 2 | | | | | | | | | | | | |
| CRISPR KD/KO | 108 | | 19 | | | | | | | | | | | | | | | | |
| ChIA-PET | 9 | 2 | 2 | | | | 5 | | 1 | | | 1 | | 2 | | | | | |
| Hi-C | | | 1 | | 6 | | | | 1 | | | | 1 | 1 | | | 1 | | |
| STARR-seq | 1 | 1 | 1 | | | | 1 | | | | | | | | | | | | |
| WGBS | 1 | 2 | 2 | 3 | 1 | 3 | | 2 | 1 | 3 | | | | | | | | | 1 |
| RRBS | 1 | 2 | 2 | 1 | 1 | 3 | 5 | 1 | 1 | 6 | 1 | 1 | 2 | 3 | | | 1 | | 2 |
| Repli-chip | | | | | | | | | 1 | 1 | | | | | | | | | |
| Repli-seq | 6 | 6 | 6 | | 2 | | 6 | | 6 | | | | | 2 | | | 2 | | 6 |
| TF | 558 | 209 | 300 | 35 | 240 | 1 | 149 | 4 | 78 | 89 | 1 | 28 | 7 | 4 | 5 | 5 | 3 | 3 | 44 |
| SNV | 1 | 1 | | | | | 1 | | 1 | | | | | | | | | | |
| SV | 1 | 1 | | | | | 1 | | 1 | | | | | | | | | | |

##### 1.5.1 (TL, ǂ) Preprocessing of Repli-seq data

Replication timing (RT) datasets of cell types representing distinct stages of cell differentiation of human embryonic stem cells towards endoderm, mesoderm, ectoderm, and neural crest were obtained from Rivera-Mulia et al[35]. Genome-wide RT analysis of the ENCODE cancer cell lines (A549, Caki2, G401, NCI-H460, SK-N-MC, T47D and LNCaP) was performed by NGS as previously described[36,37]. Briefly, cells were pulse-labeled with BrdU and separated into early and late S-phase fractions by FACS, followed by DNA immunoprecipitation with anti-BrdU antibody (Sigma-Aldrich, cat. no. B5002). Sequencing libraries of BrdU-substituted DNA from early and late fractions were prepared using the NEBNext Ultra DNA Library Prep kit for Illumina (E7370). Sequencing was performed on an Illumina-HiSeq 2500. Approximately 10 million reads per sample were generated. Reads with quality scores above 30 were mapped to the Hg19 reference genome using Bowtie 2[38]. Approximately 8 million uniquely mapped reads were obtained from each library. Read counts were binned into 10-kb nonoverlapping windows, and log2 ratios of read-counts between early and late fractions were calculated. For HepG2, K562, MCF7 and SK-N-SH cell lines, raw data for 6-fraction Repli-seq was downloaded from the ENCODE portal. The data was transformed to match the early/late repli-seq by combining G1, S1 and S2 fractions to represent early S phase and S3, S4 and G2 fractions to represent the late S phase. RT datasets were normalized using the limma package in R (R Core Team 2017) and rescaled to equivalent ranges by quantile normalization.

##### 1.5.2 (TL, ‖) Deduplication of ChIP-seq data

As of March 2018, there are 1,863 TF ChIP-seq experiments and 2,015 histone modification ChIP-seq experiments available for download in ENCODE portal. We downloaded all available IDR thresholded peaks and signal tracks for both fold enrichment and log likelihood p-value in bigWig format and used for downstream analysis.

ENCODE data has redundancies from ENCODE phase 2 and phase 3, as well as different labs have produced the same ChIP-seq datasets using different experimental protocol. Here, we aim to create a subset ENCODE ChIP-seq dataset that are uniform and non-redundant for downstream analysis. For deduplicated dataset, we used ENCODE data that was submitted and released up to October 31st, 2016 (Oct 2016 freeze). We limited TF ChIP-seq experiments to ones that either had no treatment or ethanol treatment only. We used a subset of **1,040** TF ChIP-seq experiments released for ENCODE phase 2 (n=604) and 3 (n=436) that either had no treatment or ethanol treatment only. We carefully manually de-duplicated the dataset by selecting one TF ChIP-seq experiment per each sample by the following prioritization scheme. When an ENCODE3 experiment was available, it was prioritized over the ENCODE2 experiment. When there was the same type of experiments done by different labs, we prioritized using the following order determined by the total number of ChIP-seq experiments deposited on ENCODE: *stanford*, *haib*, *broad*, *usc*, *uw*, *uta*, *uchicago*, *hms*, *yale*. We removed epitope-tagged experiments if an endogenous antibody was available. After deduplication, there were **860** unique TF ChIP-seq experiments.

*Table S 1-3. Number of unique TFs in each cell type*

| Cell Type | # of Unique TFs |
|---|---|
| K562 | 357 |
| MCF-7 | 90 |
| GM12878 | 145 |
| A549 | 47 |
| HepG2 | 224 |
| HEK293 | 201 |
| HeLa-S3 | 61 |

#### 2 More details about "Construction of the ENCODEC resource"

##### 2.1 Compact individual annotation from ENCODE cell types

###### 2.1.1 (TL, ‖) Defining proximal regulatory regions

Of the eCLIP data from ENCODE, 87 are from K562 and 69 are from HepG2. Replicates of the same RBP are merged and a cutoff of significance score 1000 is used to determine the high confidence peaks[39].

*Table S 2-1 Number of RBPs per cell type*

| Cell Type | # of Unique RBPs |
|---|---|
| K562 | 87 |
| HepG2 | 69 |

After deduplication of the TFs, we have intersected the TF peaks within 2.5 kb of the transcription start sites (TSS) of the coding transcripts as the potential proximal regulatory sites.

###### 2.1.2 (TL, ‖) Integrative enhancer definitions

In contrast to previous approaches to enhancer annotations (many of which use only histone modification and chromatin accessibility data), we proposed an ensemble method to accurately pinpoint active enhancers and link them to protein coding genes. It composes three computational pipelines (MatchedFilter, ESCAPE, and JEME) to integrate tens of data sets from six different experimental assays, including ChIP-seq, DNase-seq, STARR-seq (CapSTARR-seq), RNA-seq, ChIA-PET, and Hi-C for higher accuracy. The overall schematic has been summarized into Figure S 2-1.

*Figure S 2-1. Overall schematic of enhancer and gene linkage*

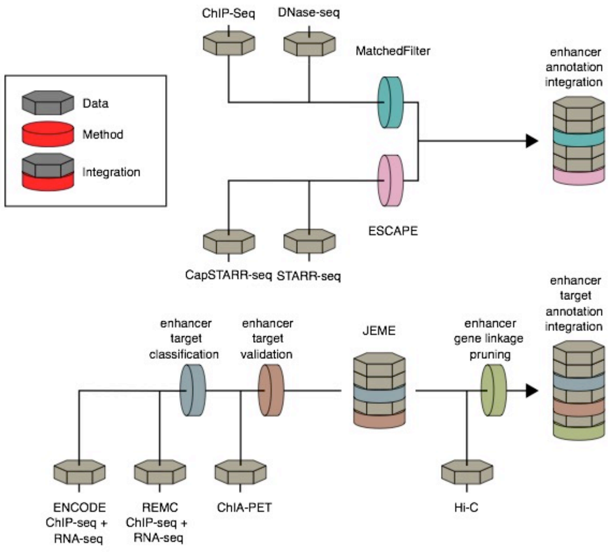

For the enhancer prediction part, we integrated prediction results from different methods and data source, as shown in Figure S 2-1. First, we proposed a computational method, called MatchedFilter, to make enhancer predictions based on pattern recognitions of histone marks**.** The upper white panel of Figure S 2-2 shows an example of the signal tracks (DNase, H3K27ac, H3K4me1 and H3K3me3) of GM12878. Based on these signals and other experiment results, we have a collective data source including enhancers predicted from MatchedFilter and ESCAPE, an enhancer-gene linkage map from JEME and Hi-C, ccREs from Phase III encyclopedia and transcription factor (TF) binding sites (middle panel). Specifically, we took intersections of predicted enhancers from MatchedFilter and ESCAPE and further filtered them from target predictions with JEME and Hi-C results. The resulting list is further refined by intersecting with ccREs from the main encyclopedia paper. In the last step, we pruned the enhancers in the list to trim both ends that are not covered by any transcription factor binding motifs (TFBS). The final list is a set of precisely refined and high-confident enhancers.

*Figure S 2-2. Integrated approach to generate high-confidence enhancers list*

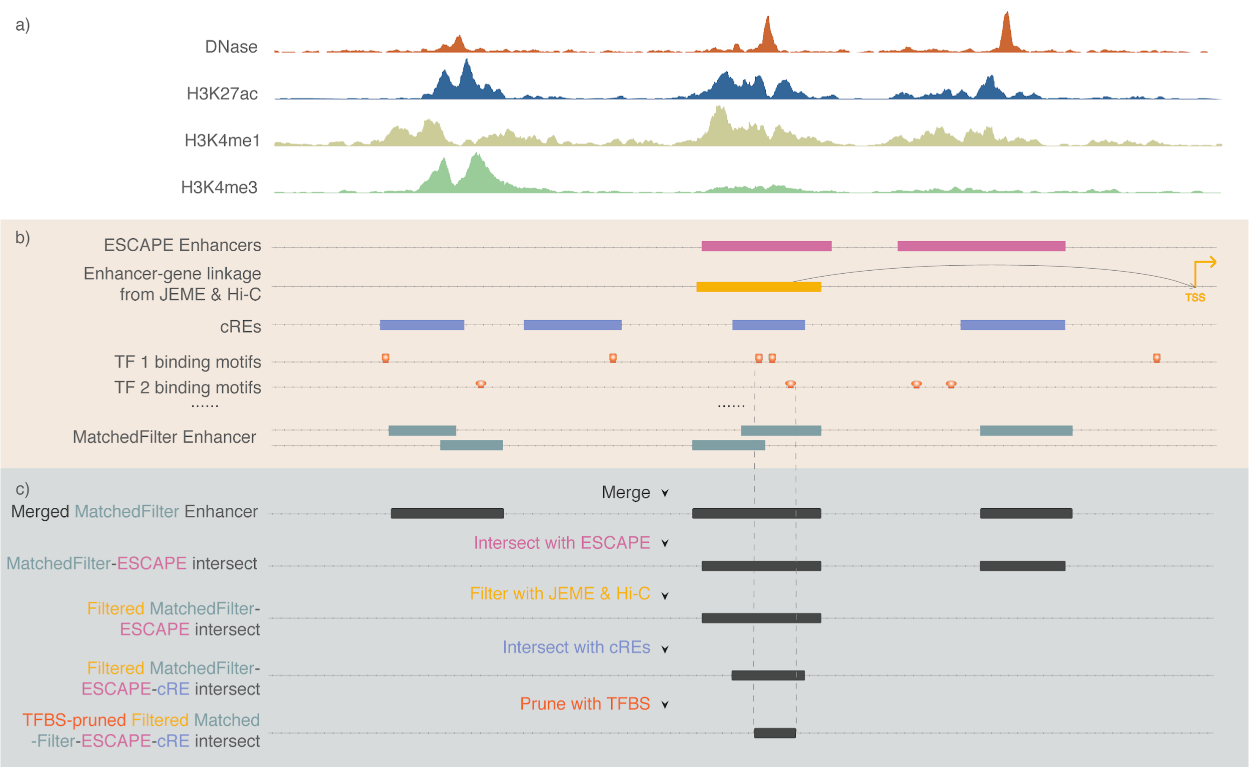

##### 2.1.2.1 (TL, ‖) Enhancer prediction Pipeline based on the MatchedFilter.

We developed a framework to detect enhancer regions across the genome through aggregated signals of epigenetic features[40]. Briefly, since the aggregated H3K27ac signal at the genome-wide STARR-seq peaks exhibits an enriched peak-trough-peak pattern[41], a framework was designed to learn the aggregated patterns for different histone modification signals at enhancer regions, and then use them to scan the whole genome ChIP-seq data from tissues or cell lines using the canonical signal processing method called MatchedFilter[40]. We use linear SVM to assemble the matched filter scores to form a discriminant function, where the sign of the result value is used to predict whether a specific region is an enhancer. After applying this framework to predict enhancers in several normal and cancer-related cell lines, including GM12878, HepG2, K562 and MCF-7, the number of predicted enhancers is provided in Table S 2-2.

*Table S 2-2.* Number of enhancers predicted by histone-shape based method

| Cell Type | GM2878 | HepG2 | K562 | MCF-7 |
|---|---|---|---|---|
| Number of Enhancers | 45,202 | 61,005 | 45,801 | 59,827 |

###### 2.1.2.2 (TL, ‖) Enhancer prediction by EnhancerS Peak CAlling PipEline from STARR-seq (ESCAPE)

The whole-genome STARR-seq was performed using a protocol conceptually similar to the previously published STARR-seq technique that was done in the *Drosophila* melanogaster genome[49]. The CapSTARR-seq is a variant of STARR-seq technique which combines STARR-seq with genome capturing technology[50]. In brief, the genomic DNA from each cell line was fragmented into ~500 bp by sonication and built into plasmid library, which was named as screening library. The screening library was subjected for next generation sequencing. We verified the sequence complexity and genome coverage of screening libraries, which were then transfected into GM12878, K562 or MCF-7 cells by electroporation. After 24 hours of transfection, the plasmid-specific mRNA was purified, reverse transcribed and PCR amplified. The PCR products, which are the so-called STARR-seq libraries, were subjected to sequencing. Both screening library and STARR-seq libraries were sequenced as 100 bp paired-end on the Illumina HiSeq 2500/4000 platforms. The general workflow of the MCF-7 CapSTARR-seq is similar to the whole-genome STARR-seq, however, we captured ~10,000 DNase I hypersensitivity sites (a total length of 9.7 Mb) from fragmented genomic DNA to build the screening library. Compared to the published STARR-seq work, we'd like to note the following innovation and improvement: (1) We significantly increased the complexity of the screening libraries to ensure comprehensive coverage to the human genome; (2) We significantly increased the electroporation scale and efficiency to maximize the size of screening library that got into the cells; (3) We introduced an extra multiplexing step to minimize the bias introduced by PCR duplicates. See details in Figure S 2-3.

For the capture-based assay for MCF-7 cell line, a total of 10,825 target regions consisting of 9,825 candidate enhancer regions and 1,000 negative control regions were selected tested for regulatory potential. Candidate enhancer regions were selected based on DHS peaks excluding both 1kb upstream and downstream of TSS. Negative control regions were selected from 500 randomly selected regions and 500 non-E2-responsive DHS regions. Details of the selection procedure can be found in Figure S 2-4. (L. Ma et al for GM12878 and K562 whole-genome STARR-seq; S. Yu et al for MCF-7 CapSTARR-seq, in preparation). Candidate enhancer regions were primed and inserted into 3’ UTR. Schematics of the experimental procedure can be found in Figure S 2-3.

*Figure S 2-3. Schematic of the STARR-seq protocol*

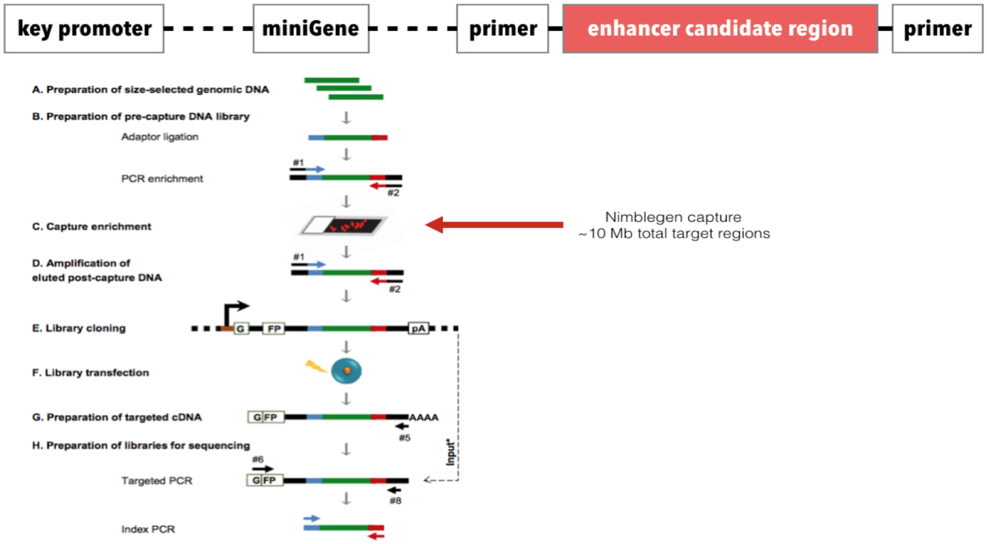

*Figure S 2-4. Schematic of capture STARR-seq*

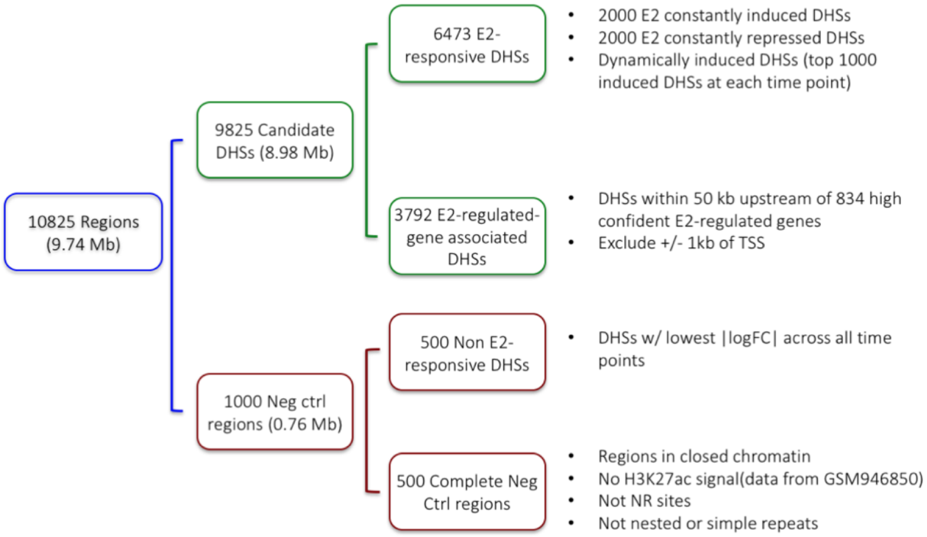

*Figure S 2-5. Schematic of ESCAPE*

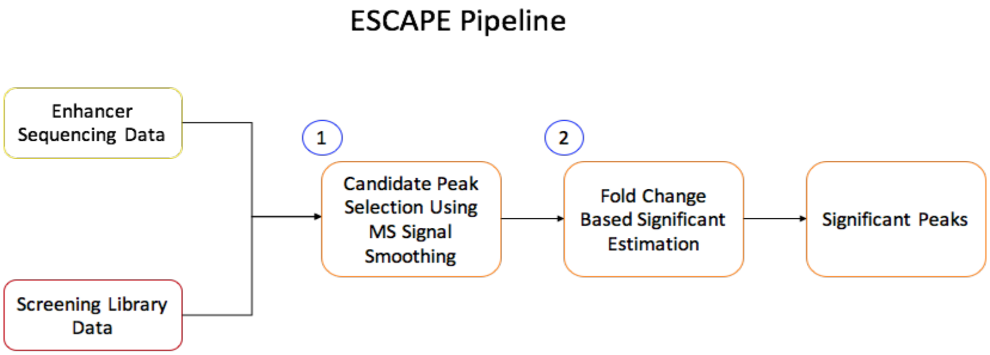

We proposed a peak calling method for whole genome STARR-seq called ESCAPE using the following strategy: First, the peak candidates are identified. For the whole genome assay, ESCAPE uses a multiscale decomposition based peak calling strategy[53]. For this, we have decomposed the signal using smoothing filters with lengths varying between 100 and 2000 base pairs. The filtering can be summarized with following formula:

$$x_i^s = median\left(\{\tilde{x}_a\}_{a\in\left[i-\frac{l_s}{2}, i+\frac{l_s}{2}\right]}\right), l_s \in (\, l_{start}, \lfloor l_{start} \times \sigma \rfloor, \cdots, l_{end}) \quad (2\text{-}1)$$

where $x_i^s$ is the $i^{th}$ signal level at scale decomposition $s$. The smoothing window length is $l_s$. Then we identified the local minima in the smoothed signal profiles and used these as possible enriched regions. For this, ESCAPE first estimates the derivative at each point:

$$dx_i^s = (x_i^s - x_{i-1}^s) \quad (2\text{-}2)$$

where $dx_i^s$ is the derivative of the smoothed signal $x_i^s$. The local extrema are found as the points where the derivative flips its sign:

$$I_{min} = \{i \mid dx_i^s < 0, dx_{i-1}^s > 0\} \quad (2\text{-}3)$$

$$I_{max} = \{i \mid dx_i^s > 0, dx_{i-1}^s < 0\} \quad (2\text{-}4)$$

where $I_{min}$ and $I_{max}$ are the sets of positions of minima and maxima of $x_i^s$, respectively. The scale specific candidate enriched regions of $x_i^s$ are identified as the regions between the consecutive minima. The multiscale decomposition approach identifies enriched regions at different length scales that correspond to punctate features like enhancers. Then, ESCAPE computes the fold change on each peak candidate as the ratio of total signal in the STARR-seq signal and screening library signal. We refer to this as $FC$:

$$FC = \frac{\sum_{i=s}^{e} x_i^s}{\sum_{i=s}^{e} y_i^s} \quad (2\text{-}5)$$

where $y_i^s$ represents the value of screening library signal profile at position $i$. For capture based assay, ESCAPE uses a more focused analysis to identify candidate peak regions. For each capture region, ESCAPE selects a bins size that balances the peak calling sensitivity and specificity. To set a threshold for the fold change to select candidate peaks, we exchanged screening library and STARR-seq and we computed the fold change on the candidate peaks, which we refer to as $FC_{random}$:

$$FC_{random} = \frac{\sum_{i=s}^{e} y_i^s}{\sum_{i=s}^{e} x_i^s} \quad (2\text{-}6)$$

These fold change scores serve as a random distribution of fold change scores. We use this distribution for selecting a fold change threshold. For a $FC$ threshold $\boldsymbol{fc}$, we estimated the false discovery rate as the ratio of number of peaks that for which $FC_{random} > \boldsymbol{fc}$ and the number of peaks for which $FC > \boldsymbol{fc}$. We set the FDR threshold at 0.1% and filtered the peaks that do not satisfy the $FC$ threshold selected using this FDR threshold. For capture based assay, ESCAPE uses the candidate enriched regions with top 10% $FC$ values.

#### 2.2 Linking compact individual annotations to genes to form the extended gene annotations

##### 2.2.1 (TL, ||) Enhancer Gene linkage prediction using JEME

Enhancer targets were predicted using JEME[42] (Joint Effect of Multiple Enhancers), which involves two main steps (Figure S 2-6). In the first step, the transcript levels around each transcription start site (TSS) in 49 ENCODE and Roadmap Epigenomics cell lines were modeled based on histone modification data at nearby enhancers without requiring any known enhancer-target pairs as examples. Specifically, for each enhancer feature the expression level of a TSS is modeled, where the summation is over all enhancers within 1Mbp from the TSS and is the value of the feature of the enhancer. The coefficients of the enhancers are learned by LASSO, which minimizes the regression error over all samples while selecting a small number of enhancers to have non-zero coefficients. The features considered include H3K4me1, H3K27ac and H3K27me3 (A separate model involving only the latter two features was built when constructing the enhancer-target network in MCF-7 since H3K4me1 data were unavailable).

In the second step, single-enhancer error terms were first computed. Specifically, an error term is computed to check how much the expression of the TSS in sample $k$ can be explained by considering each feature of each enhancer, i.e., where is the value of feature of enhancer in sample and are the coefficients learned in the first step. These error terms were then combined with genomic distance and cell-line-specific data (i.e. the levels of histone modifications across the enhancer, the TSS and the window between them in sample) to predict the enhancers that regulate a TSS in a particular cell line using a Random Forest model. The parameter values of these second-level models were learned from published ChIA-PET data from K562 and MCF-7 cell lines. A 5-fold cross-validation procedure was used to evaluate the accuracy of the predicted enhancer-target pairs. The model was then applied to those samples without ChIA-PET data.

*Figure S 2-6. Schematic of JEME*

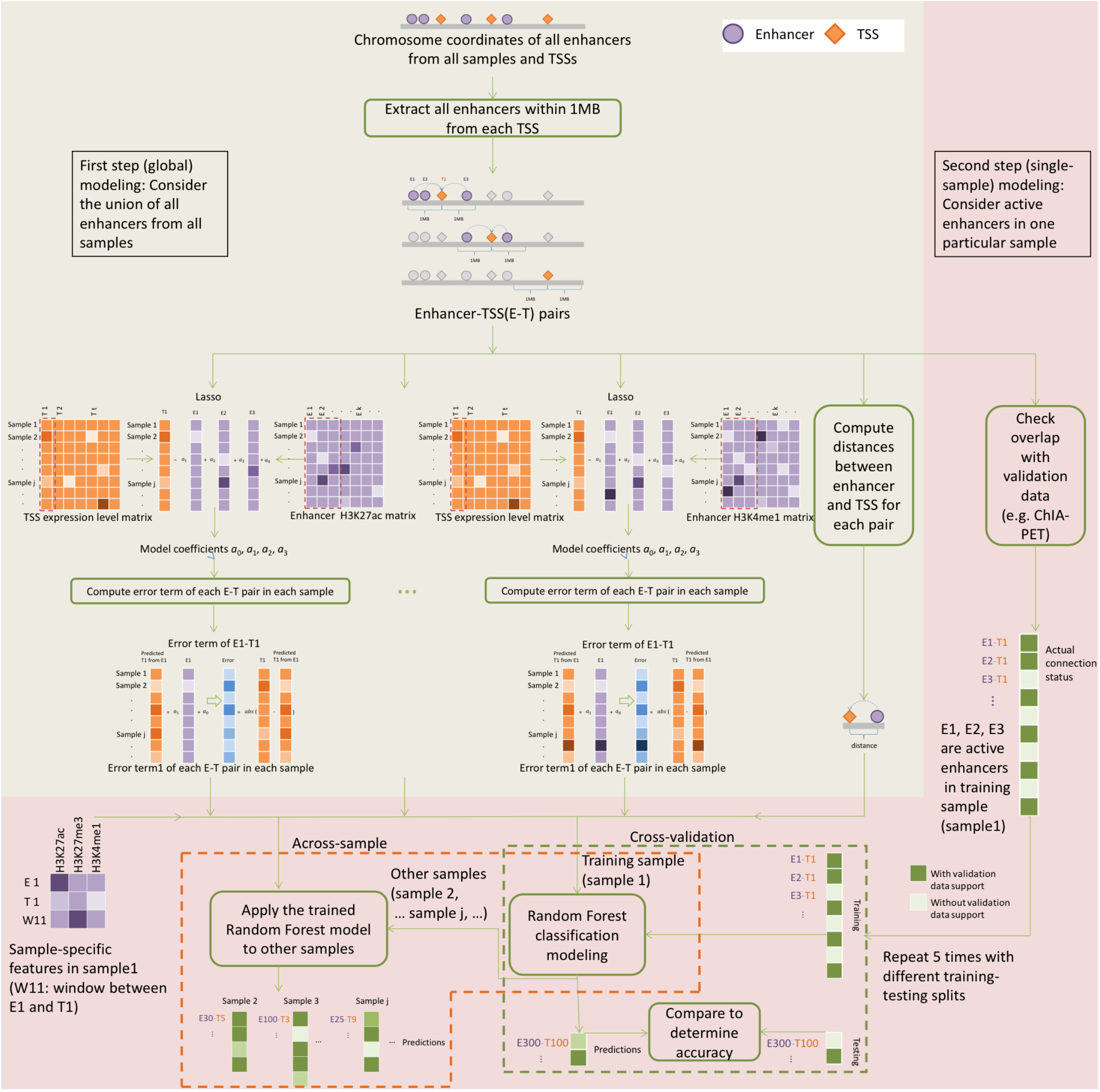

#### 2.2.2 (TL, ‖) Enhancer gene linkage pruning using Hi-C data

Enhancer target predictions are further filtered by using Hi-C data. Contact maps of individual chromosomes (in 5kb bins) for both K562 and GM12878 cell lines were obtained from Rao et al[43]. MCF-7 contact maps (40kb) were obtained from Barutcu et al[44]. An element in a contact map represents the frequency of interactions between genomic loci and for all possible, we used the tool Fit-Hi-C to estimate the statistical significance of the contact frequency based on the coverage of the loci as well as their genomic distance[45] and to keep the interactions with q-value<0.1. We then used the list of significant loci to filter the enhancer-target predictions. Only enhancer-gene pairs in which enhancer and gene respectively belong to a pair of significantly interacting loci are kept for further analysis.

#### 2.2.3 (TL, ‖) Extended gene neighborhood generation

Here we generated the extended gene neighborhoods by combining the coding region with the key non-coding proximal and distal regulatory elements together for a joint mutation burdening quantification. Details of the schematic are given in Figure S 2-7.

There are four important basic elements in our extended gene definition: CDS, TFBS, RBP binding sites, and enhancers. For each gene, based on Gencode V19 annotation we extracted all the TFBS within 2.5kb of the TSS sites of the protein coding transcript, all the eCLIP binding sites of the whole transcript (and upstream 200 bp and downstream 1500 bp), all the linked enhancers, and then merged these annotations together to form the extended gene.

*Figure S 2-7. (Shadow figure for Fig. 2 C) Schematic of extended gene definition*

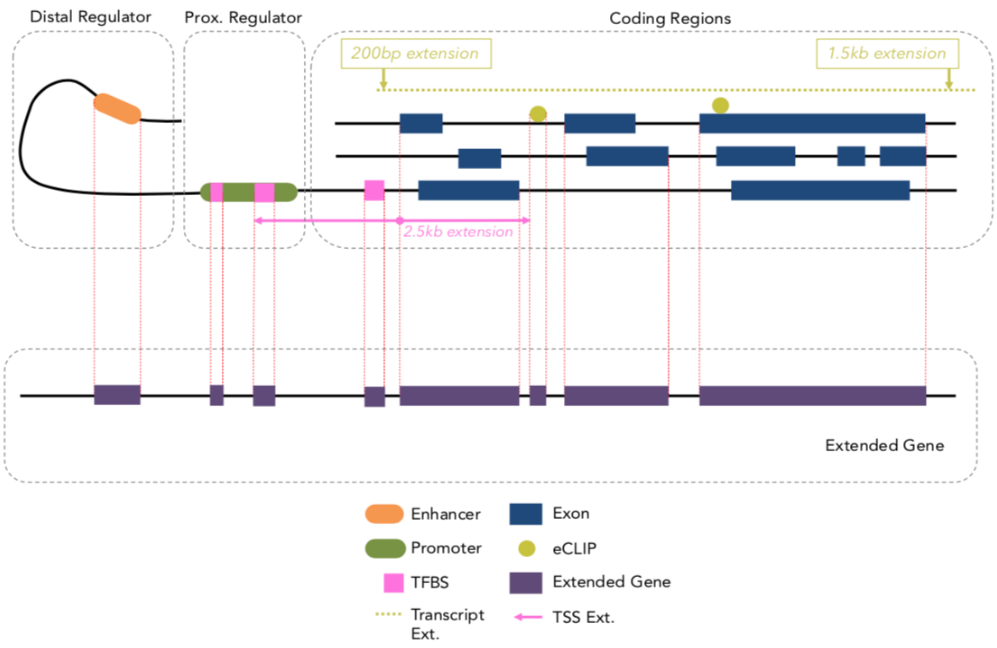

#### 2.3 Construction of regulatory networks

##### 2.3.1 (HL, ‖) Tissue-specific TF networks

TF regulatory network can be directly inferred from the experimental evidence using DNA binding assays or indirectly using imputation based on chromatin status and TF binding motifs. We built TF-gene interaction networks based on uniformly processed ENCODE ChIP-seq assays. Largely, we built TF network in two steps, by defining promoter-based linkages (proximal) and enhancer-based linkages (distal). In the first step, the promoter-based linkages were defined using two methods:

##### 2.3.2 (TL, ‖) TSS-based network (based on proximity of binding to TSS)

From IDR-thresholded peak calls, we first defined TF-gene regulatory network in each cell type by searching for TF to gene linkages based on their proximities to the TSS. We defined TSS regions as upstream and downstream of 2,500bp within TSS based on Gencode v19 annotation.

##### 2.3.3 (TL, ‖) TIP network (based on TF binding signal strength to TSS)

In addition to TSS based TF network, we also used target identification from profiles (TIP) method that quantitatively measures the regulatory relationships between TFs and target genes to define a subset of the full TF-gene network[46]. Both promoter-based linkages (either TSS or TIP) and enhancer target-based linkages were merged into one to build a complete TF-gene network. For more information about enhancer target-based linkages, please refer to section 2.2.

##### 2.3.4 (TL, ‖) Universal TF network

We also provide a more generalized network by reconciling networks from multiple cancer types and applied it in a pan-cancer analysis. We built a universal TF network by merging tissue specific networks across all available ENCODE tissue types. Two versions of universal regulatory networks were constructed; one larger network by concatenating TSS-based network and enhancer-based network and another subnetwork by concatenating TIP-based network and enhancer-based network.

##### 2.3.5 (TL, ‖) Probabilistic and TF/RBP network

For regulatory analysis, we only considered transcription factors (TF), chromatin regulators (CR), and RNA binding proteins (RBP). In total, there are ***978*** TF/CR ChIP-seq profiles and ***159*** RBP eCLIP profiles downloaded from ENCODE DCC as of January 4th, 2017 (https://www.encodeproject.org).

All ChIP-seq and eCLIP peak scores are linearly scaled into range (0,1). The regulatory score between TF peaks and gene promoters were built with "connect_host" commands from RABIT package[47] following an exponential decay model (Figure S 2-8). A regulatory score between RBPs and genes were built through counting eCLIP peaks within gene 3'UTR regions. The following steps were made to construct the network. a) For ChIP-seq data, A regulatory potential score is calculated between each pair of ChIP-seq peak and gene TSS by multiplying the ChIP-seq intensity score with an exponential decay score exp (-A*Distance) of their distance between. The coefficient A is set as log(2)/10K, so that a binding peak 10K bps away from gene TSS will decay by 50%. For each gene TSS, if there are several peaks of a TF nearby, we merged their regulatory potential scores by noisy-or: (b) For eCLIP data, only binding peaks over gene 3'UTR regions were considered for possible regulatory role of transcript stability. For each gene 3'UTR region, if there are several peaks of a RBP, we merged their regulatory potential scores by noisy-or operation. All regulatory potential scores stay within range (0,1).

Figure S 2-8. Regulatory network construction

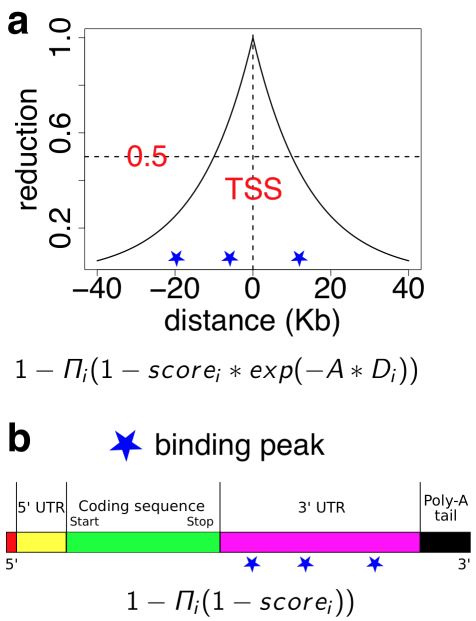

ChIP-seq and eCLIP profiles were excluded from further analysis if the total sum of regulatory scores across all human genes is less than 100. All general TFs including Pol2 and Pol3 were excluded from further analysis. For certain TF, there exist many ChIP-seq profiles profiled in different conditions. We run a hierarchical clustering among all of its ChIP-seq profiles and cut the hierarchical tree at correlation distance of 0.2. Only profiles in the largest cluster are used for further analysis. The final size of regulatory networks constructed are shown in Table S 2-3. For each data type, column "Profile" represents the number of experimental profiles (ChIP-seq or eCLIP) that passed our quality controls. Column "Regulator" represents the number of regulators (TF, CR or RBP) analyzed. Column "Condition" represents the number of experimental conditions included in profiles. Column "Target" represents the total number of human genes profiled as targets of analyzed regulators.

Table S 2-3. Statistics of regulatory networks

| | **Profile** | **Regulator** | **Condition** | **Target** |
|---|---|---|---|---|
| **ChIP-seq** | 762 | 496 | 44 | 21,348 |
| **eCLIP** | 159 | 112 | 2 | 14,593 |

We defined the RBP network based on the RBP-gene interactions from eCLIP peak calls. eCLIP peak scores between RBPs and genes were built through counting eCLIP peaks within gene's 3'UTR regions and then linearly scaled into range (0,1). To provide a strict and lenient RBP network, we use 0.9 and 0.1 as the threshold for interaction cutoffs. RBP networks are summarized in the supplementary datasheet.

##### 2.3.6 (TL, ||) Distal TF-enhancer-gene network

We defined the distal regulatory edge based on our integrative enhancer definition and enhancer target linkages. We used a subset of integrative enhancers have target gene linkage with high confidence (for more info, please refer enhancer target gene linkage section). We overlapped TF binding peaks based on ENCODE ChIP-seq data and defined them as distal regulatory linkage.

#### 2.4 Quality control of the ENCODEC resource

##### 2.4.1 (TL, ∦) QC of the TF and RBP binding sites

In Figure S 2-9, we take the density of the average peak length of RBPs from eCLIP and TF as a comparison of the difference in data quality. The resolution of eCLIP and ChIP-seq differ, though they are thought to have similar binding site lengths. For each RBP and TF, the rare DAF is calculated to be the number of rare variants divided by the number of rare and Figure S 2-9. Also, we show that for the common annotations including the CDS, 3’ UTR, 5’ UTR, 3’ UTR extended, 5’ UTR extended and proximal intron regions, overlap with eCLIP peaks results in a higher Phastcon score[48] than compared to annotations that do not overlap eCLIP peaks.

*Figure S 2-9. QC of TF and RBP binding sites*

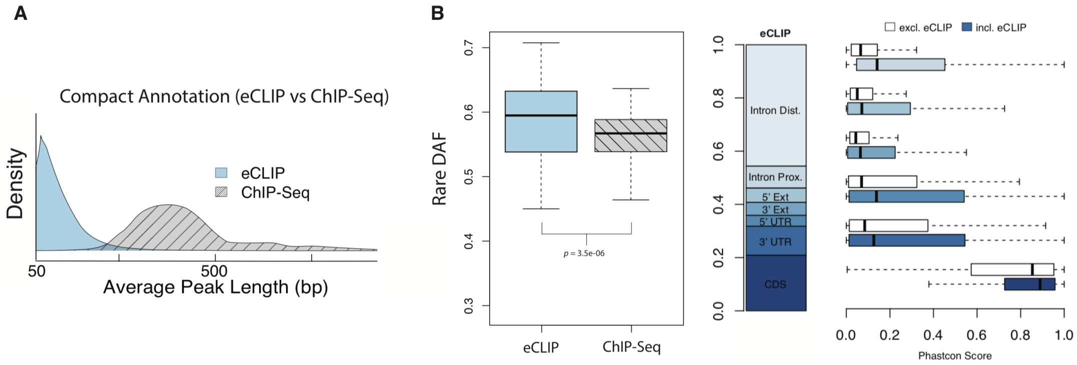

For the eCLIP peaks, we further determine the quality of the data by taking the rare DAF of each RBP and correcting it for its GC content. The green dots in Figure S 2-10 demonstrate the observed rare DAF, and the yellow dot refers to the GC corrected background for comparison. Almost all RBPs demonstrate a higher than genome average rare DAF.

*Figure S 2-10. Rare DAF of the RBP binding sites*

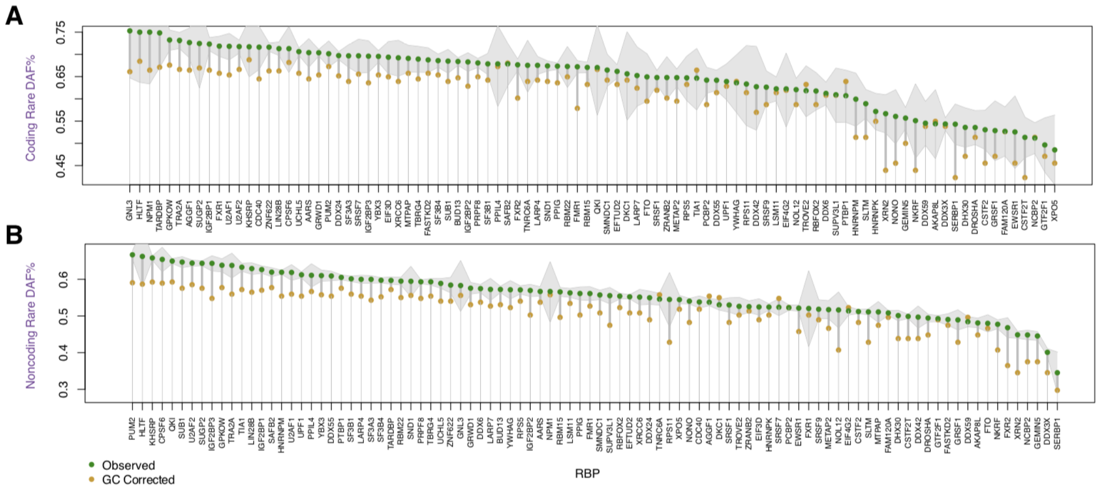

##### 2.4.2 (TL, ∦) QC of the enhancers

Here we took the FANTOM enhancer data set and assess the overlap percentage of our enhancer annotation in each ensemble step. We showed that each ensemble step indeed increases the percentage of overlap between our annotation and the FANTOM enhancer set. The overlap percentage for our annotation is much higher than that of the Roadmap annotation and is also higher than the candidate cis-regulatory element (ccRE) enhancer-like signature (ELS) annotation.

*Figure S 2-11. Overlap of the ENCODEC enhancer with FANTOM*

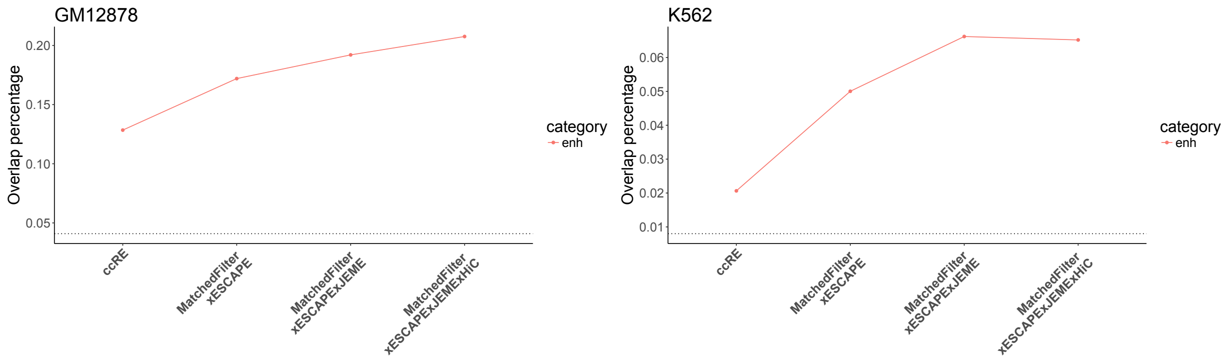

##### 2.4.3 (TL, ∦) QC of the enhancer gene-linkages

Previously, we developed a computational approach JEME to predict enhancer-gene linkages. We have done extensive benchmark against other methods, such as IM-PET, Prestige, and Targetfinder. Details can be found in Cao et al[42].

In this paper, we used a 2-step approach of finding enhancer-target gene linkages. First, we used our previously published JEME algorithm to find the linkages. We then filtered the enhancer-target gene linkages using the significant Hi-C interactions that are found using the method Fit-Hi-C[45]. This is done to make sure that all of our enhancer-target gene linkages have physical interactions between them.

To show how our JEME+Hi-C approach captures enhancer-gene linkages compared to existing linkages, we used published chromHMM[49] derived enhancer-gene as the comparison dataset and GTEx whole blood eQTLs as the benchmark. We found the linkages, which the enhancer has an eQTL that changes the expression of the target gene significantly. After finding all the eQTL supported linkages for chromHMM and JEME+Hi-C, we calculated the fraction of enhancer-gene linkages that has eQTL support for various types of linkages in chromHMM and in JEME+Hi-C. As can be seen in figure below, JEME+Hi-C has higher fraction overlapped with eQTL-gene linkages.

*Figure S 2-12. Overlapping the gene-target linkages with GTEx eQTLs*

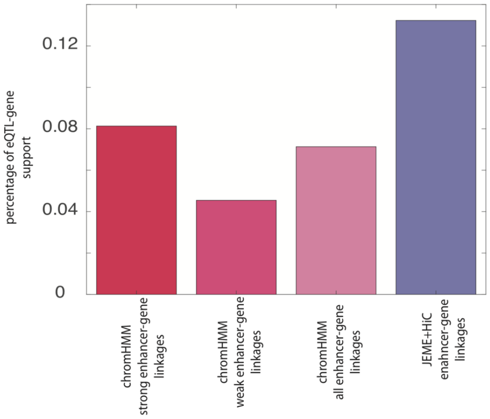

#### 2.4.4 (TL, ∦) QC of networks

##### 2.4.4.1 (TL, ∦) Comparison with Biogrid and String networks

To evaluate the quality of ENCODE transcriptional regulatory networks, we utilized the TRRUST database, which manually curated transcriptional regulations from Pubmed articles[50]. We defined the TRRUST interactions as the standard and tested the fraction of standard interactions that other networks can recapitulate. The ENCODE network can capture a higher fraction of standard interactions than protein physical networks, including Biogrid and String experimental interactions (Figure S 2-13). Moreover, the fraction of standard networks that ENCODE network recapitulated is consistently higher than random. These results supported the higher relevance of ENCODE networks on transcriptional regulation compared to other networks. We also constructed another post-transcriptional network between RBPs and target genes through linking the RBP binding sites on gene 3'UTR regions. To the best of our knowledge, the current study is the first one to study RBP-gene interactions systematically; thus, we are not aware of any previous resources that can provide gold standard regulations for comparison.

*Figure S 2-13. ENCODE networks captured a higher fraction of curated regulations than other networks*

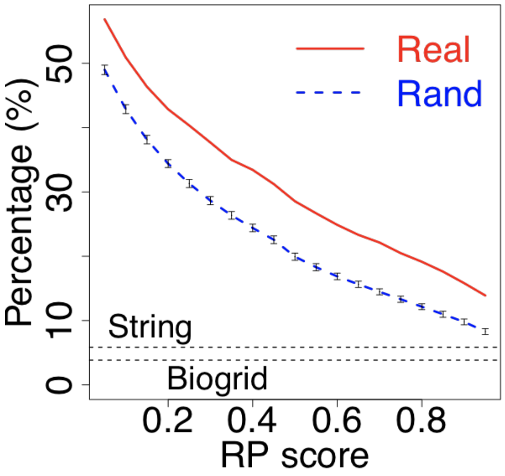

The TRRUST database manually curated 8,412 transcriptional regulatory interactions from Pubmed articles[50]. We computed the fractions of TTRUST interactions that other networks can recapitulate. Since each ENCODE ChIP-seq interaction has a regulatory potential (RP) score, we showed the fractions with different RP thresholds. The random fraction for ENCODE network was estimated through 100 perturbed TTRUST networks using the stub-rewiring method that preserved the gene network degrees[51].

###### 2.4.4.2 (TL, ∦) Comparison with imputed networks

Our new regulatory network edges are derived from ENCODE TF ChIP-seq experiments, and they provide more accurate gene linkages than imputed networks from other genomic features. To demonstrate the superiority of our new network, we have evaluated our experimentally derived ChIP-seq networks with DHS-based imputed networks from previous publications. We have used two types of ChIP-seq networks. The first one is based on proximity to TSS and the second one based on target identification from profiles (TIP) method. For imputed network, we used TF-to-TF network imputed from DNase I hypersensitive footprints as in Neph et al[52]. In addition to Neph et. al. DHS network, we also built our own version of similar DHS network by utilizing the ENCODE DNase-seq dataset. To test the gene linkages, we have utilized ENCODE RNAi based TF knockdown and CRISPR-based TF knockout datasets to test how the target gene linkages defined by various network definition are affected by after KD/KO. Overall, target genes of ENCODE ChIP-seq networks had larger differential expression after knocking down (Figure S 2-14). Moreover, DHS-imputed network derived from ENCODE DNase-seq performed better than the previously published method.

*Figure S 2-14. Evaluation of ENCODEC networks with previously published methods using ENCODE knockdown data*

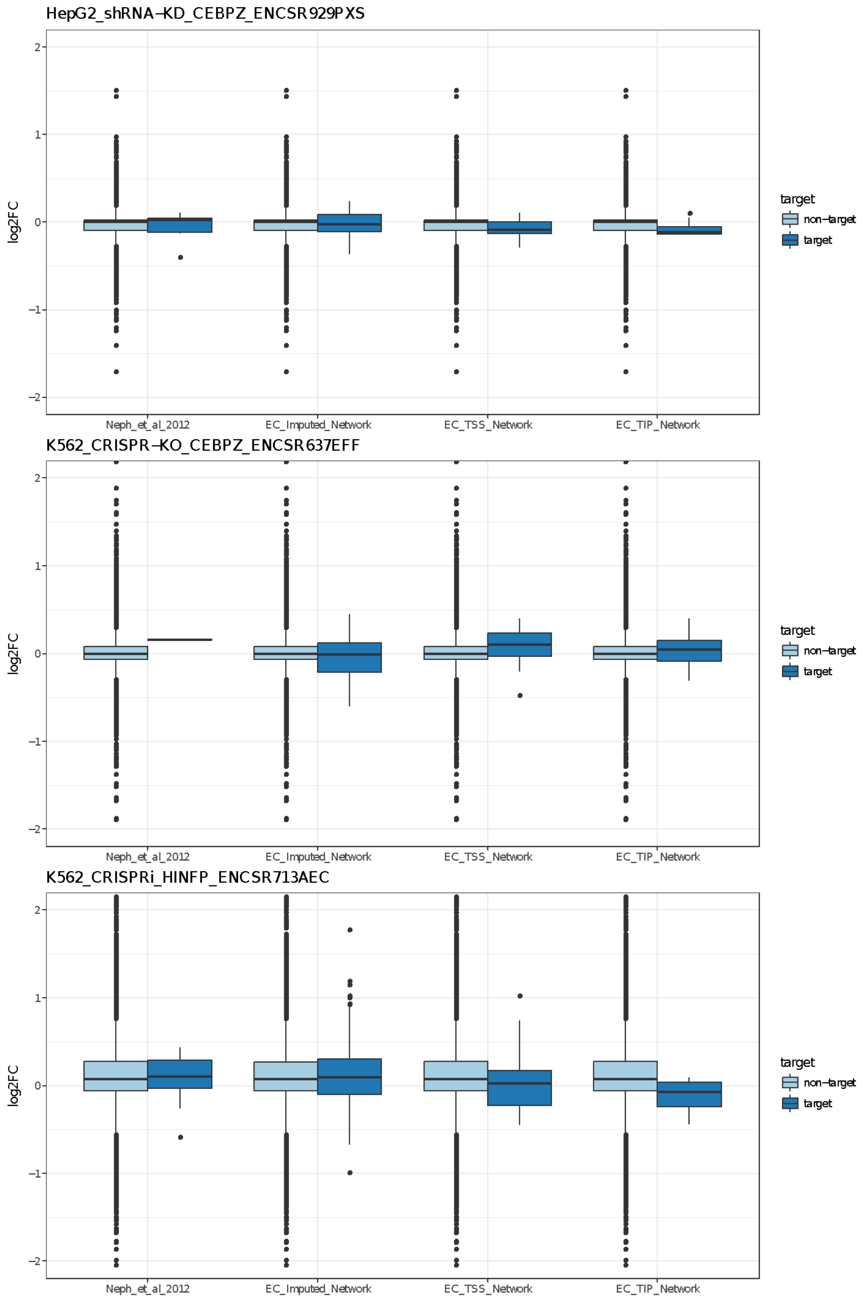

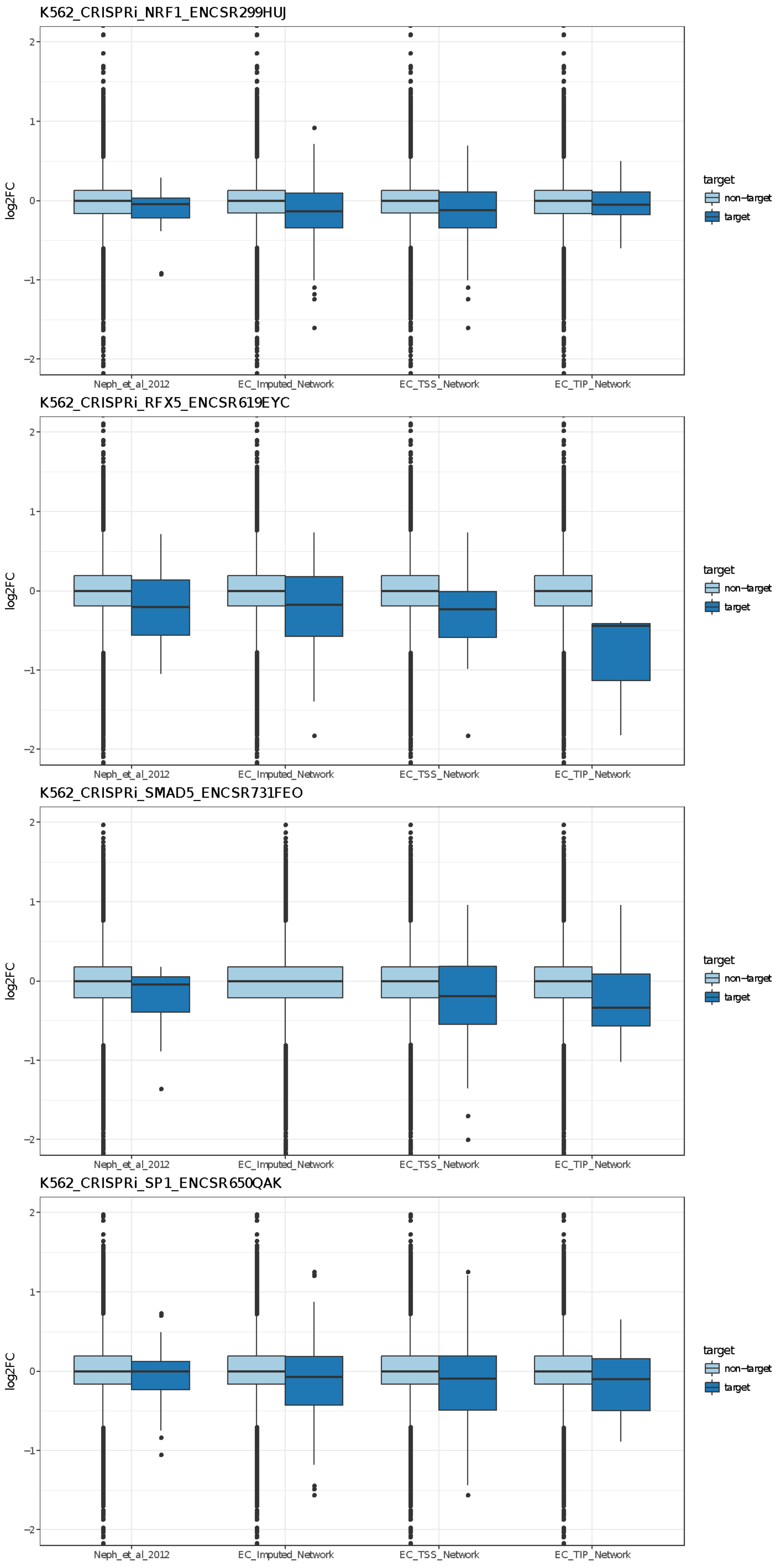

K562_CRISPRi_NRF1_ENCSR299HUJ
K562_CRISPRi_RFX5_ENCSR619EYC
K562_CRISPRi_SMAD5_ENCSR731FEO
K562_CRISPRi_SP1_ENCSR650QAK
log2FC
2
1
0
−1
−2
Neph_et_al_2012
EC_Imputed_Network
EC_TSS_Network
EC_TIP_Network
target
non−target
target

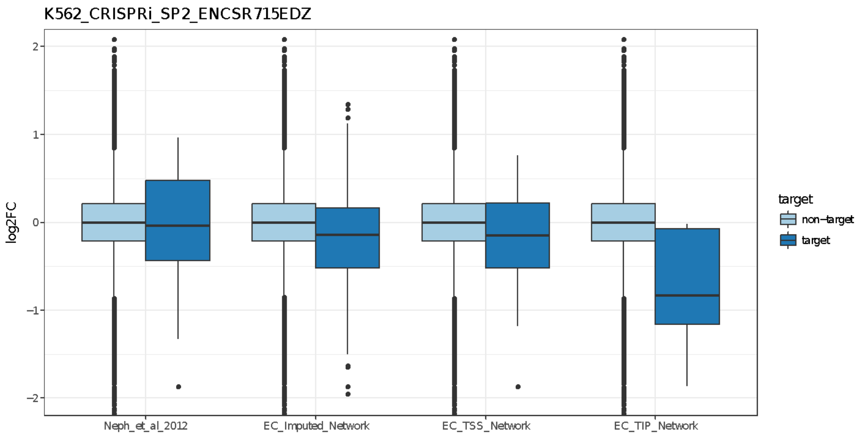
K562_CRISPRi_SP2_ENCSR715EDZ
log2FC
2
1
0
−1
−2
Neph_et_al_2012
EC_Imputed_Network
EC_TSS_Network
EC_TIP_Network
target
non−target
target

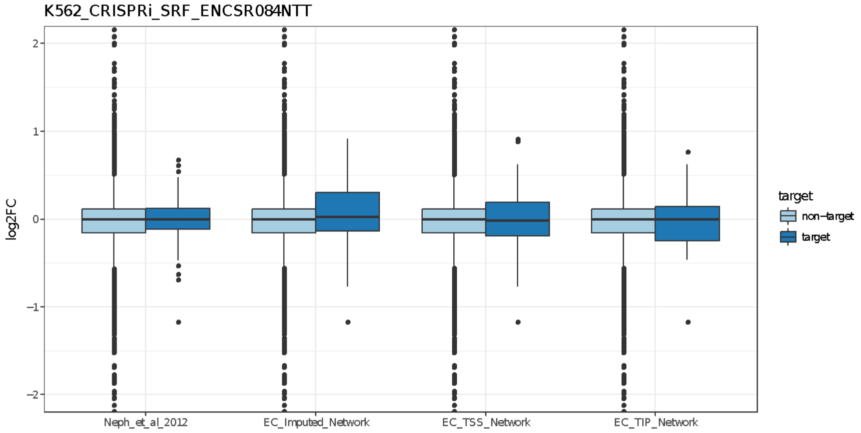
K562_CRISPRi_SRF_ENCSR084NTT
log2FC
2
1
0
−1
−2
Neph_et_al_2012
EC_Imputed_Network
EC_TSS_Network
EC_TIP_Network
target
non−target
target

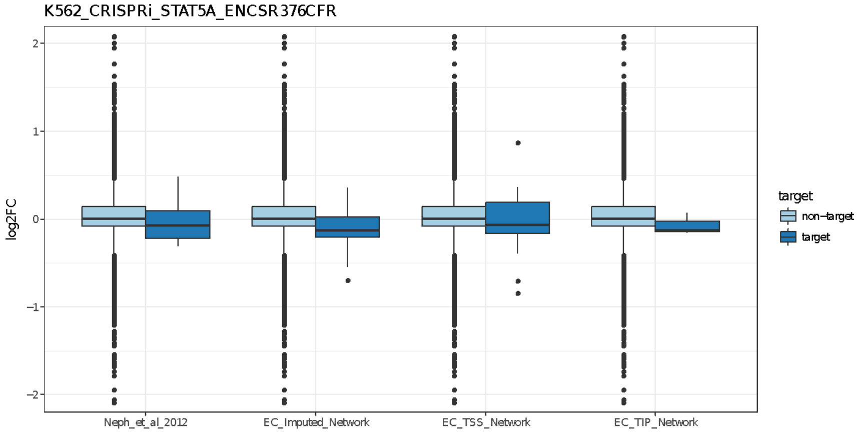
K562_CRISPRi_STAT5A_ENCSR376CFR
log2FC
2
1
0
−1
−2
Neph_et_al_2012
EC_Imputed_Network
EC_TSS_Network
EC_TIP_Network
target
non−target
target

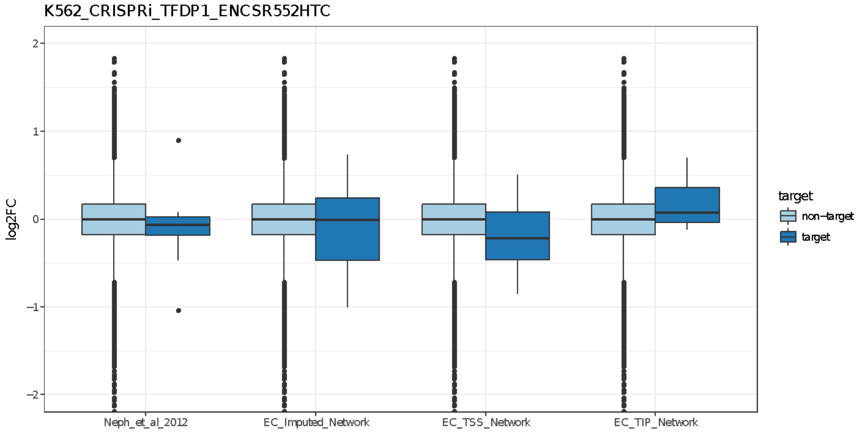
K562_CRISPRi_TFDP1_ENCSR552HTC
log2FC
2
1
0
−1
−2
Neph_et_al_2012
EC_Imputed_Network
EC_TSS_Network
EC_TIP_Network
target
non−target
target

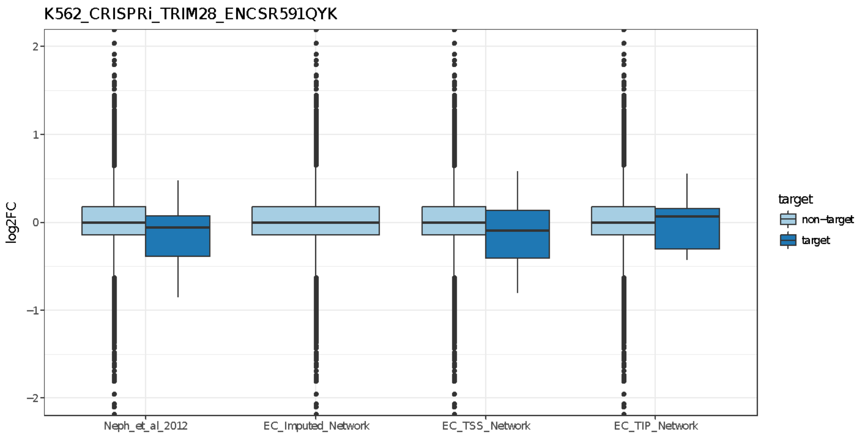
K562_CRISPRi_TRIM28_ENCSR591QYK
log2FC
2
1
0
−1
−2
Neph_et_al_2012
EC_Imputed_Network
EC_TSS_Network
EC_TIP_Network
target
non−target
target

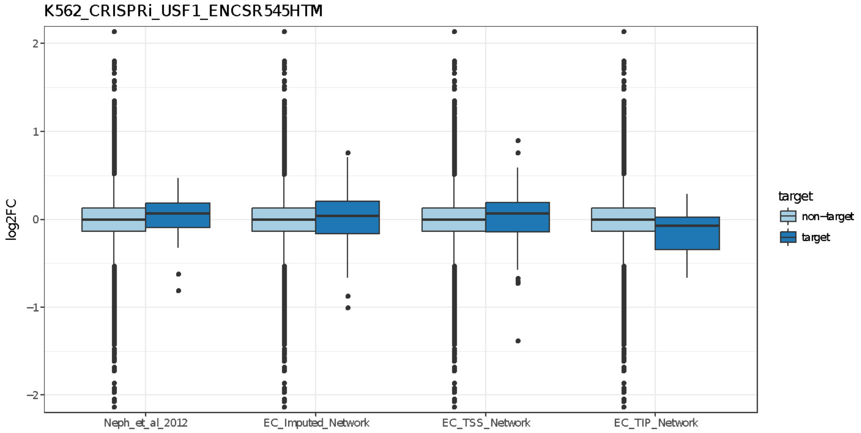
K562_CRISPRi_USF1_ENCSR545HTM
log2FC
2
1
0
−1
−2
Neph_et_al_2012
EC_Imputed_Network
EC_TSS_Network
EC_TIP_Network
target
non−target
target

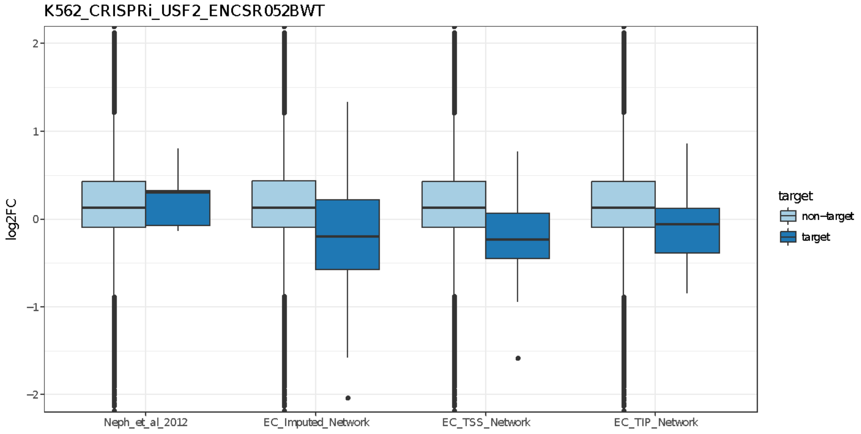
K562_CRISPRi_USF2_ENCSR052BWT
log2FC
2
1
0
−1
−2
Neph_et_al_2012
EC_Imputed_Network
EC_TSS_Network
EC_TIP_Network
target
non−target
target

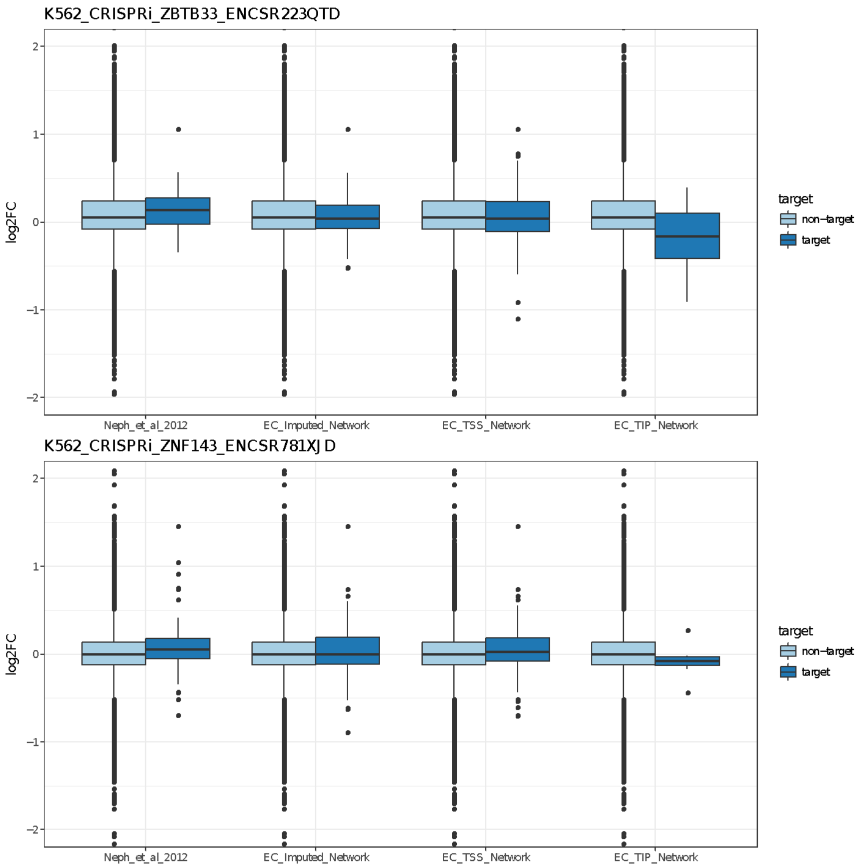

#### 2.5 (TL, ∦) Reconciliation with the main ENCODE encyclopedia

Both promoter and enhancer annotations from EN-CODEC were carefully consolidated with the main ENCODE Encyclopedia resources. The ENCODE Encyclopedia comprises of three levels, two integrative levels of annotations and the ground level raw data. The ground level includes peaks and quantifications produced by uniform processing pipelines for individual data types. The integrative level contains annotations produced by integrating multiple data types. The core of the integrative level is the candidate Registry of candidate Regulatory Elements (ccREs). The registry contains approximately ~1.31M human ccREs and each ccRE has a cell-type non-specific accession number, which then can be browsed from SCREEN (Search Candidate Regulatory Elements by ENCODE, http://screen.encodeproject.org/).

Annotations from EN-CODEC were merged against the Registry of candidate Regulatory Elements (ccREs). We assigned cell type non-specific ccRE accession numbers to ESCAPE and MatchedFilter integrated enhancer annotations when the region had more than 1bp overlap. When there were more than one accession numbers associated with the annotation, we assigned multiple accession numbers to the element. Overall, there was 99% overlap between integrated enhancer annotations and ccREs with each element being mapped to 2.5 ccRE accessions on average.

For cases without an overlap, we assigned special accession numbers EH37EXXXXXXX-C where XXXXXXX are replaced with numbers starting from 0000001. To access the ccRE using accession number, one can use the URL http://screen.encodeproject.org /search/?q={accession}&assembly=hg19# where {accession} is replaced with the ccRE accession number. From SCREEN, one can look up H3K4me3, H3K27ac, CTCF, and DNase Z-scores and signal profiles across all available ENCODE cell types.

#### 2.6 Use the compact and extended annotation to increase statistical power

##### 2.6.1 (TL, ‖) Overall parameterization

We consider a model in which an element is partitioned into a 'core' region of length $l_1$ and a 'peripheral' region of length $l_2$ (total length $L = l_1 + l_2$) Additionally, we let $\mu$ be the BMR across the element, and $\lambda$ be the per-base effect size of functional bases within the element.

If all the sites in the core are functional with probability 1, and those in the periphery are functional with probability 0, the probability[53,54] to observe at least one mutation per patient under the null and alternative hypotheses are respectively are:

$$p_0 = 1 - (1-\mu)^L = 1 - (1-\mu)^{l_1+l_2}$$

$$p_1 = 1 - (1-\lambda\mu)^{l_1}(1-\mu)^{l_2}. \quad (2\text{-}7)$$

For a sample size $N$ and desired significance level $\alpha$, the number of samples with at least 1 mutation is represented by $x$. Suppose we need to test $K$ such regions. The power $Q$ is determined by first calculating $X_0 = min\,\{X|B(x > X; p_0, N) < \alpha/K\}$, to determine $Q = B(x \geq X_0|p_1, N)$, where $B(x > X_0; p, N)$ denotes the probability that $x$ is greater than $X_0$ under a Binomial distribution with parameters $p$ and $N$.

##### 2.6.2 Compact annotation can increase power

###### 2.6.2.1 (TL, ‖) Increasing power by removing noise sites per individual annotation element

In a case where compactification removes only non-functional sites, power will be increased by creating a compact annotation. This can be shown by noting that the per patient probabilities in the condensed model are:

$$p_0^{'} = 1 - (1-\mu)^{l_1+l_2^{'}}$$

$$p_1^{'} = 1 - (1-\lambda\mu)^{l_1}(1-\mu)^{l_2^{'}} \quad (2\text{-}8)$$

where $l_2^{'}$ is the length of the compactified peripheral region ($l_2' < l_2$). The differences in binomial means before and after compactification are respectively: $p_1 - p_0 = (1-\mu)^{l_2}[(1-\mu)^{l_1} - (1-\lambda\mu)^{l_1}] = \delta$ and $p_1^{'} - p_0^{'} = (1-\mu)^{l_2^{'}}[(1-\mu)^{l_1} - (1-\lambda\mu)^{l_1}] = \delta'$. Thus, we have

$$\frac{\delta^{'}}{\delta} = (1-\mu)^{l_2^{'}-l_2} > 1 \quad (2\text{-}9)$$

Hence, the difference in means always increases as we remove non-functional sites ($l_2' < l_2$), while the variance of each binomial decreases since $p_0' < p_0$, $p_1' < p_1$, and $p_0, p_1 < 0.5$; hence the power following compactification increases. We show how power decreases with increasing noise level in the annotation in Figure S 2-15.

*Figure S 2-15. Power simulation of noise level within annotation*

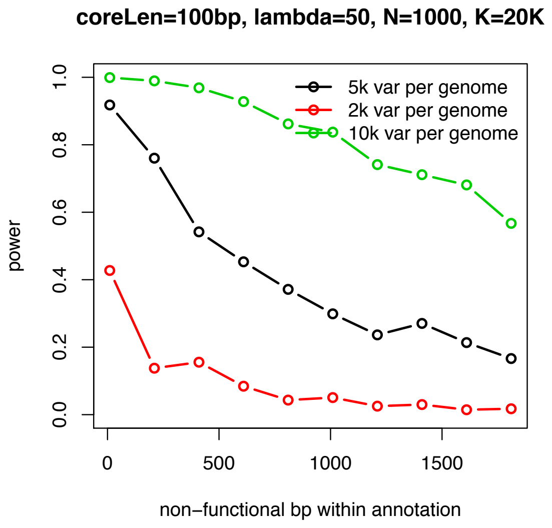

###### 2.6.2.2 (TL, Ⅱ) Increasing power by removing false positives

The number of tests *K* will obviously affect the multiple test burden. A smaller *K* will increase power, which is shown in the following figure.

*Figure S 2-16. Power simulation of false positives effect*

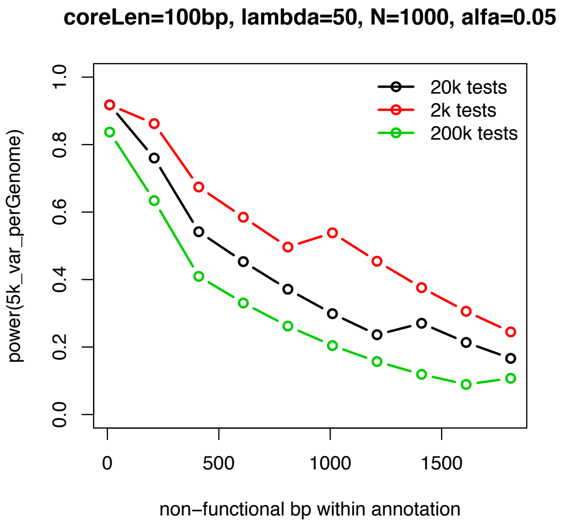

###### 2.6.2.3 (TL, Ⅱ) A real example of importance of compact annotations

We provided one example to explain the motivation of our compact and extended gene annotations and why we feel our assumptions for the power analysis is reasonable. Conventionally, researchers will test the up to 2.5K promoter region for mutation hotspots.

However, we can trim the test set to a core set of the promoter region where many TFs bind, which perfectly correlates with the mutation hotspots (red block) for this well-known driver site (blue line for pan-cancer and green line for liver cancer).

*Figure S 2-17. Exmaple of benefit from compact annotation*

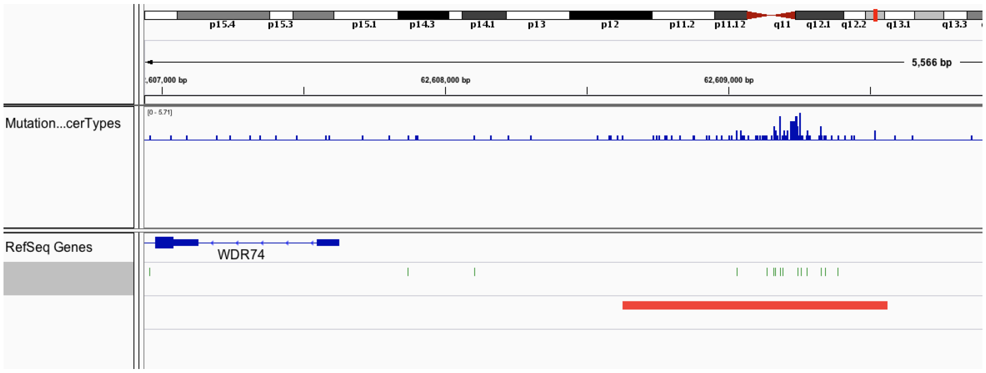

##### 2.6.3 (TL, ‖) Motivation of the extended gene by adding more functional sites

When the number of noise base pairs are fixed, $p_1 - p_0 = (1-\mu)^{l_2}[(1-\mu)^{l_1} - (1-\lambda\mu)^{l_1}] = \delta$ will increase by adding more functional points. This will induce an increase in power to detect highly mutated regions.

*Figure S 2-18. Power simulation of functional site length*

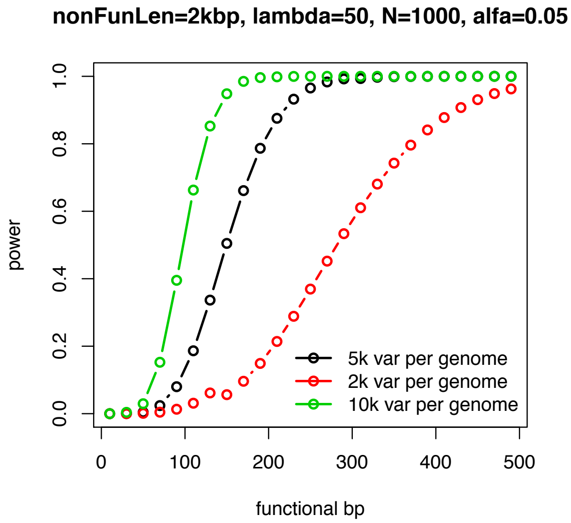

##### 2.6.4 (TL, ‖) Pros and Cons to using the extended gene region as a whole test unit

We now introduce a probability for each base being functional in the core (more functional region) and periphery (noisier region), represented by the parameters $\rho_1$ and $\rho_2$, which can alternatively be treated as densities. Then, for the combined model (which may represent an extended gene, or a single element) we have:

$$p_0 = 1 - (1-\mu)^{l_1+l_2} = 1 - (1-\mu)^L$$

$$p_1 = 1 - (1 - \lambda\mu)^{\rho_1 l_1 + \rho_2 l_2}(1 - \mu)^{(1-\rho_1)l_1 + (1-\rho_2)l_2} \tag{2-10}$$

and for the test on one region model:

$$p_0' = 1 - (1 - \mu)^{l_1}$$

$$p_1' = 1 - (1 - \lambda\mu)^{\rho_1 l_1}(1 - \mu)^{(1-\rho_1)l_1}. \tag{2-11}$$

In this case, for fixed $\lambda, \mu, l_1, l_2$, whether or not the joint test can increase the statistical power depends on the relationship among $\rho_1$, $\rho_2$, and $\lambda$. Specifically, we require that:

$$\rho_2 > min\{B(x > X_0'; p_1', N) \geq B(x > X_0; p_1, N)\}, \tag{2-12}$$

where primes denote quantities calculated in the condensed model. We introduce the following example to discuss the pros and cons in a quantitative way. When the functional site fraction in the peripheral annotation is high (after the first three points in the black line), the combined test will give a power because basically, the combined test is almost adding pure noise to the core region. However, when the quality of the "peripheral" annotation improves (after the 3rd point on the black line), the combined test will start to increase power.

*Figure S 2-19. Power simulation with combined test*

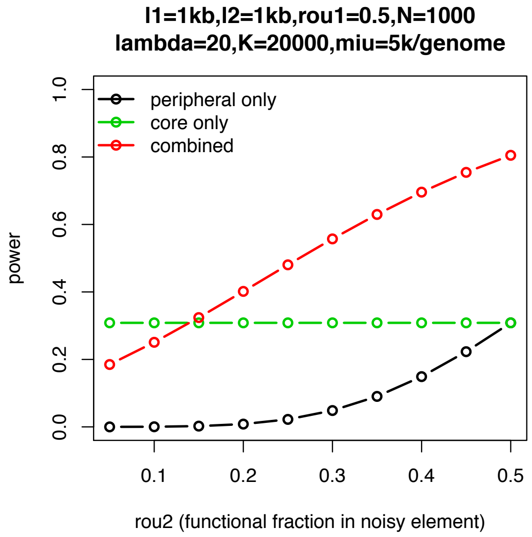

We also show in another example the range of values of $(\rho_1, \rho_2)$ for which the power increases with compactification (red crosses) with other parameters fixed ($l_1 = l_2 = 100,\ \mu = 3e - 6,$ $\lambda = 10,\ \alpha = 0.001,\ N = 600$). We note that $\rho_2$ is necessarily greater than $\rho_1$, since functional sites must be enriched in the core if compactification is to increase power; the amount of enrichment required is determined by the remaining parameters of the model.

*Figure S 2-20. Power simulation with combined test under different situations (blue dots indicates combined test is better)*

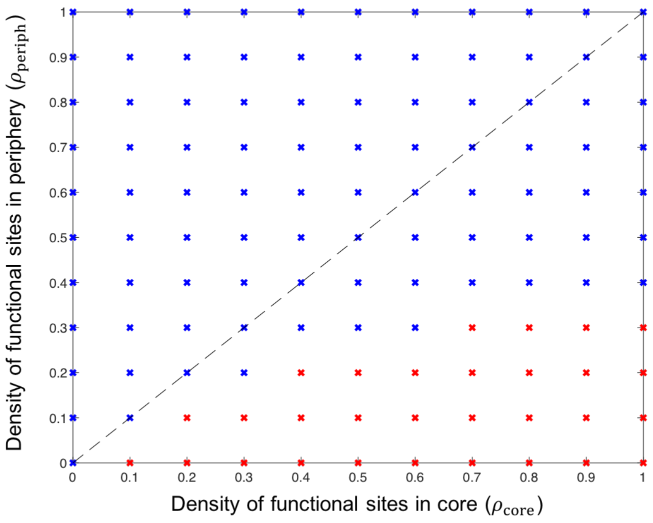

#### 2.6.5 Discussion of "optimal" test combination

We briefly consider a number of refinements to the above analysis by giving two more examples. First, as shown in Figure S 2-21, in principle if we know the exact distribution of the core and noisy regions, we can design an "optimal" test to maximize statistical power. In this case, when the density of functional sites in the periphery is non zero, the optimal compactification (red dotted line) can be slightly larger than the core region (black dotted line, parameters are: $l_1 = l_2 = 50,\ \mu = 3e - 4,\ \lambda = 4,\ \alpha = 0.1,\ N = 600,\ \rho_1 = 0.5, \rho_2 = 0.05$.). Hence, some optimal annotation scheme the predicted core can be slightly larger than the actual core.

*Figure S 2-21. power changes when selecting different test units*

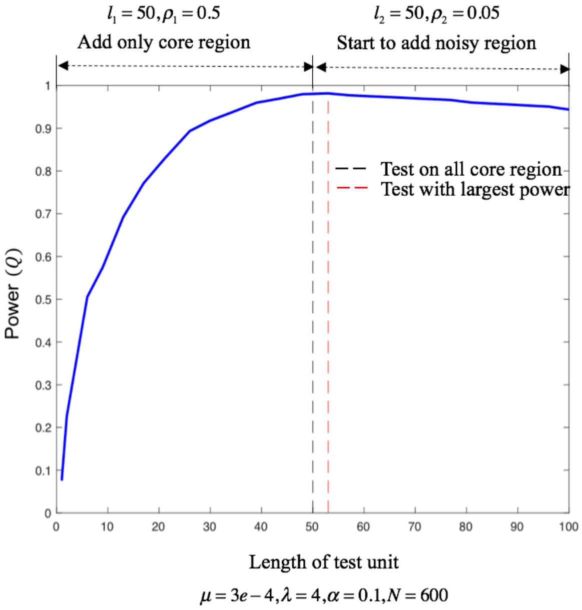

Second, we show that additional power can be achieved by using a weighted test where a weight is assigned to each base matching its probability of functionality, i.e. using the test statistic:

$$S_j = \sum_{i \in core} B_{ij} + w b_{ij}, \tag{2-13}$$

where $b_{ij}$ is a binary indicator which is 1 if a mutation is present at base $i$ in individual $j$, and 0 otherwise, and $w$ determines the relative weight of mutations observed in the periphery relative to the core. Since $b_{ij}$ is a Bernoulli variable, $S_j$ is distributed as a weighted sum of Bernoulli variables; since this distribution does not have a closed form representation, we use simulations to estimate the distribution of $S_j$ under null and alternative hypothesis in a core-periphery model as above, and evaluate the power that can be achieved while varying $w$ between 0 and 1. The figure below shows that the best power is achieved for an intermediate $w$. Further, the dotted lines show the power achieved by a binomial test as above, including mutations from across the whole element or the core only (lower and upper dotted black lines respectively) and the best intermediate binomial model (red dotted line). In general, the full/partial condensed binomial models above can be seen as particular weighting schemes where the weights are constrained to take binary values and the max rather than sum is used the form the per-patient statistic. Potentially then, it may be possible to increase power further in mutation burden testing by using weighted annotations and a summed statistic, although the figure suggests that the power achieved by the best binomial test is close to that of the best weighted summation test (although in general this will depend on the underlying model parameters). Parameters used are: $l_1 = l_2 = 50,\ \mu = 3e-4,\ \lambda = 2,\ \alpha = 0.1,\ N = 600,\ \rho_1 = 0.5, \rho_2 = 0.06$, and 200 trials are used to generate the empirical distributions for the power calculation.

*Figure S 2-22. power simulation using weighted test*

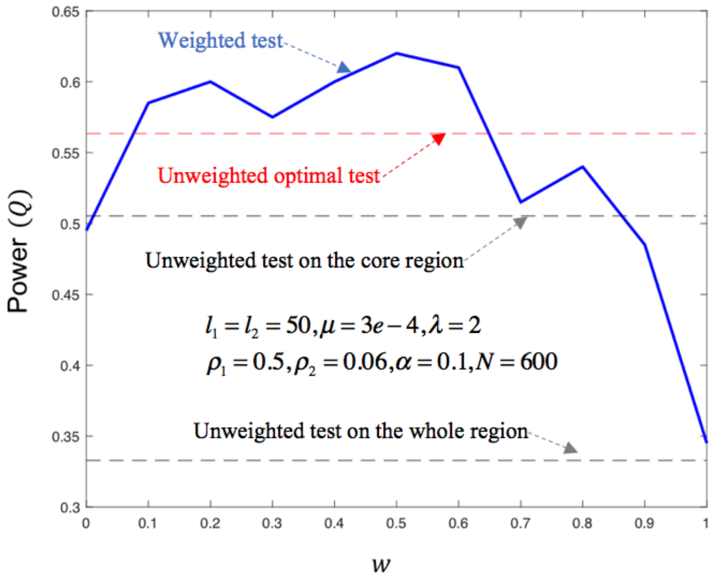

### 3 More details about “Leveraging ENCODE networks to prioritize regulators”

#### 3.1 (TL, ‖) TF/RBP joint network hierarchy

TF network was derived based on TIP merged network using FDR cutoff of 0.1. RBP network was based on 3’UTR based eCLIP merged network. For more information about constructing each network, please refer to the section 2.3. After merging TF and RBP networks, a subset of regulatory edges targeting other regulators were subjected for downstream analysis. Hierarchy of the joint TF and RBP network was determined using HierNet algorithm[55] using 3 layers with default modeling parameters. Each regulator node was colored based on the target expression correlation t-value.

#### 3.2 (TL, ǂ) Expression pattern of TF and RBPs across cancer types.

To investigate the expression patterns of transcription factors across cancer types, we collected the gene expression levels of all TFs and RBPs from TCGA portal for 15 different cancer types. We first performed a data processing step on TCGA expression levels (presented at the next section) for a fair comparison and then plotted the changing expression levels for these TFs (Figure S 3-1 left panel) and RBPs (Figure S 3-1 right panel) across cancer types. As can be seen from the heatmaps, RBPs overall show higher expression levels in these cancer types compared to TFs with a few exceptions. On the other hand, there is more diversity in TF expression levels across different cancer types compared to RBP expression levels.

*Figure S 3-1. Heatmaps of TF and RBP Expressions*

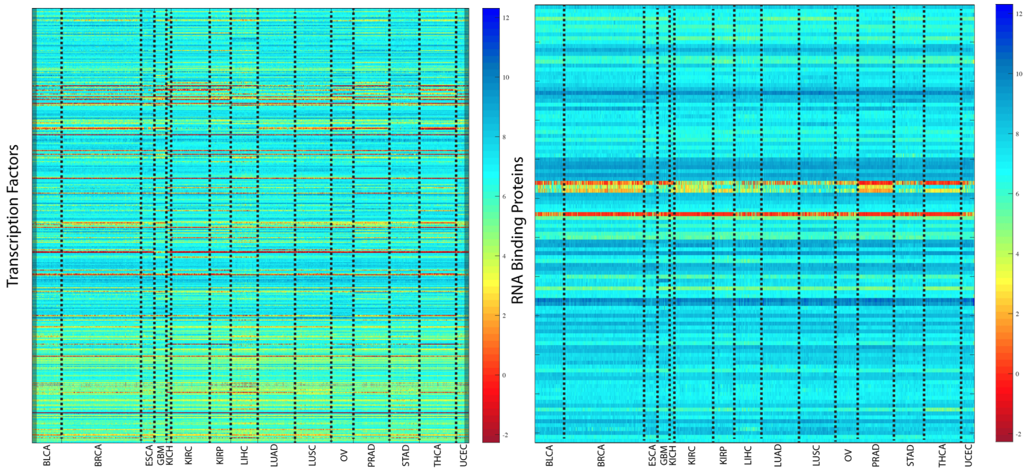

#### 3.3 (TL, ǂ) TCGA expression data processing

All TCGA expression, methylation and mutation data were downloaded from GDAC firehose (http://gdac.broadinstitute.org) with data version of 2016_01_28. For cancer types with normal control samples profiled, the expression values of each gene are subtracted with the average value of all normal controls. For cancer types without any normal samples profiled, the expression profile of each gene is transformed to zero mean and unit deviation. The DNA methylation values are also normalized in the same way as RNA-seq data, according to the availability of normal control samples in each cancer type. For copy number alteration (CNA),

GDAC firehose does not provide standardized data, therefore we downloaded the data matrix from cBioportal with data version of 2016_10_20 (http://www.cbioportal.org). A schematic of this is shown below in Figure S 3-2.

*Figure S 3-2. Schematics of RNA-seq data processing*

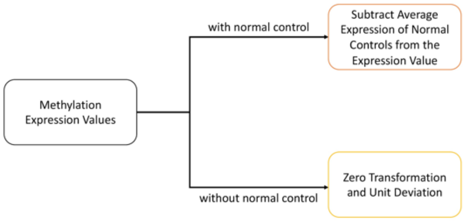

#### 3.4 (TL, ǁ) Identify TF & RBP regulators from cancer genomics data

To systematically search for TFs that drive tumor-specific gene expression patterns, we used a previously developed integration framework RABIT[47] (Regression Analysis with Background InTegration, http://rabit.dfci.harvard.edu). In the RABIT framework, for a given TF ChIP-seq binding profile, candidate target genes are identified by weighting the number of binding sites by their distance to the transcription start site (TSS) of each gene. For a given eCLIP RBP binding profile, candidate genes are identified through searching the binding sites within the gene 3'UTR regions. RABIT uses three steps to identify TFs (or RBPs) that drive tumor-specific gene expression patterns at both the individual tumor level and the whole cancer type level. In Step one, RABIT screens for TFs that significantly affect the gene expression patterns in each tumor and selects the most relevant ChIP-seq (or eCLIP) profile if multiple profiles exist for the same regulator. In Step two, RABIT further selected a subset of TFs among those screened in Step one to achieve an optimized model error. In Step three, RABIT investigates how well the public ChIP-seq profiles can capture the active TF targets in each cancer type and clean up insignificant TFs. The final output of RABIT framework is a set of TFs or RBPs that shape the tumor-specific expression patterns at individual tumor level in each cancer type.

Based on ENCODE ChIP-seq data and TCGA profiles, we applied RABIT framework to identify TFs whose target genes are differentially regulated in cancer. The fractions of patients with TF targets differentially regulated are shown in Figure S 3-3. Only TFs with targets differentially regulated in over 40% patients in at least two cancer types are included. We further extracted those TFs with stronger signals. Except the well-known MYC targets showing consistent up-regulation pattern across multiple cancer types, we also found novel TFs such as ZNF687 to be strongly up-regulated in breast and prostate. Also, the breast tumors were further classified into subtypes according to PAM50 classification and ER status to show the scores predicted by RABIT for each subtype by boxplots. We further checked in each TCGA cancer type, the fractions of patients detected with different types of ZNF687 (Figure S 3-4).

*Figure S 3-3. Heatmap of TF regulator activities in cancer*

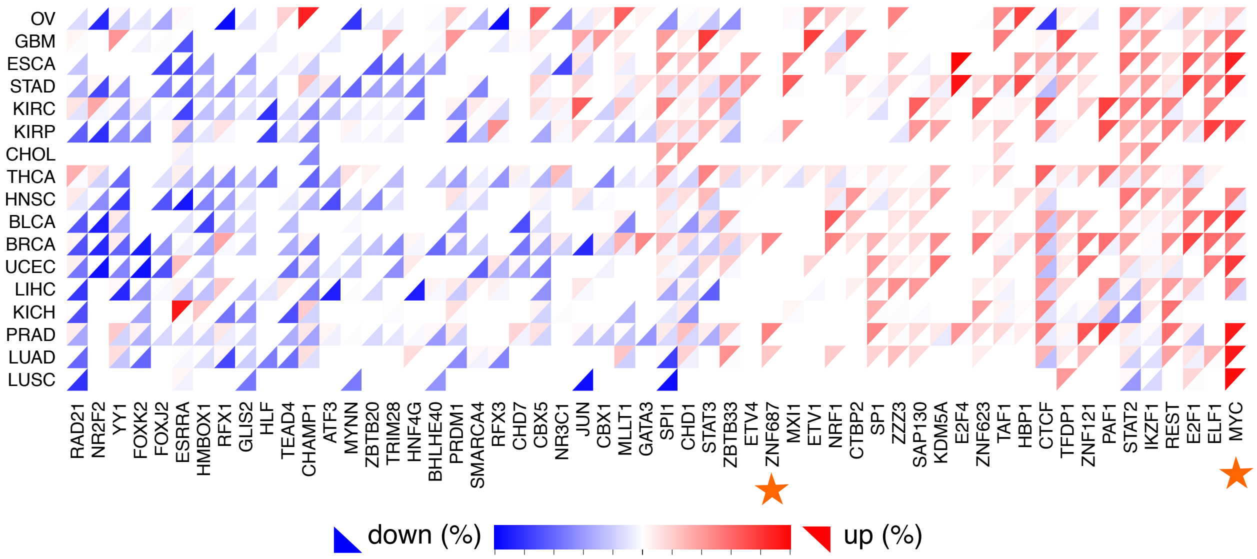

*Figure S 3-4. Potential role of ZNF687 in cancer*

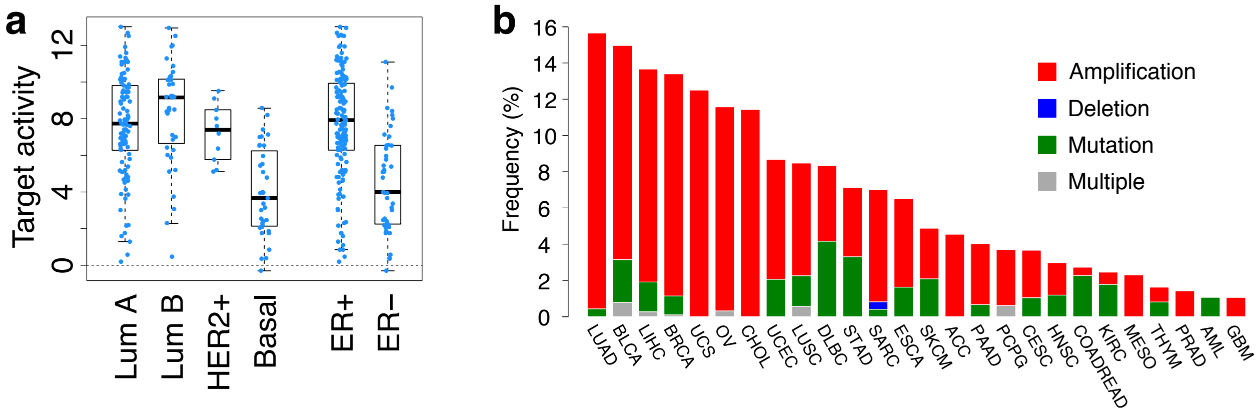

SUB1 was also predicted to be significantly associated with expression changes in multiple tumor types. Here we have listed the full predictions in all cancer types for SUB1. In each cancer type, the association between SUB1 expression and SUB1 regulatory activity predicted by RABIT was tested through t-test in linear regression. Only significant associations above FDR threshold 0.05 are shown in Table S 3-1.

For the SUB1 survival plot in Fig. 2, the association between SUB1 regulatory activity computed for each TCGA tumor for each cancer type studied in TCGA and overall survival was tested through two-sided Ward test in Cox-PH regression. In the KM-plot, all patients with SUB1 regulatory activity larger than 2 were categorized as high and the rest were categorized as low.

*Table S 3-1. Correlation between SUB1 expression and target activity*

| Cancer | Coef | Stderr | *t*-value | *p*-value |
|---|---|---|---|---|
| THCA | 4.79 | 0.46 | 10.46 | 9.03E-23 |
| OV | 4.47 | 0.61 | 7.37 | 1.53E-12 |

| | | | | |
|---|---|---|---|---|
| LUAD | 2.87 | 0.46 | 6.22 | 3.25E-09 |
| PRAD | 2.9 | 0.48 | 6.02 | 4.56E-09 |
| HNSC | 2.61 | 0.46 | 5.72 | 2.73E-08 |
| KIRP | 3.6 | 0.63 | 5.73 | 5.66E-08 |
| GBM | 2.93 | 0.54 | 5.47 | 2.60E-07 |
| LIHC | 3.22 | 0.64 | 5.02 | 1.18E-06 |
| BLCA | 3.21 | 0.66 | 4.83 | 3.91E-06 |
| LUSC | 2.8 | 0.66 | 4.25 | 6.39E-05 |
| KIRC | 1.91 | 0.5 | 3.83 | 1.62E-04 |
| STAD | 2.16 | 0.57 | 3.76 | 2.14E-04 |
| ESCA | 1.67 | 0.54 | 3.09 | 2.34E-03 |
| UCEC | 1.47 | 0.52 | 2.84 | 5.28E-03 |
| KICH | 1.64 | 0.58 | 2.8 | 6.71E-03 |

#### 3.5 Experimental details of the knockdowns in HepG2

##### 3.5.1 (TL, ‖) Transient Knock-down MYC and SUB1 in HepG2 cell line

To detect predicted target genes of MYC and SUB1, shRNA plasmids containing 4 targets sites of each gene were used to transfected to HepG2 cell using Lipofectamine$^{TM}$ 3000 following the manufacturer's instructions (Invitrogen). Target sites for each gene are listed in Table S 3-2.

*Table S 3-2. shRNA plasmids target sites*

| Gene Symbol | Clone ID | Target Seq | Vector | Matching Transcripts |
|---|---|---|---|---|
| **SUB1** | TRCN0000014968 | GAACAGATTTCTGACATTGAT | pLKO.1 | NM_006713.3 |
| **SUB1** | TRCN0000014969 | ACATTGATGATGCAGTAAGAA | pLKO.1 | NM_006713.3 |
| **SUB1** | TRCN0000014970 | GTACGTTAGTGTTCGCGATTT | pLKO.1 | NM_006713.3 |
| **SUB1** | TRCN0000014971 | AGCCAGCTGAAGGAACAGATT | pLKO.1 | NM_006713.3 |
| **MYC** | TRCN0000039640 | CAGTTGAAACACAAACTTGAA | pLKO.1 | NM_002467.4 |
| **MYC** | TRCN0000039641 | CAGGAACTATGACCTCGACTA | pLKO.1 | NM_002467.4 |
| **MYC** | TRCN0000039642 | CCTGAGACAGATCAGCAACAA | pLKO.1 | NM_002467.4 |
| **MYC** | TRCN0000174055 | CCTGAGACAGATCAGCAACAA | pLKO.1 | NM_002467.4 |

Briefly, 0.12 M HepG2 cells were seeded in each well of one 24-well plates 24 hours before transfection. 500 ng plasmids containing either single shRNA or 4 shRNA plasmids as pool were mixed with 0.75 uL Lipofectamine$^{TM}$ 3000 in Opti-MEM I medium (Invitrogen) and loaded to HepG2 cells in each well. Blank plasmids without shRNA target sequence was used as control. To improve transfection efficiency, 2 ug/mL puromycin was used to select successful transfected cells. 72 hours after transfection, total RNA was extracted using RNeasy Mini Kit (Qiagen) and followed by cDNA generation using SuperScript III (Invitrogen). Knockdown efficiency and target gene expression level were quantified and compared to BACTIN by qPCR using KAPA SYBR® FAST qPCR Master Mix (2X) Kit (Sigma). The qPCR primers were listed in Table S 3-3.

*Table S 3-3. qPCR primers*

| qPCR primers | 5'-3' | qPCR primers | 5'-3' |
|---|---|---|---|
| **qMYC-1F** | GGCTCCTGGCAAAAGGTCA | qBIRC5-1F | AAGAACTGGCCCTTCTTGGA |
| **qMYC-1R** | CTGCGTAGTTGTGCTGATGT | qBIRC5-1R | CAACCGGACGAA TGCTTTT |
| **qSUB1-1F** | GGTGAGACTTCGAGAGCCCT | qUNG-1F | CCCCACACCAAGTCTTCACC |
| **qSUB1-1R** | GCGAACACTAACGTACCTCATTT | qUNG-1R | TTGAACACTAAAGCAGAGCCC |
| **qMCM7-1F** | CCTACCAGCCGATCCAGTCT | qAURKB-1F | CAGTGGGACACCCGACATC |
| **qMCM7-1R** | CCTCCTGAGCGGTTGGTTT | qAURKB-1R | GTACACGTTTCCAAACTTGCC |
| **qPFAS-1F** | CCCAGTCCTTCACTTCTATGTTC | qPLK1-1F | CACCAGCACGTCGTAGGATTC |
| **qPFAS-1R** | GTAGCACAGTTCAGTCTCGAC | qPLK1-1R | CCGTAGGTAGTATCGGGCCTC |
| **qRPA2-1F** | GCACCTTCTCAAGCCGAAAAG | qMCM2-1F | ATGGCGGAATCATCGGAATCC |

##### 3.5.2 (TL, ǁ) MYC and SUB1 dual knockdown by lentivirus

To make stable MYC and SUB1 dual knock-down HepG2 cells, PLKO-eGFP-shMYC and PLKO-mCherry-shSUB1 plasmids were generated by replacing puromycin expression cassette with eGFP and mCherry into BamH1 and Kpn1 restriction enzyme sites of PLKO.1 backbone plasmids which targeted MYC and SUB1. shRNA lentiviral stocks were prepared by transient co-transfection of 293T cells with second generation viral packaging vectors (Cat. No. TLP5912, Thermo scientific) as well as PLKO-GFP-shMYC, PLKO-mCherry-shSUB1, or PLKO-eGFP/mCherry empty vectors (control). Three days post-transfection, virus-containing supernatant was collected and filtered through a 0.45 µm filter and supplemented with 8 mg/ml polybrene. HepG2 cells were seeded in 6-well plates one day before transduction, transduced with filtered PLKO-mCherry-shSUB1 virus-containing supernatants and then incubated overnight. Two days after transduction, the successful lentivirus integrated cells were sorted by mCherry signal using BD FACS Aria SORP high-performance cell sorter. mCherry-positive cells were cultured for 2 weeks to generate stable SUB1 knockdown HepG2 cells. Stable SUB1 knockdown HepG2 cells were then transduced with PLKO-eGFP-shMYC virus to generate stable MYC/SUB1 dual knockdown HepG2 cells by dual-color cell-sorting. PLKO-eGFP/mCherry empty virus were used to generated stable dual-color control HepG2 cells in the same way. Efficacies of MYC and SUB1 dual knockdown and downstream targeted gene expression level were evaluated by qRT-PCR.

*Figure S 3-5. MYC-SUB1 dual knockdown*

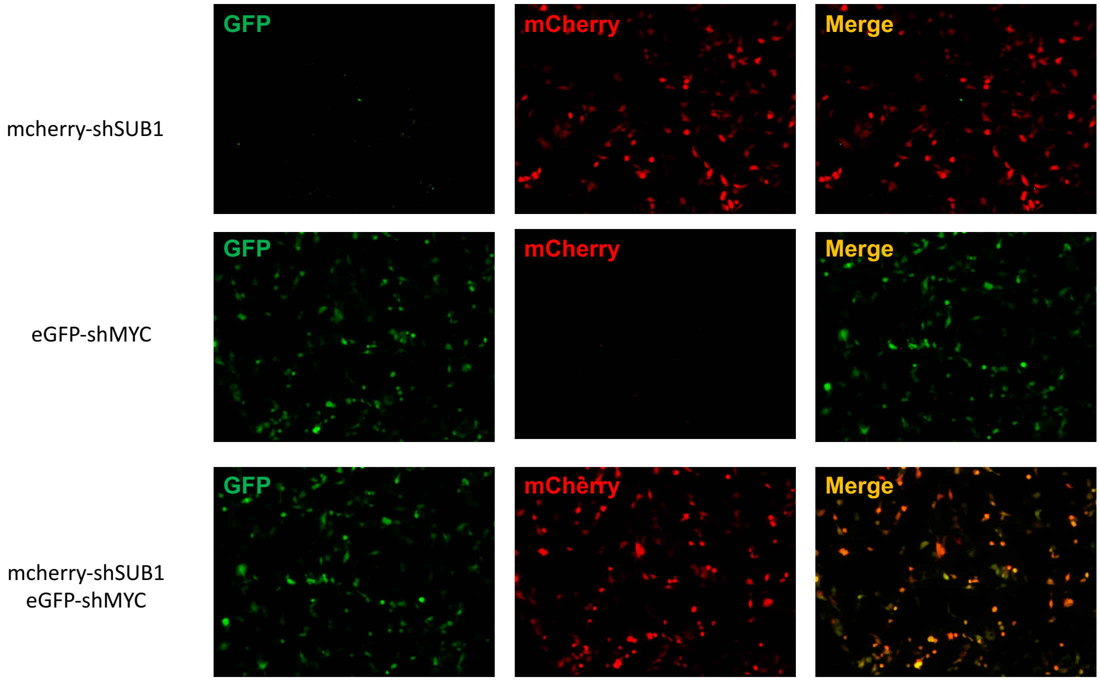

#### 3.6 (TL, ‖) Further investigation of SUB1

We found in 15 out of 16 cancer types, the target gene expression of SUB1 is upregulated (see Figure S 3-6). Hence, we further investigated it from multiple aspects.

*Figure S 3-6. SUB1 regulation in multiple cancer types*

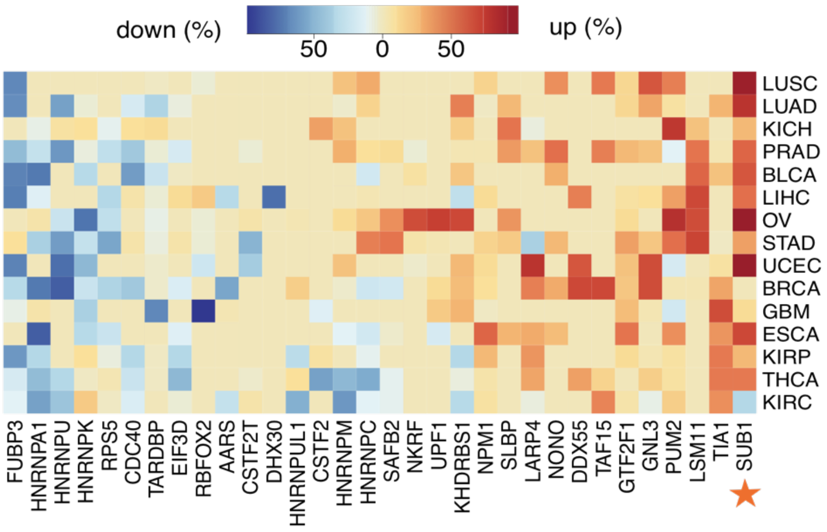

##### 3.6.1 (TL, ‖) Binding preference of SUB1 and its targets

We checked the 3' UTR expression level of SUB1 target genes and found that the target genes are significantly down-regulated upon SUB1 KD. In addition, we found enrichment of SUB1 target genes for CGC (Cancer Gene Census) genes. Among many of such genes, we have shown some examples together with SUB1 binding sites on the 3' UTRs.

*Figure S 3-7. Analysis of SUB1*

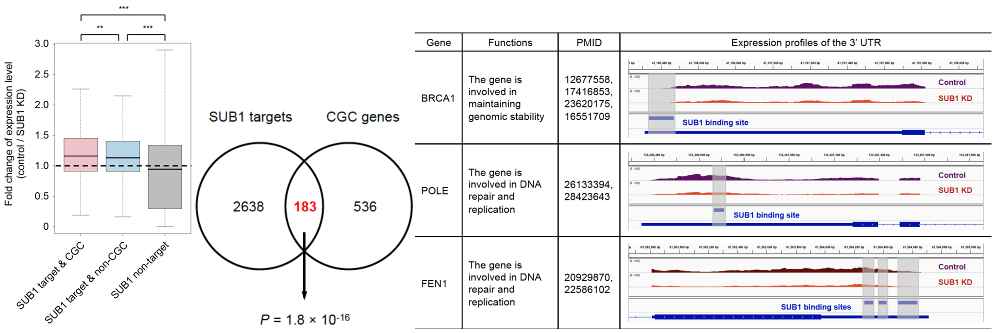

#### 3.6.2 (TL, ‖) Slower decay rate of SUB1 targets

The mRNA decay rates were determined by a previous study in HepG2 cell line[56]. We compared the accumulative distributions of mRNA decay rates between genes with 3'UTR SUB1 targets (weight = 1) and other genes without targets (weight = 0). The significance of decay rate difference was computed through the two-sided Wilcoxon rank-sum test. We observed a significant shorter mRNA decay rate for SUB1 targets compared to controls, implicating that SUB1 may stabilize mRNA transcripts (Figure S 3-8).

*Figure S 3-8. Slower Decay of SUB1 targets*

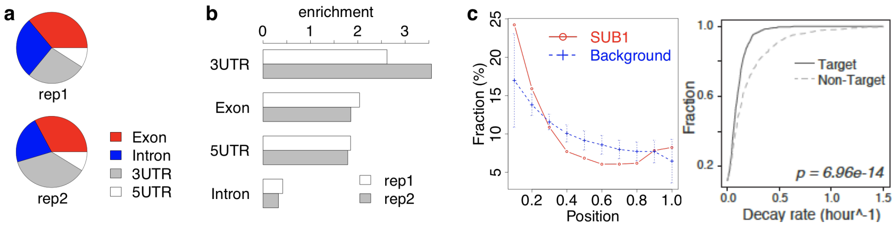

#### 3.6.3 (TL, ‖) Co-binding of SUB1 and MYC

Binding profiles of SUB1 and MYC (using eCLIP and ChIP-seq of HepG2) show these two regulatory proteins work together in certain genes. These genes include: BIRC5, CKS1B, MCM2, PLK1, ALYREF, LSM4, and MCM7. The MYC uniformly targets the TSS region of the gene, and the SUB1 predominantly targets in the 3' UTR region of the same gene, suggesting a potential co-regulatory nature.

Below is the signal track of several genes, such as MCM2 and BIRC5, which are well known oncogenes and demonstrate co-binding nature of MYC and SUB1.

*Figure S 3-9. Co-binding of MYC and SUB1*

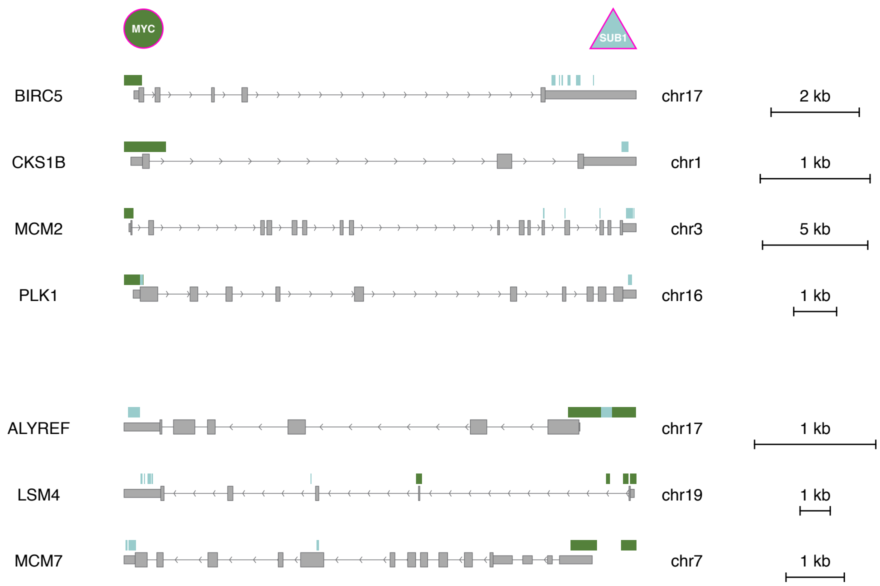

For each functional category from MSigDB, we applied a logistic regression to compute the enrichment of SUB1 targets after correcting the background effect of 3'UTR length and AU content.

a * 3'UTR degree + b* 3'UTR AU content + c* SUB1 eCLIP score + d = Function annotation

The 3'UTR length and AU content are background factors in the regression, which may have a significant association with specific functional categories. The reason for controlling these factors in regression is that the binding peaks of RBP, including SUB1, are typically enriched in long 3'UTR regions with a particular AU content depending on the RBP recognition motif. The response variable of logistic model is the binary functional annotation for each gene. For each function category, we computed the enrichment score of SUB1 targets as "c/StdErr(c)" through the max-likelihood estimation method in logistic regression. We applied this procedure for both Hallmark and KEGG sets from MSigDB.

Among genes whose 3'UTR regions have SUB1 eCLIP sites, we observed significant enrichment of functional categories including MYC targets, oxidative phosphorylation, and spliceosome. MYC activation induces an increase in total precursor messenger RNA synthesis, which increases the burden on the core spliceosome to process pre-mRNA. These results together indicate that SUB1 may stabilize the MYC target genes and pathways to promote the malignant growth of cancer cells.

*Figure S 3-10. Functional characterization of SUB1 targets*

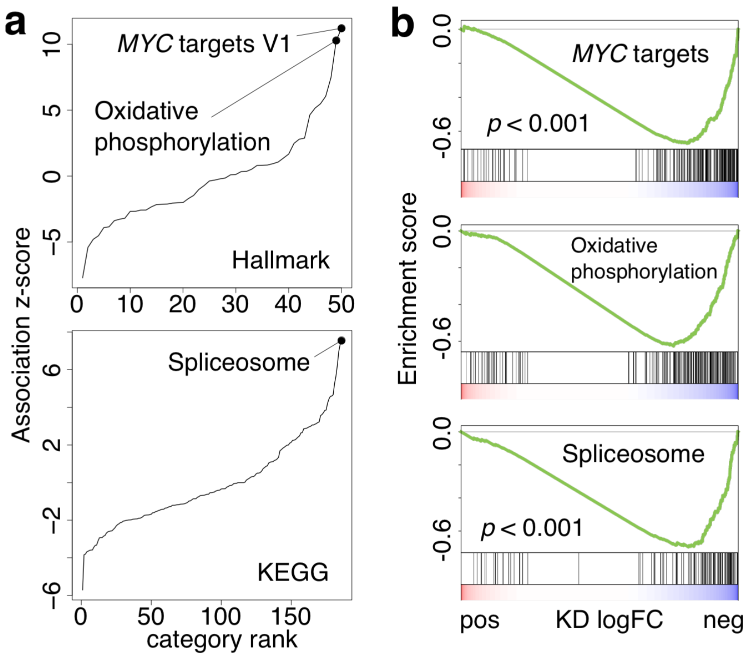

#### 3.7 (TL, Ⅱ) TF-RBP cross-talks

After building the join TF-RBP network for hierarchy map, we first evaluated the difference of TF/RBPs from various hierarchies to drive tumor-to-normal differential expressions. We found that the top level of TF and RBPs had larger correlation to drive target gene expression change. Details are given in Figure S 3-11.

*Figure S 3-11. Top level TFs have larger power to drive tumor-to-normal differential expression*

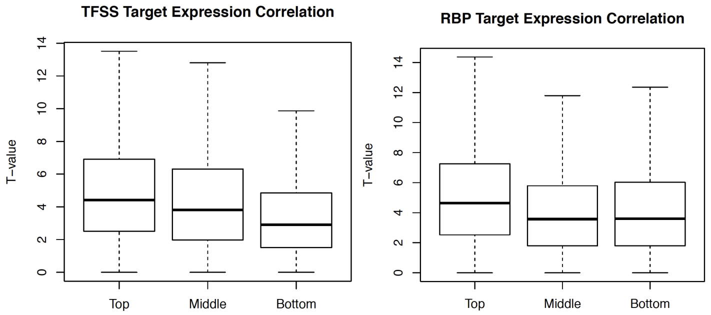

We evaluated the extent of crosstalk between TF and RBP regulators. We counted the number of regulatory edges between TF and RBP (crosstalk) in each level of hierarchy and tested for statistical significance. Comparing regulators in level 3 and 1 (top vs bottom), Wilcoxon rank sum test with continuity correction (one-sided) had p-value of 3.381e-16. Similarly, between level 2 and 1 (top vs bottom), the same test had p-value of 1.221e-09.

In addition, one dominant example of TF and RBP cross talk is given by MYC and SUB1. Specifically, MYC is seen to bind to the promoter region (across multiple cell types) and SUB1 is seen to bind to the 3' UTR region. This pattern of binding between MYC and SUB1 further strengthens the concept of TF and RBP cross talk in co-regulation. Examples are given below.

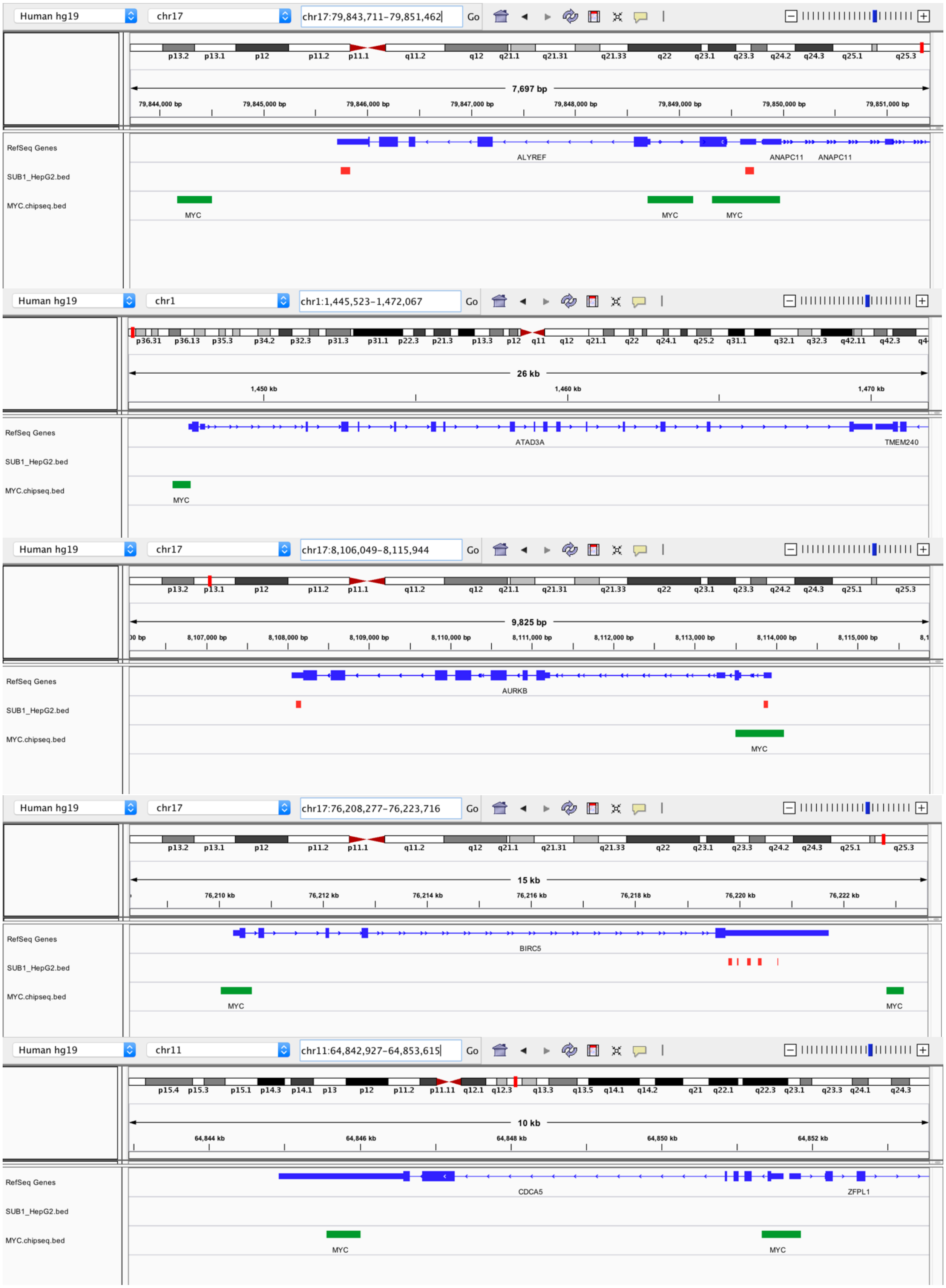
Human hg19
chr17
chr17:79,843,711-79,851,462
Go
7,697 bp
RefSeq Genes
SUB1_HepG2.bed
MYC.chipseq.bed
ALYREF
ANAPC11
ANAPC11
MYC
MYC
MYC
Human hg19
chr1
chr1:1,445,523-1,472,067
Go
26 kb
1,450 kb
1,460 kb
1,470 kb
RefSeq Genes
SUB1_HepG2.bed
MYC.chipseq.bed
ATAD3A
TMEM240
MYC
Human hg19
chr17
chr17:8,106,049-8,115,944
Go
9,825 bp
RefSeq Genes
SUB1_HepG2.bed
MYC.chipseq.bed
AURKB
MYC
Human hg19
chr17
chr17:76,208,277-76,223,716
Go
15 kb
76,210 kb
76,212 kb
76,214 kb
76,216 kb
76,218 kb
76,220 kb
76,222 kb
RefSeq Genes
SUB1_HepG2.bed
MYC.chipseq.bed
BIRC5
MYC
MYC
Human hg19
chr11
chr11:64,842,927-64,853,615
Go
10 kb
64,844 kb
64,846 kb
64,848 kb
64,850 kb
64,852 kb
RefSeq Genes
SUB1_HepG2.bed
MYC.chipseq.bed
CDCA5
ZFPL1
MYC
MYC

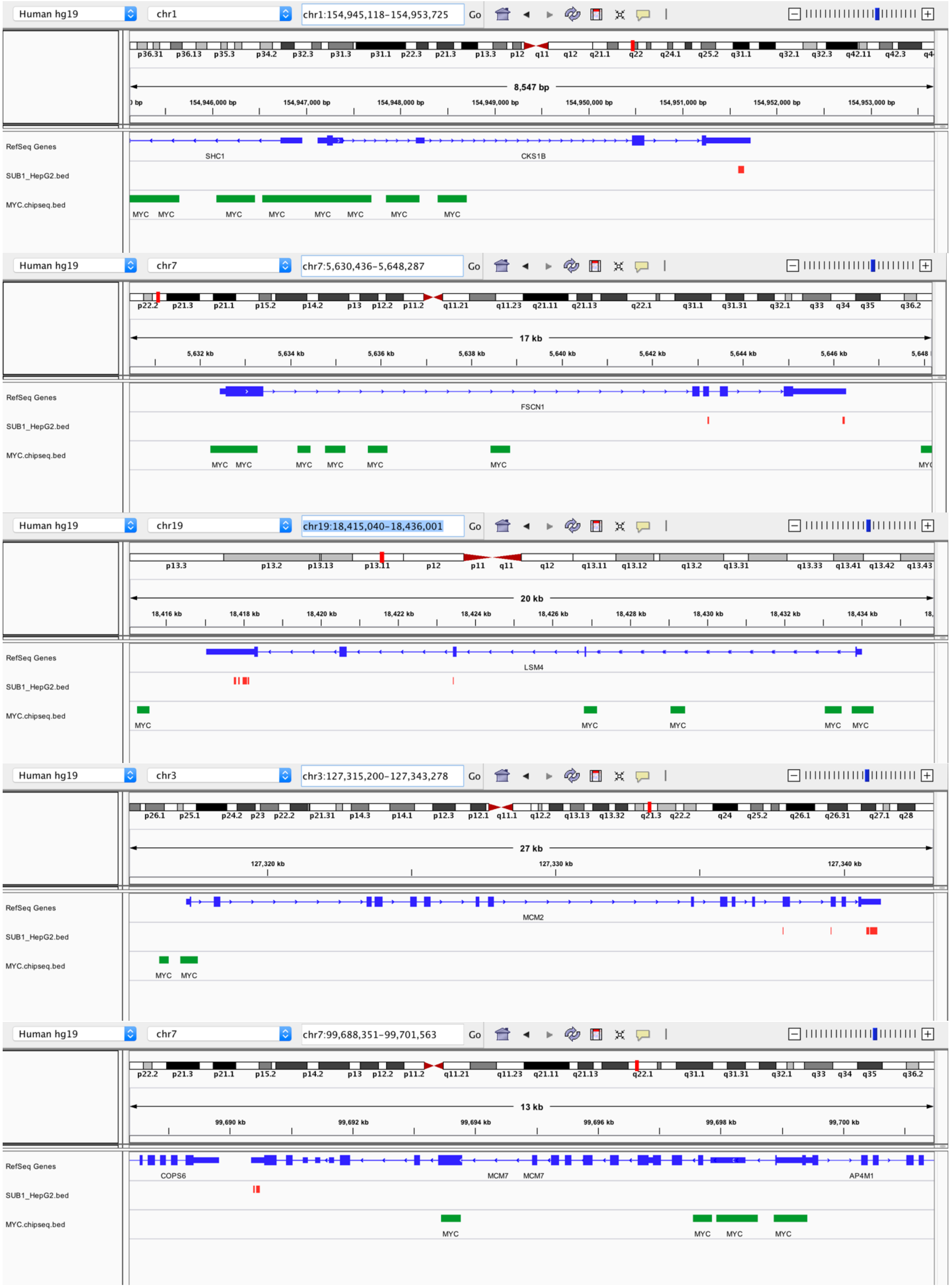
Human hg19
chr1
chr1:154,945,118-154,953,725
8,547 bp
RefSeq Genes
SUB1_HepG2.bed
MYC.chipseq.bed
SHC1
CKS1B
MYC
chr7
chr7:5,630,436-5,648,287
17 kb
FSCN1
chr19
chr19:18,415,040-18,436,001
20 kb
LSM4
chr3
chr3:127,315,200-127,343,278
27 kb
MCM2
chr7:99,688,351-99,701,563
13 kb
COPS6
MCM7
AP4M1

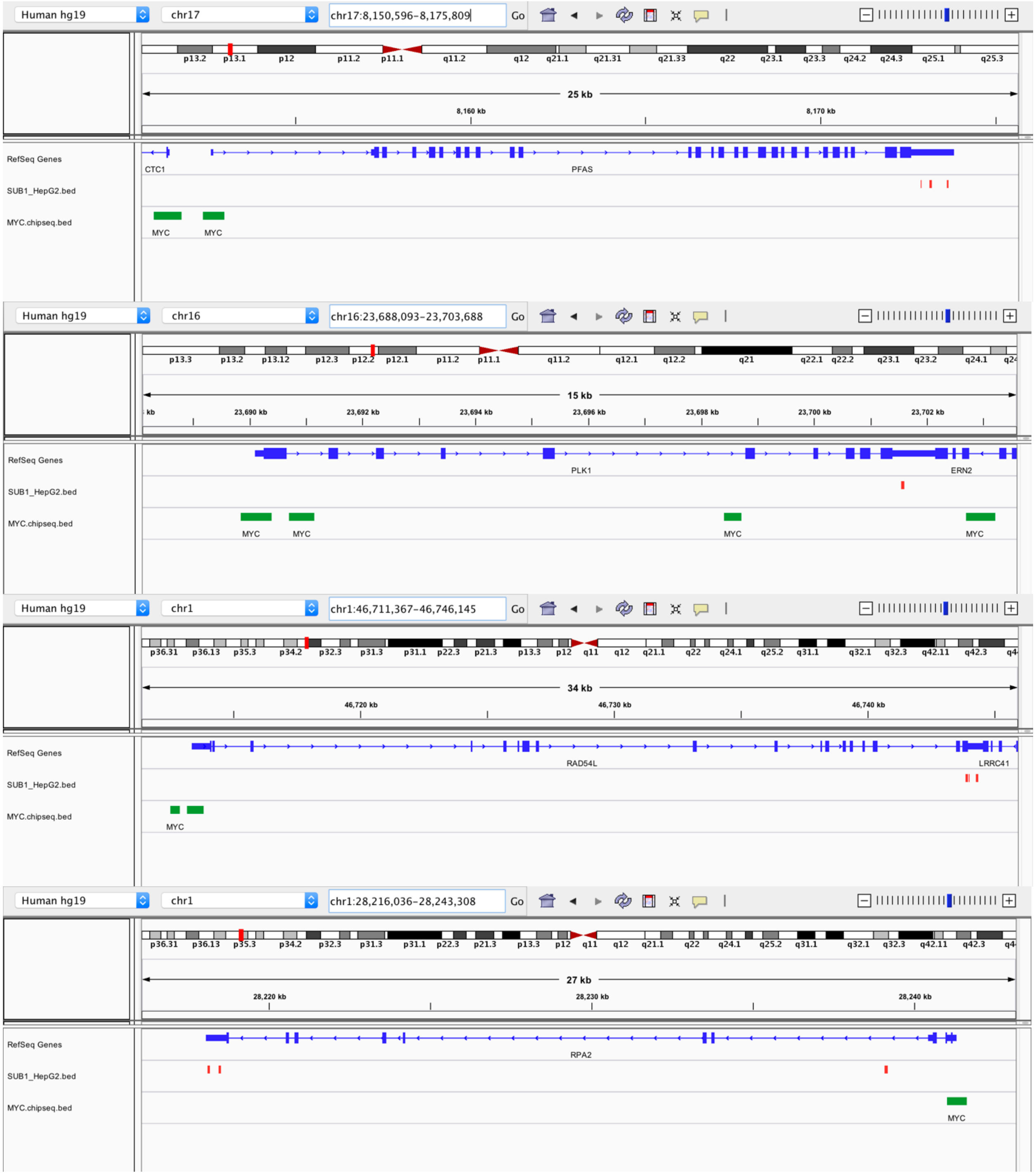

Human hg19
chr17
chr17:8,150,596-8,175,809
Go
25 kb
8,160 kb
8,170 kb
RefSeq Genes
SUB1_HepG2.bed
MYC.chipseq.bed
CTC1
PFAS
MYC
MYC
Human hg19
chr16
chr16:23,688,093-23,703,688
Go
15 kb
23,690 kb
23,692 kb
23,694 kb
23,696 kb
23,698 kb
23,700 kb
23,702 kb
RefSeq Genes
SUB1_HepG2.bed
MYC.chipseq.bed
PLK1
ERN2
MYC
MYC
MYC
MYC
Human hg19
chr1
chr1:46,711,367-46,746,145
Go
34 kb
46,720 kb
46,730 kb
46,740 kb
RefSeq Genes
SUB1_HepG2.bed
MYC.chipseq.bed
RAD54L
LRRC41
MYC
Human hg19
chr1
chr1:28,216,036-28,243,308
Go
27 kb
28,220 kb
28,230 kb
28,240 kb
RefSeq Genes
SUB1_HepG2.bed
MYC.chipseq.bed
RPA2
MYC

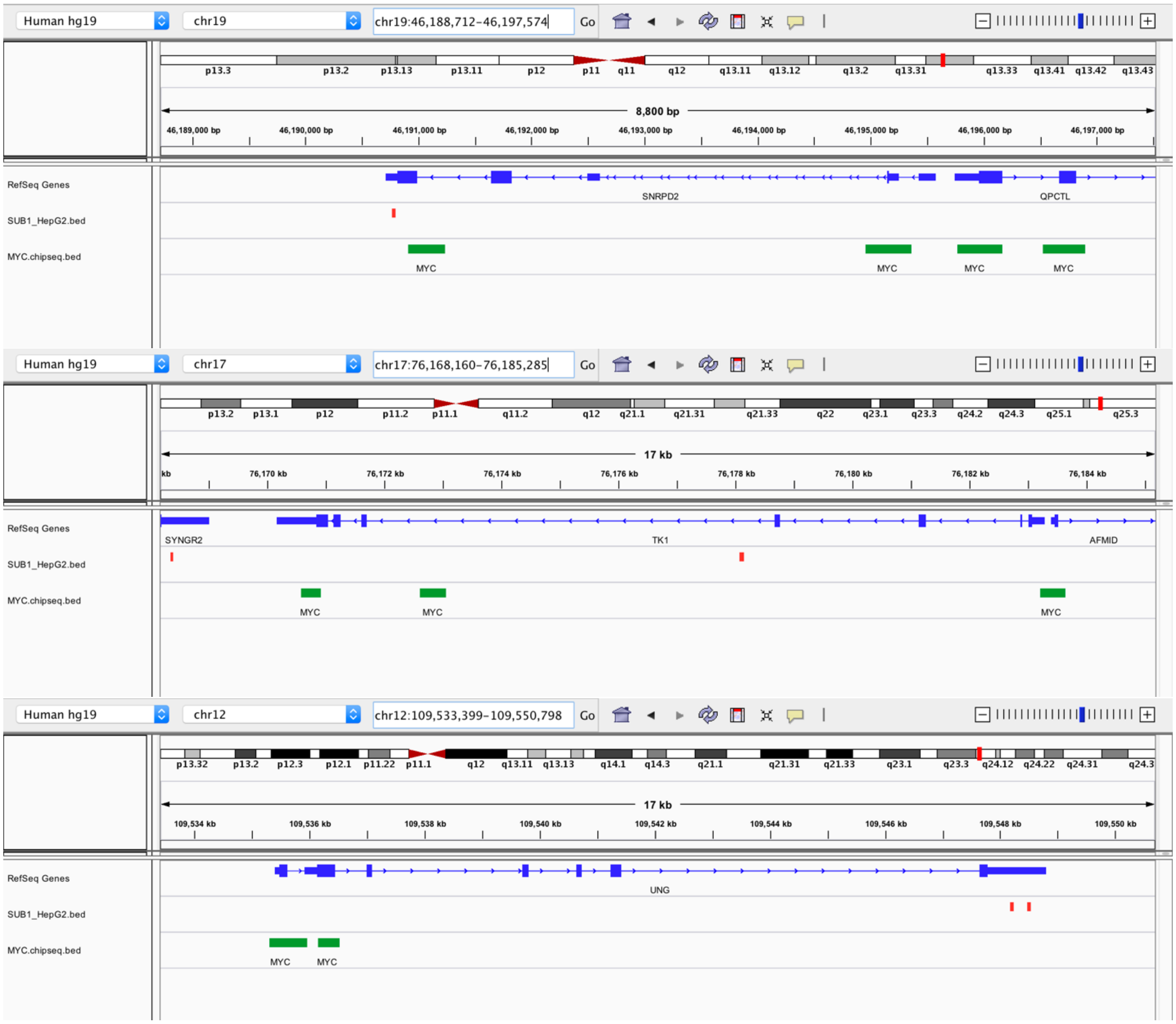

#### 3.8 (HL, ∦) Patient survival analysis based on TF activities

To evaluate the quality of ENCODE ChIP-seq derived network, we examined the association of TF regulatory activities with clinical data. We hypothesized that ENCODE regulatory network has clinical utility if it could be used to predict patient's survival.

##### 3.8.1 (TL, ∦) Systematic survival analysis of TFs on AML cohorts

In this analysis, we systematically calculated TF activity in 6 different AML datasets using the ENCODE ChIP-seq data. 292 ChIP-seq experiments from K562 (231 TFs) and 120 ChIP-seq experiments from GM12878 (101 TFs) were used to generate TF binding weight profiles from the TIP output. These profiles were generated by first implementing a z-test on TF binding scores to calculate p-values corresponding to each TF-gene binding interaction. These p-values were -log10-transformed and trimmed at -10 or 10 to construct the final weight profile. Weight profiles were min-max normalized so that all values were between 0 and 1. These weight profiles were used as input into the BASE algorithm[57]. The BASE algorithm takes in a single sorted (decreasing) patient's gene expression profile and calculates a running sum statistic by moving the down the profile and weighting each gene by its corresponding weight taken from the TF weight profile. This generates a foreground function. Similarly, a background function can be generated by repeating the process by multiplying by 1-weight instead of multiplying the gene by its weight. The maximum deviation between these two functions represents the TF activity score. This score is high when the highly-expressed genes in a patient's profile also tend to be bound tightly by the TF (as determined by the TF weight profile). It calculates TF activity scores for AML patient samples derived from the following gene expression datasets.

*Table S 3-4. Gene expression dataset of AML patients*

| Source | Accession | Author | Cohort |
|---|---|---|---|
| GEO | GSE37642 (GPL_96) | Herold | n=422 |
| NCI caArray | willm-0019 | Wilson | n=170 |
| GEO | GSE14468 | Wouters | n=526 |

Survival analysis was performed for each TF to identify those that were significantly associated with AML patient mortality. Namely, the TF's iRASs (activity scores) across patient samples were used as the independent variable in a Cox proportional hazards model. A hazard ratio <1 indicates that a TF's activity is associated with favorable prognosis and a hazard ratio of >1 indicates that a TF's activity is associated with unfavorable prognosis in AML patient samples. Since a separate model was fit to each TF's iRASs, p-values corresponding to the hazard ratios were adjusted for multiple hypothesis testing by using the Benjamini-Hochberg correction procedure.

In the results, we report the HR, P-value, and Adjusted P-value for each TF and their association with patient survival in each of the 3 AML gene expression datasets. The column labeled "number_datasets_significant_P005" indicates the number of datasets in which the TF's activity was observed to be significantly associated with AML patient prognosis at P<0.05. In particular, EZH2, STAT1, and NR2C2 TFs were found to be significantly associated with prognosis in all 3 datasets. 15 other TFs were found to be significant in 2 datasets. Notably, IKZF1, a well-known oncogene in hematopoetic and lumphoid cancers, was found to be significantly associated with prognosis in 2 datasets with p-value of 9.7e-4 and 1.9e-2 in the Herold and Wouters set, respectively.

##### 3.8.2 (TL, ∦) MYC-specific survival analysis on breast cancer cohorts

For this analysis, we focused on the METABRIC breast cancer cohort[58], which contains comprehensive survival information and gene expression profiles for 2,509 breast cancer patients. Using this dataset, we investigated the association between TF activity and disease-specific survival (DSS). Presumably, DSS provides stronger evidence compared to overall survival (OS) when evaluating whether some variable (TF activity) is associated with mortality from the actual disease of interest. In some cases, cancer patients die from some other co-morbid condition or the cause of death is unclear. Unlike OS where any deaths are considered "events" in a survival analysis, DSS considers only deaths caused by cancer. In most cases, evaluating overall survival provides greater statistical power but may not be as accurate as disease-specific survival. Since the METABRIC dataset provides detailed survival information which includes DSS, we used this metric over OS. Please note that all target genes were defined as those with p<0.01 from the TF TIP profile.

To determine if MYC activity was associated with breast cancer patients' survival, we utilized three separate MYC target gene sets to interrogate patient gene expression profiles derived from the METABRIC breast cancer gene expression compendia (n=1992).

First, based on ChIP-seq signal profiles, target genes of MYC were grouped into three categories; MCF-7 specific, MCF-10A specific, and common targets of the two cell lines. MYC activity as well as target genes from each of the three gene categories were calculated in each METABRIC breast cancer patient sample using BASE algorithm[57]. Second, we extracted time-to-event (death from breast cancer) information from the patients' metadata and correlated disease-specific patient survival with MYC activity score using univariate Cox regression models (proportional hazards).

An individual regression model was built for each MYC activity score corresponding to the three different gene sets. A positive association indicated that patients with high MYC activity exhibited longer survival compared to patients with low MYC activity and conversely, a negative association indicated that high MYC activity is associated with shorter patient survival time.

*Figure S 3-12. Survival analysis on MYC's from common target genes between MCF-7 and MCF-10A*

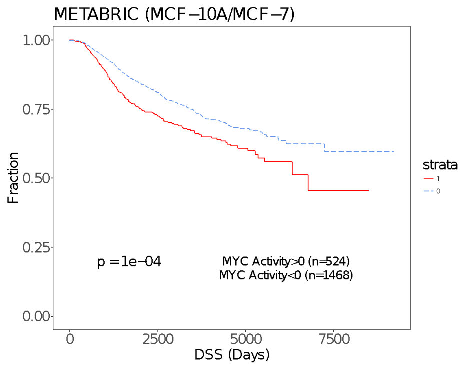

*Figure S 3-13. Survival analysis on MYC's MCF-7 specific target genes*

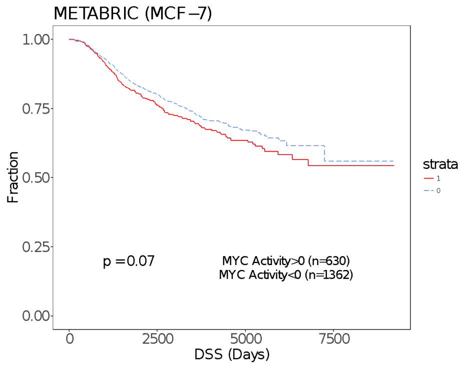

*Figure S 3-14. Survival analysis on MYC's MCF-10A specific target genes*

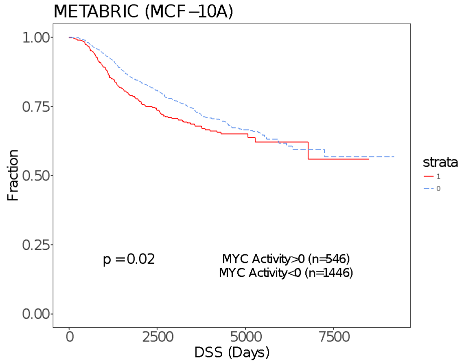

MYC activity survival analysis was also carried out focusing on the entire MYC binding profile in MCF-7 cell lines. Briefly, the p-values outputted by the TIP algorithm were transformed and used as weights as performed in the AML TF activity analysis. MYC activity levels were then used as the independent variable in a Cox regression model. This analysis was performed for three ENCODE replicates (Figure S 3-15).

*Figure S 3-15. Survival analysis on MYC's target gene expression levels in three replicates*

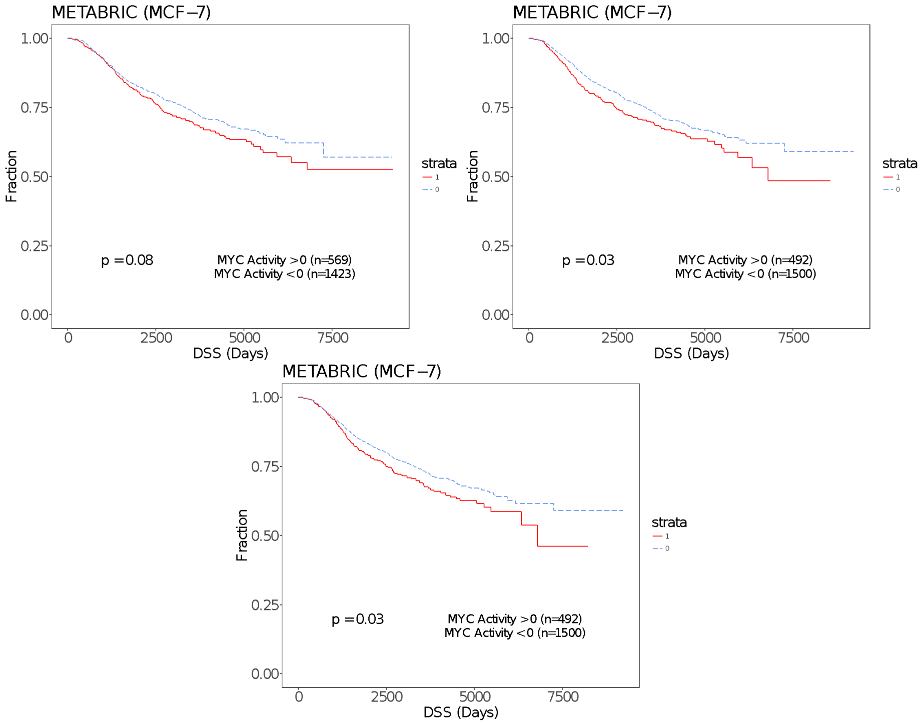

##### 3.8.3 (TL, ∦) Association of MYC and patient severity in chronic myeloid leukemia

For the CML analysis, target genes of MYC were categorized as K562-specific or GM12878-specific. MYC activity was calculated in each CML patient sample belonging to the GSE4170 dataset using target genes from each of the three gene categories. MYC activity scores were then compared between three different severity levels (chronic, accelerated phase, and blast crisis, Figure S 3-16) using ANOVA.

*Figure S 3-16. MYC activity scores were then compared between three different severity levels using ANOVA*

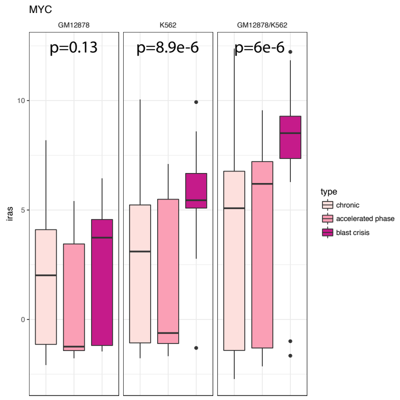

#### 3.9 (TL, ∦) Clinical relevance of the cell line model

We have made several efforts to verify the human clinical relevance of cell line data. For example, we clarified that although ENCODE data are profiled in cell culture models, the regulatory targets are still representative of the gene regulations in human cancers. Specifically, we predicted the regulatory activities of the transcription factor MYC using a ChIP-seq profile in MCF-7 cells. The MYC regulatory activity is highly correlated with the MYC expression across TCGA breast tumors (Figure S 3-17 a). For most TFs, their regulatory activities predicted using ENCODE ChIP-seq profile in cell lines are significantly correlated with their expression levels across breast tumors (Figure S 3-17 b). Moreover, using the same MCF-7 ChIP-seq profile, the MYC regulatory activity predicted for lung tumors is also significantly correlated with MYC expression level in TCGA lung cancer (Figure S 3-17 b). These results indicate that the ChIP-seq profiles from a particular cell line can capture the regulatory targets in human tumors from diverse cancer types. To select ChIP-seq or eCLIP profiles that are representative of the regulatory targets in human cancers, we only reported the results of TFs or RBPs whose regulatory activities are significantly correlated with their gene expression level in each TCGA cohort (Figure S 3-17 c).

*Figure S 3-17. Clinical relevance of cell line models; (a) The correlation between MYC expression level and regulatory activity across tumors; (b) The distribution of correlation p-values in TCGA breast cancer; (c) The fraction of regulators with statistically significant correlations in different cancer types for ChIP-seq and eCLIP networks*

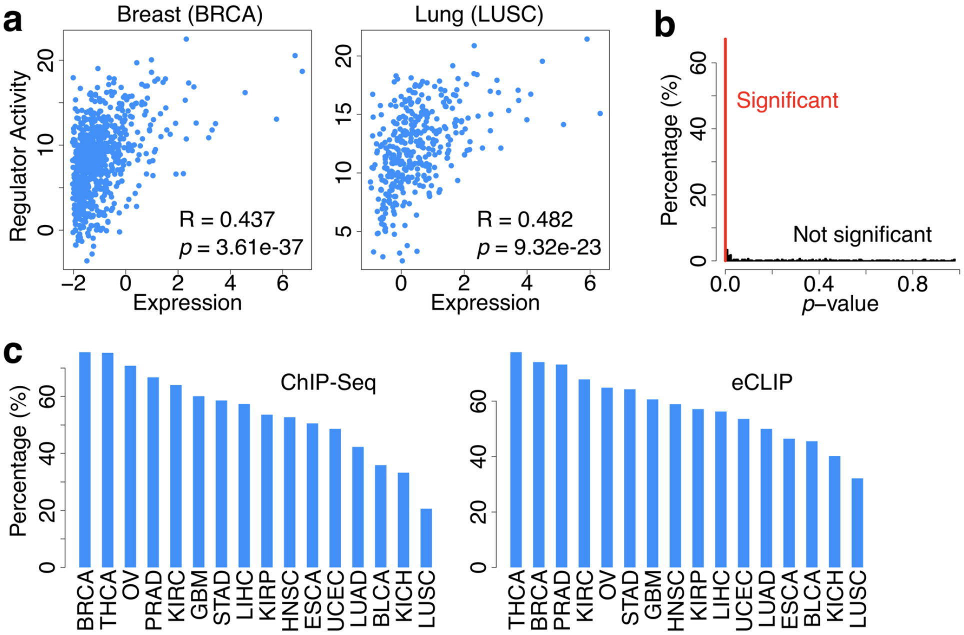

#### 3.10 (TL, ‖) Other MYC knockdown experiments

To establish the robustness of our network to the specific cell type and data provenance, we identified an alternative dataset. Specifically, we utilized a gene expression data set for MYC knockdown with a corresponding control in the Gene Expression Omnibus (GEO accession number GSE86504). For these alternative data, gene expression values were measured by RNA-seq in the HT1080 cell line. We note that, even though these alternative analyses were conducted on a different cell line, the results validate the behavior of the network, and they are consistent with our results using gene expression in the MCF-7 cell line.

*Figure S 3-18. MYC knockdown comparisons*

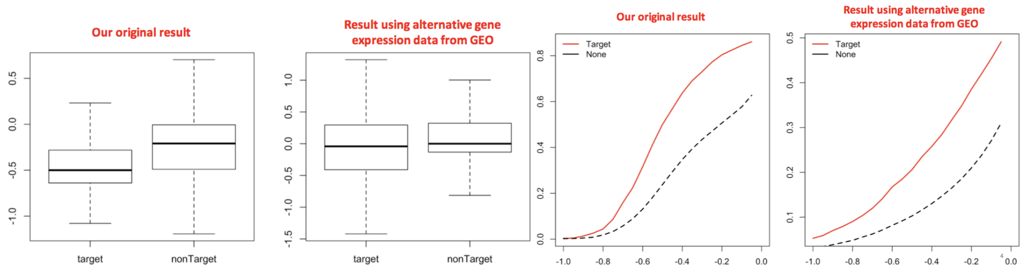

We also present microarray-based MYC knockdown data from previous work (GSE5823) in MCF-7, and we show that the results agree with our discoveries.

*Figure S 3-19. MYC knockdown in MCF-7 followed by microarray*

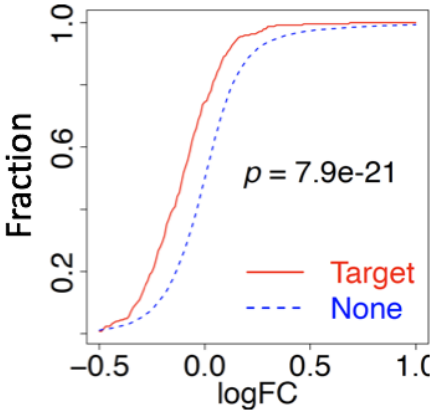

### 4 More details about “Measuring network rewiring”

#### 4.1 (TL, ∦) TF co-binding analysis

A score for each transcription factor pair $(i,j)$ is calculated based on the intersection of their binding sites. Let $TF_i$ has $n_i$ binding sites, each with a length of $l_{i,t}$ . Using the intersect function of bedtools, we found $n_{i,j}$ intersecting binding sites, each with length $l_k$ between $TF_i$ and $TF_j$ and calculated the co-binding score between $TF_i$ and and $TF_j$ as:

$$c(i,j) = \frac{\sum_{k=1}^{n_{i,j}} l_k}{\sum_{t=1}^{n_i} l_{i,t}} \tag{4-1}$$

*Figure S 4-1. Illustration of the co-binding analysis*

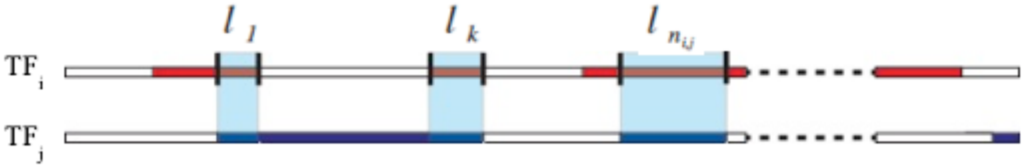

Please note that $c(i,j) \neq c(j,i)$, as the denominator is the total length of binding sites of TFi. We calculated the co-binding scores for each transcription factor pair in both GM12878 and K562 cell lines. We then calculated the differences between the cell lines by subtracting the co-binding scores.

We found that among all common transcription factors of GM12878 and K562 cell lines, ZNF274 has a larger binding score in K562 cell line compared to in GM12878 cell line. ZNF274 has the highest difference in the co-binding score with 54 out of 68 transcription factor partners, with an average difference of 0.61. The average difference for co-binding scores for all transcription factor pairs is 0.02. This shows that co-binding of ZNF274 increased significantly in K562 cell lines.

*Figure S 4-2. Co-binding scores c(i,j) of the transcription factors both in GM12878 and K562 cell lines. Rows represent TFi and columns represent TFj. Red color indicates high co-binding score, where blue color indicates low co-binding scores*

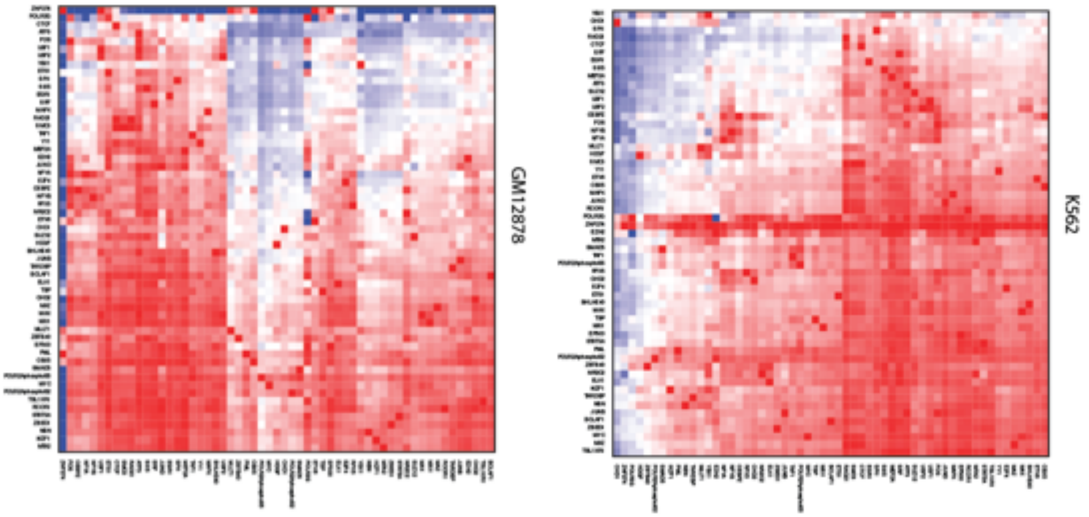

*Figure S 4-3. Difference of co-binding scores c(i,j) of the transcription factors between K562 and GM12878 cell lines. Rows represent TFi and columns represent TFj.*

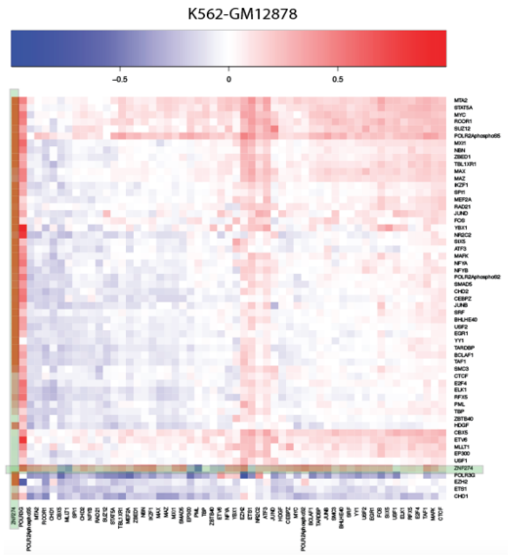

#### 4.2 (TL, ǁ) Epigenetic and expression change associated with rewiring

To evaluate the effect of TF-gene network rewiring, we investigated the associated epigenetic changes and target gene's expression with the gained and lost edges between normal and tumor samples. For expression, we used RESM quantification of ENCODE DCC uniformly processed long polyA RNA-seq and averaged TPM values over all available replicates. For DNase-seq, histone ChIP-seq, and methylation features, we further processed from fold enrichment signal tracks as follows. We averaged the fold enrichment signal across 200bp upstream and downstream of the unique TSS, the same canonical TSS site used define proximal TF-gene linkage. For all expression, DNase-seq, histone ChIP-seq, and methylation feature were expressed as a log2 ratio between tumor to normal samples. To avoid division by zero error, a pseudo-count of 0.0001 was added to each feature.

#### 4.3 Details about network rewiring analysis

##### 4.3.1 (TL, ||) Direct rewiring index calculation

We evaluated the rewiring of TF to gene linkages between normal and cancerous cells. To define TF rewiring between cell types, we first defined TF-gene regulatory network in each cell type using simple count-based target gene linkage. We used two different methods that examine TF to gene linkages based on their proximities to the TSS. For the TSS-based method, we simply used 2,500bp upstream and downstream of transcription start site (TSS) based on Gencode v19 annotation as a boundary for the proximal regulatory region. On average, 33.5% of TF ChIP-seq peaks fell into promoter region. We defined a target gene linkage if TF ChIP-seq peak was found within the boundary. However, we discovered, in Gencode annotation, there were numbers of genes that have more than 50 alternative TSS, which gave these genes unfair advantages of having more target gene linkages than others since their proximal regulatory regions can span up to 250kbp. Therefore, we selected one canonical TSS for each gene based on the total number of aggregated ENCODE TF ChIP-seq peaks. While this method is far from perfect, we believe this is the best method to capture the high-level TF network rewiring and quantify epigenetics changes around TSS while minimizing artifacts when counting all TSSs from all possible alternative transcripts.

In addition to TSS-based TF-gene linkages, we used target identification from profiles (TIP) method that quantitatively measures the regulatory relationships between TFs and target genes to define a subset of the full TF-gene network. For each TF, TIP model builds a characteristic, averaged profile of binding around the TSS and then uses this to weight the sites associated with a given gene, providing a continuous-valued 'regulatory' score relating each TF and potential target[46]. We used false discovery rate of 0.1 for cutoff. Since TIP uses narrower promoter definition than TSS-based method, we defined the TIP-based network as a subnetwork of the TSS-based network.

Both promoter-based linkages and enhancer target-based linkages were merged into one to build a complete TF-gene network. For more information about enhancer target-based linkages, please refer to Section 2.2. Two versions of full regulatory networks were constructed; one larger network by concatenating TSS-based network and enhancer-based network and another subnetwork by concatenating TIP-based network (details in 2.3.3) and enhancer-based network (details in 2.3.6). In addition, we built a merged network by combining all available ENCODE tissue types.

To quantify rewiring events, we first calculated rewiring score for each regulators (TFs). The fraction of the number of gain, loss, and common edges to the number of fully connected network edges, where all available TF nodes are fully connected with all available gene targets in the whole network was used to calculate the raw rewiring score.

Rewiring of edges between TF and target genes were compared in normal and tumor cells as shown in Figure S 4-4. If a target gene linkage was found in normal but lost in tumor, the edge was marked as loss edge. Similarly, if a target gene linkage was found only in tumor, it was labeled gain edge, and for edges found in both, they were labeled common or retained edges

*Figure S 4-4. Network rewiring schematics*

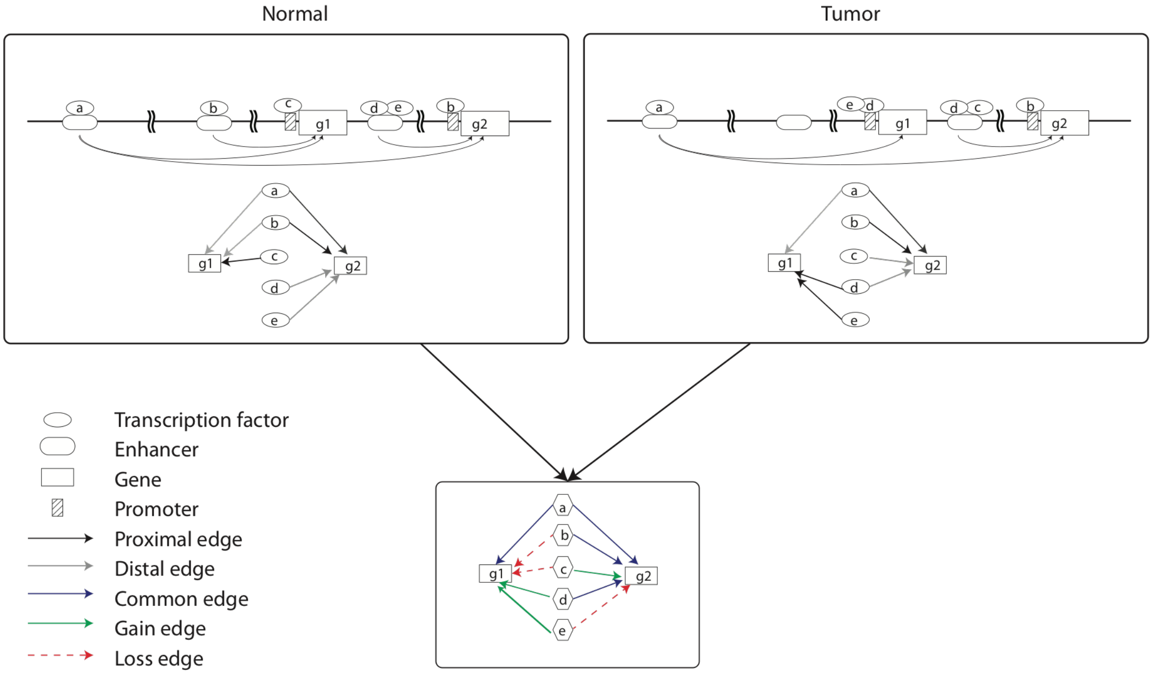

$$n_{fully-connected} = n_{TF} * n_{gene} - 1$$

$$rScore_{TF} = \frac{\frac{G_{in}+G_{out}}{L_{in}+L_{out}}}{\left|\frac{G_{in}+G_{out}}{L_{in}+L_{out}}\right|} \cdot \frac{(G_{in}+G_{out}+L_{in}+L_{out})}{n_{fully-connected}} \quad (4\text{-}2)$$

The rewiring score, rScore, after taking normalization over the maximum rScore, was used to rank the TF from the gainer to loser.

We evaluated whether the fraction of edge changes is related to the total number of edges for a given TF. For all 68 TFs common in K562 and GM12878, we calculated rScores using TIP network and plotted against the number of edges present in the original cell (GM12878 in this case). We fitted a linear model and found the adjusted R-squared of 0.6333 (. This result demonstrates that regulators with more regulatory edges (i.e., network hub) have higher tendency to rewire and thus have a higher rewiring index score.

*Figure S 4-5. Correlation of rScore with total number of edges*

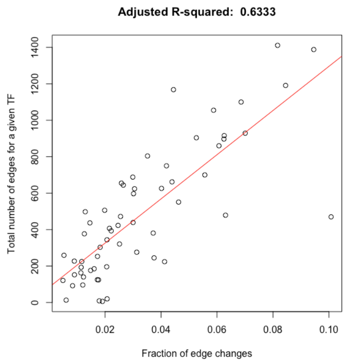

###### 4.3.1.1 (TL, ‖) Different null hypotheses in the network rewiring analysis

There are several scenarios that are possible when a given TF undergoes rewiring, such as

- No change: during tumorigenesis, the regulatory network may remain unchanged. In this case, all edges are retained. **This is our null hypothesis.**
- A total gain of edges: during tumorigenesis, a TF may become hyperactive and gain new regulatory edges. In this case, most edges are gained.
- Total loss of edges: during tumorigenesis, a TF may completely lose all its edges. This shut down of a TF can be caused by a deleterious mutation.
- Total change: all existing edges may be completely lost and replaced with a new set of regulatory edges. This is the expected behavior when edges are randomly permuted.

In our analysis, we assume a null hypothesis to be no change in regulatory edge across cell types. We expect no or minimal change in edges when two cellular contexts are similar. To demonstrate, we selected all available GM12878 ChIP-seq experiments that have at least two replicates, and we then calculated the same rewiring index between isogenic replicates of the same cellular context. We expect very small rewiring score given they are the same cellular context, and the edge changes between two networks will be simply a noise from ChIP-seq experiments.

As expected, when two cellular contexts are similar, as shown in “baseline”, a minimal number of edges do change targets. However, in “rewiring”, TF do change targets extensively when compared across cancerous (K562) to normal (GM12878) cell lines. To put this into perspective, we calculated the fraction of regulatory edges that are due to noise. We estimate that, on average, 1.36% of observed regulatory edges could be false positives.

*Figure S 4-6. Baseline levels of the rewiring analysis*

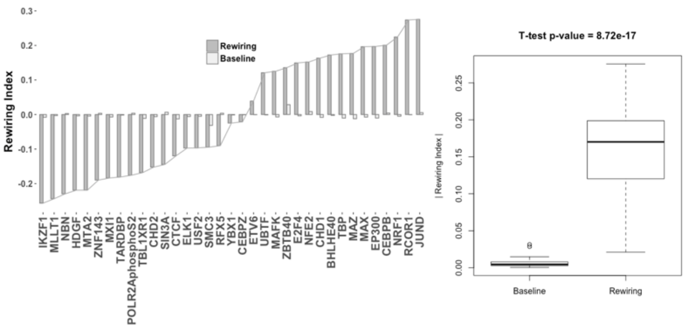

We evaluated the statistical significance of our model by measuring how much of the H1 network changes are due to noise and use of other normal cell types to evaluate how much of rewired edges overlap with H1. Bars represent the fraction of edges in H1. Above the blue dotted line, TFs were predicted to rewire toward H1. Below the red dotted line, TFs were predicted to rewire away from H1 (Figure S 4-7).

*Figure S 4-7. Rewiring analysis using different replicates*

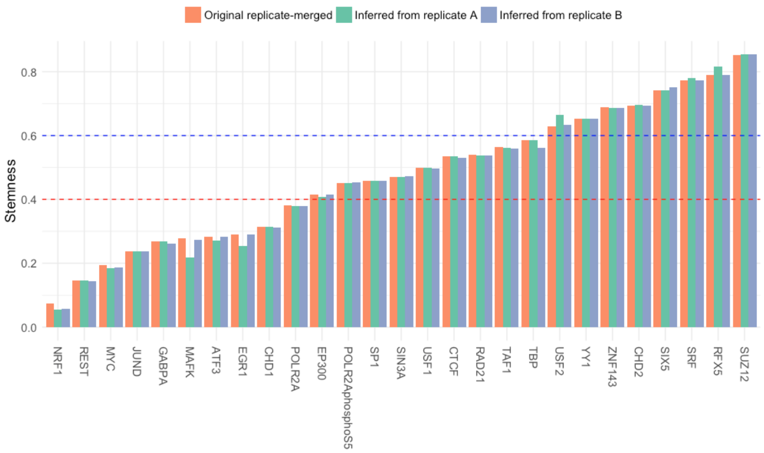

##### 4.3.2 (TL, ∦) Clustering of rewired TFs using rSCORE

Based on the fraction of gained, lost, and retained edges with respect to the total number of edges for each TF, rewired TFs were clustered into three groups using Kmeans clustering. Hartigan-Wong algorithm with 10 iterations were used a K-means algorithm by Hartigan shows the clustering result for rewired TFs between K562 and GM12878. NFE2 and RCOR1 were identified as one of the strongest members of the gained group, CTCF was identified as a member of the common group, and YBX1 was identified as a member of loss group (Figure S 4-8).

*Figure S 4-8. Kmeans clustering of rewired TFs in K562 and GM12878 using rSCORE*

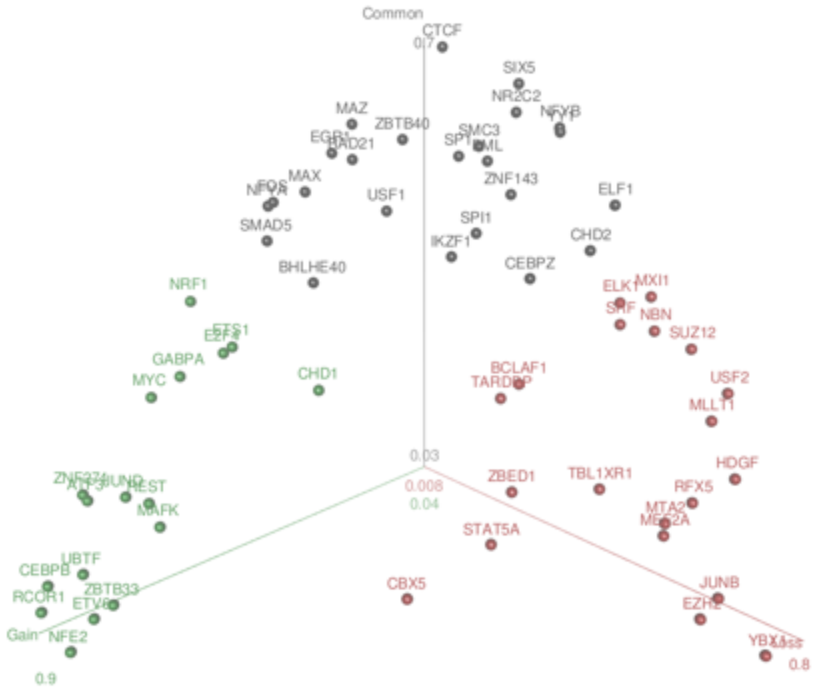

#### 4.3.3 Rewiring analysis based on mixed membership algorithm

##### 4.3.3.1 (TL, ‖) Details about the gene community models

We used mixed membership algorithm to investigate the rewiring changes between GM12878 and K562 cell lines. TF-target matrix ($M \times V$) is converted from enhancer and TSS regulatory network, where $M$ is the number of TF and $V$ is the number of unique target genes for all TFs. Each row represents a target gene of a TF $i = 1, 2 \dots M$. The regulatory pattern of each TF is comprised of $K$ latent communities. Each community includes contributions from $N$ target gene and $N$ varies for different TF.

The target gene of TF $w_i, \vec{w} = \langle w_{i,1}^v, w_{i,2}^v, \dots, w_{i,n}^v \rangle$ and $= 1$ means target, 0 means non-target. The observation denotes TF $i$, target gene $j$ with status $v$. Similarly, $Z_{i,j,v}$ is the community distribution of each target gene $j$ for TF $i$ with status $v$. $\beta(M \times V)$ denotes the probability of target gene $j$ belonging to community $k$, which is the parameter of a multinomial distribution. $\theta_i$ each and denote the distribution of communities for TF $i$. $\theta \sim Dirichlet(\alpha)$, where $\alpha$ is the super-parameter of $\theta$. When inferring the latent gene community model, we are most interested in the communities' parameter $\beta$, the Dirichlet parameter $\alpha$ and the latent community distribution $\theta$ of TF. So, the key is to find the posterior distribution of latent variables.

Figure S 4-9. Schematic of gene community-based rewiring analysis

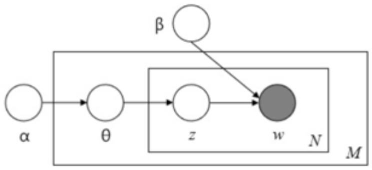

Variational EM algorithm (implemented using mixedMem R package) is used to infer the $\alpha$ and $\theta$ as described in Blei *et al*[59]. However, computational benefits of EM lead to optimization uncertain and make it easily converge to local maxima. We have no prior knowledge for the $\theta$ and $\alpha$, which is impossible to use near plausible value to find a reasonable optimum. To hack this, we repeat multiple times (100) and use the median of rewiring changes from all the non-early stops simulation to represent the most optimal regulatory changes of TF. One example of the $\theta$ distribution was given in the following figure**.** The rewiring of TF regulation is defined by the changes of distribution in K gene communities using $Distance_i = \sqrt[3]{\sum_{i,j}[\sqrt[3]{q_{K562,i,j}} - \sqrt[3]{q_{GM12878,i,j}}]}$, where $q_i$ is the distribution of communities for TF $i$.

Figure S 4-10. Example distribution difference in tumor and normal cell lines

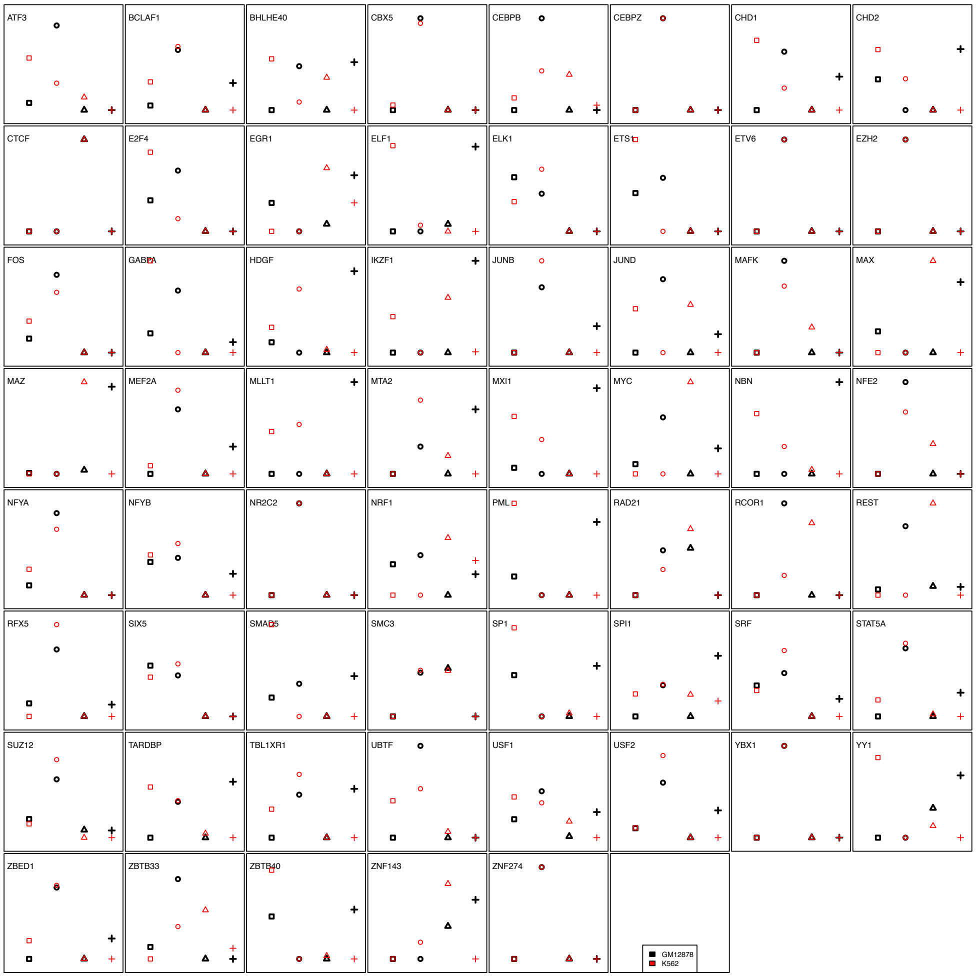

###### 4.3.3.2 (TL, ∦) Comparison of the gene community model with other models

Mixed membership model is a hierarchical Bayesian topic model framework and can help to uncover the underlying semantic structure of a document collection. The core of topic models is Latent Dirichlet Allocation (LDA), which cast the mixed-membership (topics) problem into a hidden variable model of documents. The LDA model has been widely used to analyze a wide variety of data types, including but not limited to text and document data, genotype data, survey and voting data. The advantage of LDA over other algorithms (like SVD, PLSI) used in semantic analysis has been described in Blei *et al*[60]. In particular, LDA allow document to belong to multiple topics simultaneously, and the topic mixture weight was treated as k-hidden random variable to reduce overfitting problem rather than a set of individual parameters that explicitly link to the training set.

Due to the specificity of data type (TF target network) and problem-definition, there is no other benchmark method to compare. If we treat the LDA mixed-membership analysis as a dimensionality reduction problem, it is possible to compare how well of a model can reproduce the information of original data, as described in the paper by Guo & Gifford[61]. The correlations of the original target gene vectors between two TFs are compared with those of dimension reduced vectors. The better method should be much close to original vectors correlations.

*Figure S 4-11. Comparison of mixed membership model with other methods*

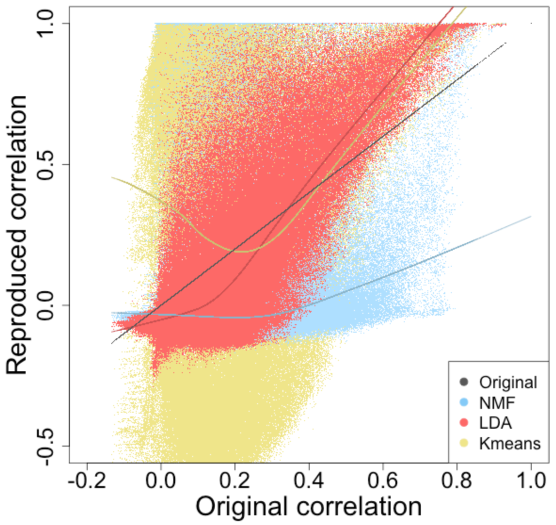

To explore how well the LDA mixed-membership analysis on TF regulatory network, we extend our dataset from 122 GM and K526 samples to all the 862 TF ChIP-seq assays included in ENCODE data portal. In order to get a reliable correlation, we also increase the number of topic to 50 as the number of TF sample increases. The non-negative matrix factorization (NMF) and Kmeans clustering are used for comparison because the nature of regulatory network requires a non-negative decomposition. The same target dimension K =50 was used to NMF and target number of clusters K=50 for Kmeans. The Euclidean distance between each data the centroids are used to calculated the correlation. As shown in the figure, the x-axis is original correlation of two TF regulatory target, y-axis is reproduced correlation from LDA document to topic distribution and NMF decomposed matrix. The solid line is the ‘loess’ smoothing curve for the scattered dots. We can see the LDA method can reproduce the original correlation better than either NMF or Kmeans. Overall correlation between the reproduced pairwise correlation and the original correlation were 0.123 in Kmeans, 0.404 in NMF and 0.788 in LDA (Figure S 4-11).

#### 4.4 (TL, ǂ) Rewiring of TF across tumor types

Rewiring of several key TFs in leukemia (K562) were evaluated across lung (A549), liver (HepG2), and breast (MCF-7) cancer models. In some extreme cases, the overall direction of rewiring was reversed. For BHLHE40 in CML as an example, the direction of rewiring is mostly towards gaining patterns, whereas in lung adenocarcinoma, the edges were dominantly lost. For both JUND and MYC, the pattern of gain edges was consistently observed in liver or breast cancer samples.

*Figure S 4-12. Example of TF rewiring in other cell types*

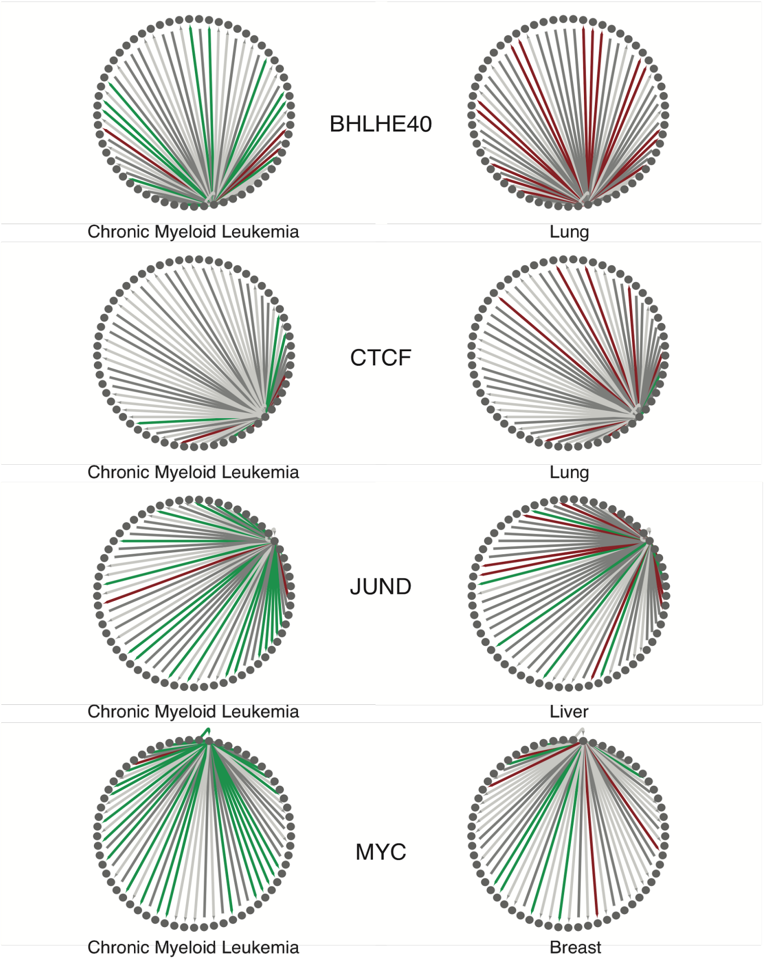

#### 4.5 Genomic profiles around the rewired edges

##### 4.5.1 (TL, ‖) Chromatin mark aggregation

We defined 200 bp upstream and downstream of TSS to be the core regulatory regions of the gene. We calculated the average DNase-seq and various histone ChIP-seq signals across 400 bp window for all protein coding genes. We then compared the average chromatin level at the core regulatory regions between normal and cancerous biosamples (i.e., K562 to GM12878) by taking log fold change (with base 2). To measure the chromatin effects on rewiring, we aggregated activating and repressive chromatin marks around gained and lost JUND regulatory edges and found that activating marks are enriched with gained edges while repressive marks are depleted.

##### 4.5.2 (TL, ‖) Rewired edges affected with variants

We examined gained and lost edges that harbor SNVs and SVs. We first identified TSS regions affected with gained and lost events. We intersected SNV and SVs with 2500 bp up and downstream of TSS region. Specifically, we first merged all deletions with 50% reciprocal overlapping from two call sets: Zhou et al[62] and Dixon et al[63].

*Figure S 4-13. JUND in CML*

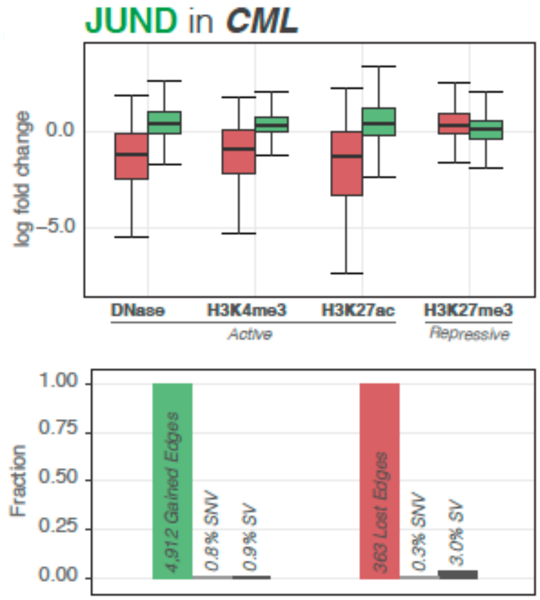

#### 4.6 (TL, ‖) Tissue-specific network hierarchies

We used a subset of K562 proximal network containing 46 common sequence-specific TF (TFSS) in K562 and GM12878. Hierarchy of the TF within the network was determined using HierNet algorithm[55] using 3 layers with default modeling parameters. Each TF node was colored based on the rewiring index, and the size was determined based on the percent of TFBS affected with cancer somatic mutations. We colored the edges based on rewiring status from rewiring index analysis.

#### 4.7 (TL, ‖) Building imputed networks and application to PCAWG dataset

We first built a cell-type specific imputed network by concatenating and merging the DHS peaks from all available ENCODE DNase-seq experiments of similar tissues and cell types. We assigned the average DHS score to each element and then searched for TF-binding events on the combined DHS sites using motif PWM. We applied the imputed networks to PCAWG dataset by matching cancer types to networks using their cell-of origin or tissue-of-origin. PCAWG variants ware mapped to evaluate potential motif disruption events. Using the distance of motif to TSS, strength of DHS peak, and motif disruption score (d-score) to evaluate edge loss per gene. We retained the largest d-score for each motif position. We further filtered out the motif disruptive event that has d-score lower than 10 or DHS score lower than 20.

Given each TF-gene pair a disruptive score D:

$$D = 1 - \Pi_i(1 - d_i * \exp(-A * l_i)) \;\; (i = 1,2,\dots,n)$$

Where $n$ is the number of the disruptive events in each TF-gene pair, $d = \frac{1}{1+\exp\,(x*\alpha_1+\alpha_0)}$, where $\alpha_1 = \log\left(\frac{1}{0.9} - 1\right)/10,\;\; \alpha_0 = \alpha_1 * (-10),\;\; A = \log(0.5)\,/(-1000)$, $l$ is the distance of motif to TSS.

*Figure S 4-14 D-score function and decay function*

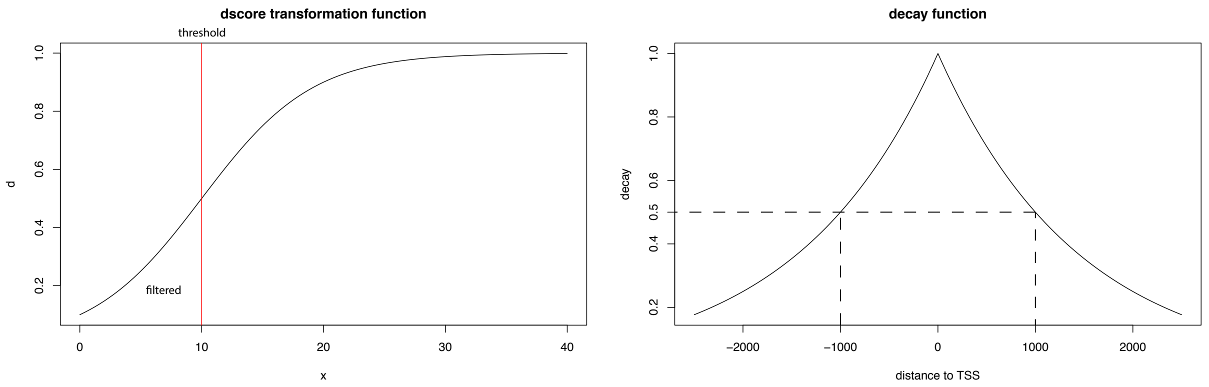

For each cancer type of PCAWG, we summarized all the TF-gene pairs that have disruptive events with a disruptive score. We further filtered the edge based on PCAWG gene expression profile for each cancer type, where we only retain the disruptive events where the TF has average expression greater than FPKM of 1.

Finally, we took the average disruptive score of each gene within each cancer type. And we chose several gene list (PCAWG driver, Cancer Gene Census, Transcription Factors, RNA-Binding Proteins, Olfactory Genes, Essential Genes) and compared their average disruptive scores normalized by the average disruptive score of all genes.

### 5 More details about "Placing cancer cells in the context of many cell types"

#### 5.1 Cell type clustering using PCA/RCA

##### 5.1.1 (TL, ‖) Gene Expression

We collected all available poly A long RNA-seq data from the ENCODE portal. There was 329 RNA-seq experiments as of March 2018. Based on uniformly processed gene expression quantification on Gencode v19 annotation, we merged replicates by taking the average FPKM across replicates. We filtered for 20,345 protein-coding genes based on Gencode v19 annotation. The ENCODE RNA-seq matrix containing 20,345 genes and 329 biosamples across 108 cell types was projected onto 2-dimensional space using Reference Component Analysis (RCA). RCA was an algorithm developed to improve clustering in single-cell and was comparable to PCA[64]. We removed the 1st RCA axis (similar to PC1 in PCA) to remove potential batch effect and projected using RC2 and RC3 (see Figure S 5-1). However, as shown in the shadow figure below, the overall stemness trend was unchanged whether the first axis was used or not.

*Figure S 5-1. Clustering of ENCODE RNA-seq gene expression using PC1 and PC2*

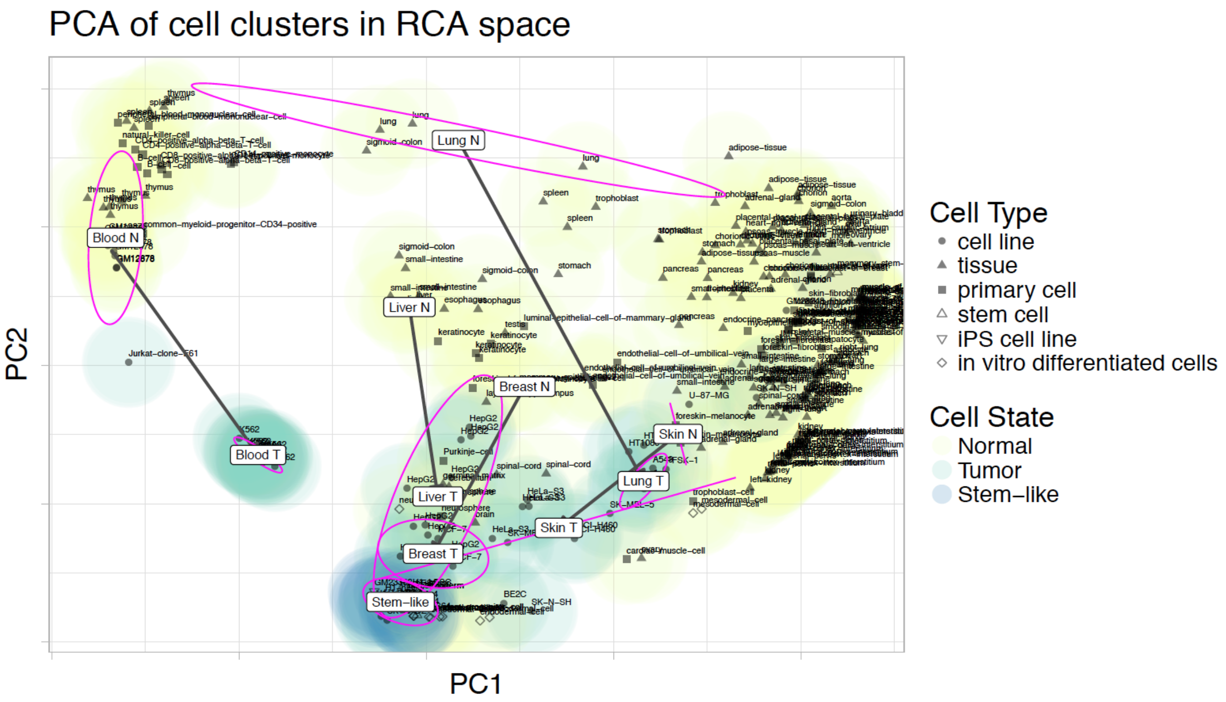

Figure S 5-2. Shadow figure of Figure 4A, Clustering of ENCODE RNA-seq gene expression

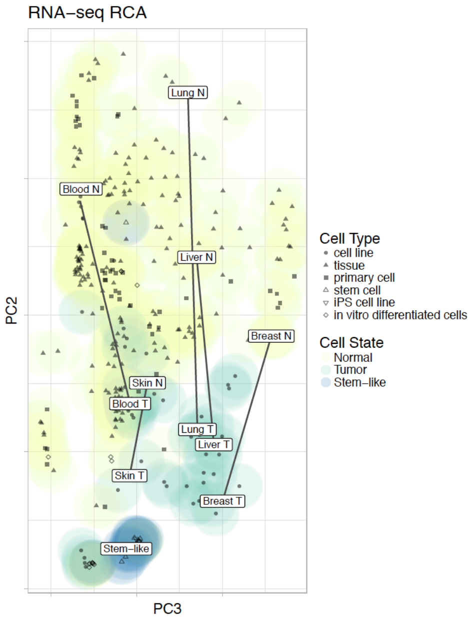

#### 5.1.2 (TL, ‖) Proximal Network

For the proximal network, we created a matrix of 14,536 TSS (2.5kb up/downstream) with CTCF peaks across 207 cell types. We have chosen CTCF ChIP-seq since it has (1) broad cell type coverage in ENCODE and (2) plays a role in defining chromatin structure. We then looked for CTCF binding sites that are found in at least 20 different cell types called hotspots. We found 9,506 CTCF hotspots near TSS across 207 cell types. We projected this proximal network using PCA.

Figure S 5-3. Clustering of ENCODE CTCF ChIP-seq proximal network and ccRE enhancer distal network

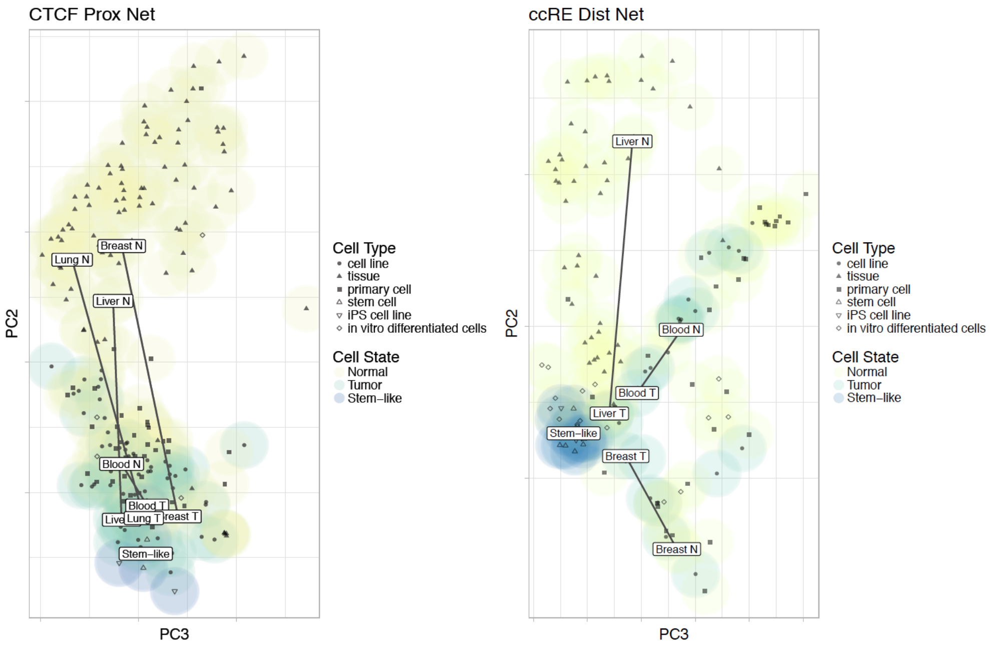

##### 5.1.3 (TL, ||) Distal Network

For the distal network, we collected 990,079 merged ccRE ELS (Enhancer-Like Signature) sites across 609 ccRE annotations. Since the network was too sparse to be effectively projected onto PCA space, we further subset the network by two criteria. First, we looked for an element that are at least 100kb away from TSS. Second, we looked for ccRE ELS element that are found in at least 20 different cell types. We found 13,497 ccRE ELS hotspots across 134 cell types and projected them onto PCA space (see Figure S 5-3).

In summary, we found consistent stemness pattern using network information as compared to results using RNA-seq. Cancerous biosamples had closer distance (similarity) to stem-like biosamples than normal biosamples, measured in terms of gene expression, proximal network, and distal network.

*Figure S 5-4. Relative distance to stem-like clusters*

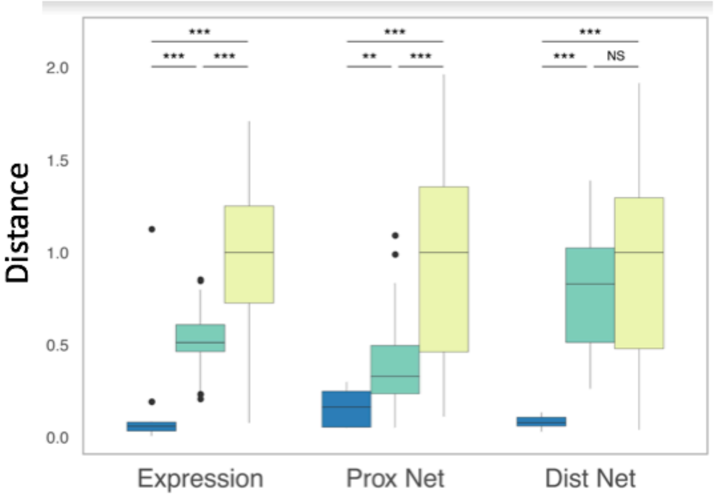

#### 5.2 Incorporating TF knockdowns

##### 5.2.1 (TL, ||) Projecting TF knockdowns on the RCA figure

We collected 661 knockdown data (531 shRNA-KD, 4 siRNA-KD, 50 CRISPR-KO, 76 CRISPRi) targeting 589 TF/RBP from ENCODE portal. We processed the data similarly to ENCODE RNA-seq data. We merged replicates by averaging FPKMs across replicates, annotated in Gencode v19 protein coding genes. For each experiment, we paired with respective control experiment and projected the data onto RCA space defined in RNA-seq analysis.

*Figure S 5-5. Projection of ENCODE knockdown data onto the RCA space. Blue represents control and purple represents a cell after knocking down respective protein target*

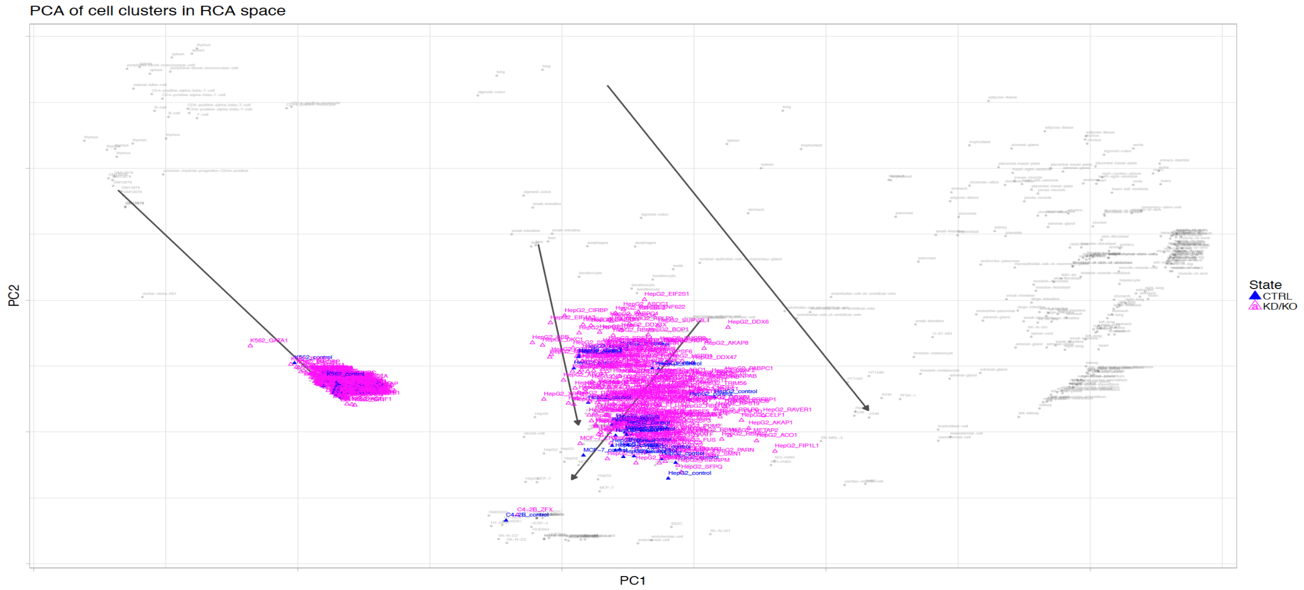

*Figure S 5-6. projection of ENCODE knockdown data onto the RCA space. Start of an arrow represents the placement of the cell before knockdown (control experiment) and the tip of arrow represents the placement of the cell after knocking down respective protein target*

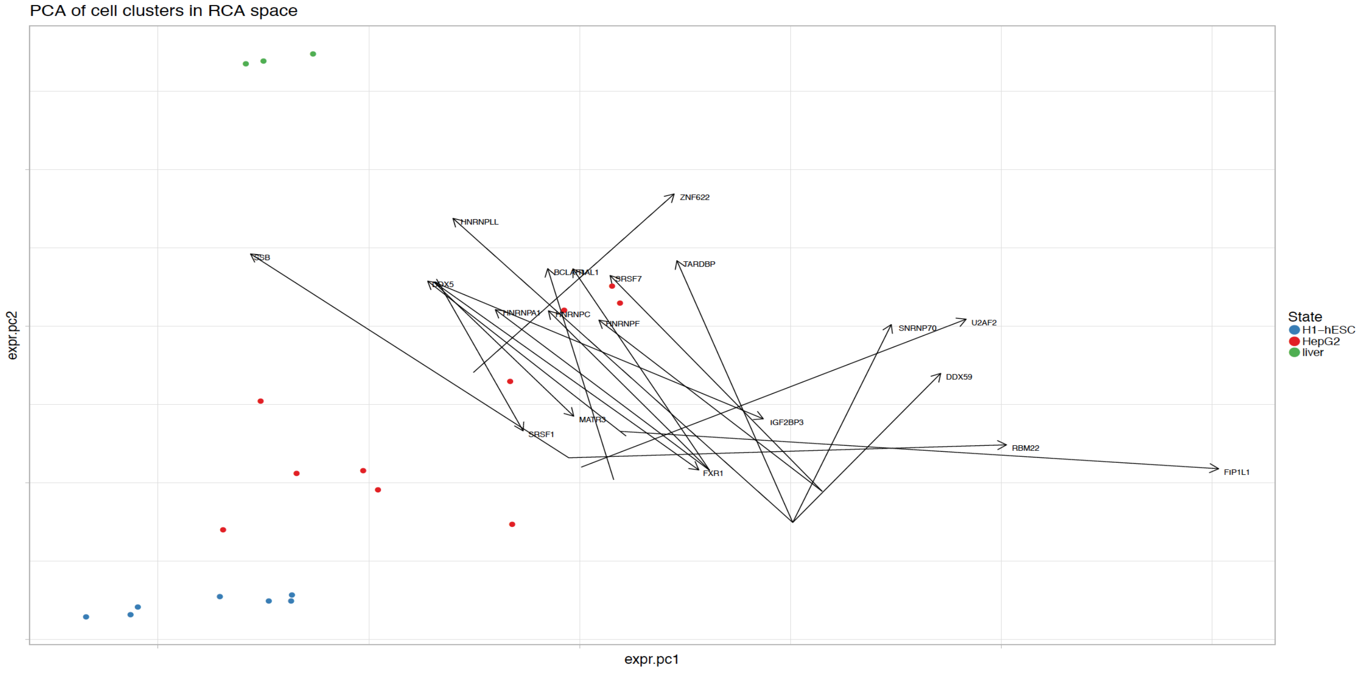

#### 5.2.2 (TL, ‖) TF knockdown consistency analysis

Based on the Cancer Gene Consensus (https://cancer.sanger.ac.uk/census), we categorized TF knockdown experiments into different classes; 8 knockdown (KD) genes were tumor suppressors (TSGs), 10 KD genes were oncogenes, and 210 KD genes were neither TSGs or oncogenes. The cancer transcriptome was defined as the expression of the 529 cancer related genes in the cancer gene consensus. For each of the 18 gene KDs in K562, we first calculated the changes of the 529 cancer genes from those in the K562 control, i.e. increases or decreases in gene expression. If a cancer gene's change in the K562 KD was in the same direction as in GM12878, compared to the K562 control, this cancer gene upon knockdown expresses similarly as in GM12878. The fraction of such cancer genes indicated the similarity of cancer transcriptomes between the K562 upon a knockdown and GM12878.

#### 5.3 Using ENCODE features to jointly estimate BMR in cancer

##### 5.3.1 Collection and processing of the features

###### 5.3.1.1 (HL, ǂ) Histone modifications data

The full catalog of histone modification data is collected from the ENCODE project website and binned into 1Mb windows in a ready-to-use format to predict BMR. The summary of cell types is give in Figure S 5-7, with the majority of histone data coming from primary cell and tissue sources:

*Figure S 5-7. Summary of Histone Marks*

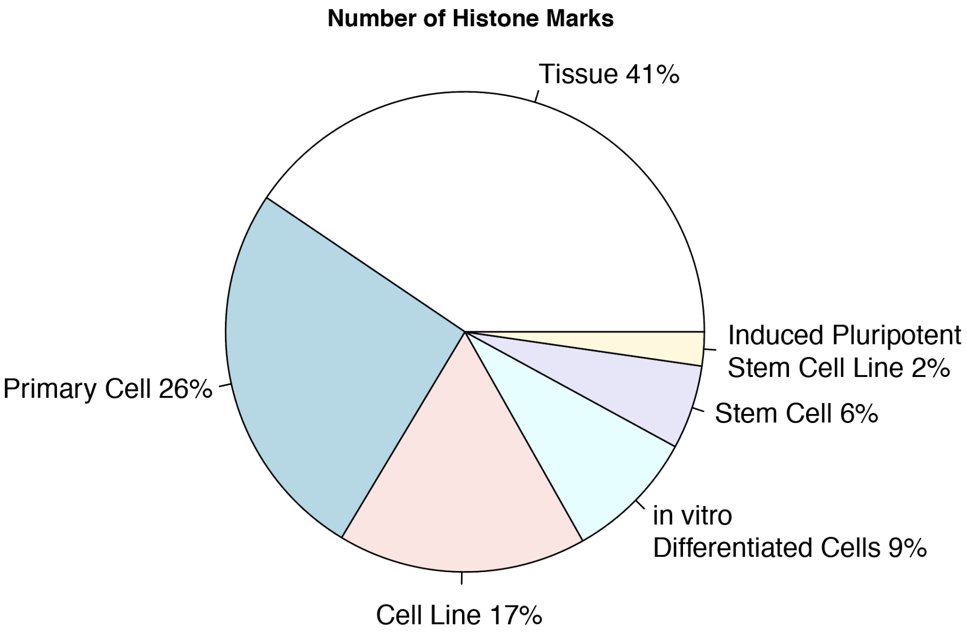

###### 5.3.1.2 (HL, ǂ) Replication timing data

In total we aggregated 51 replication timing features across different cell types, summarized in Figure S 5-8. Detailed data processing was described in 1.5.1. The resulting signal tracks was used to calculate average signals per 1Mb windows in a ready-to-use format to predict BMR.

*Figure S 5-8. Summary of Replication Timing Data*

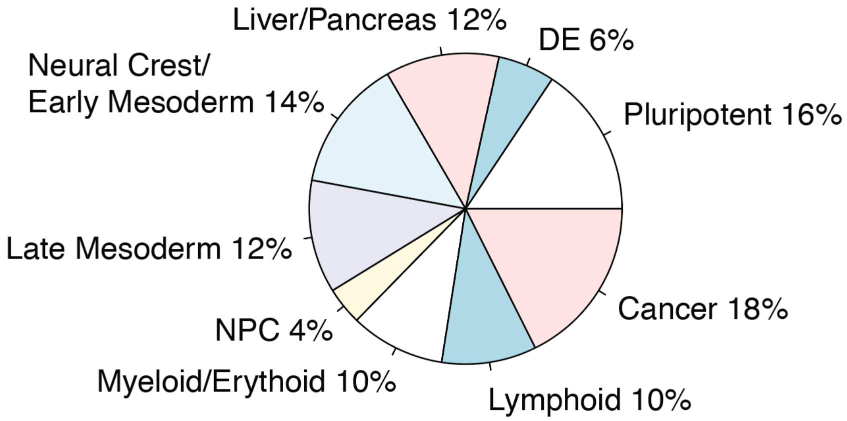

#### 5.3.2 (HL, ‖) Benefit of increased number of features

There are three main reasons why the increased number of features is beneficial for BMR estimation.

Firstly, accurate cell-of-origin definitions are challenging. Distinct subtypes of a tumor may derive from different 'cells of origin'[65]. Recent literature has pointed out the cell-of-origin effect on tumors from multiple aspects, such as mutational process and tumor classifications. However, to accurately define the cell of origin of tumors is sometimes challenging. For example, even different subtypes of a tumor from the same organ may originate from different cell types. In this case, the richness of ENCODE data provides us with a larger pool to find the best representative cell of origin.

Secondly, epigenome from a single cell type is usually incomplete and due to the correlated natures of features from different cell types, it is usually beneficial to features from sub-optimally matched cell types rather than ignore the missing data from the optimally mapped cell type. For example, the Roadmap Epigenome data fall into different classes, with class 1 being a complete epigenome for samples. However, Class 1 data is very limited, and many of the samples do not include comprehensive epigenetic features.

*Figure S 5-9. Lack of completed Epigenomes from the Roadmap Data*

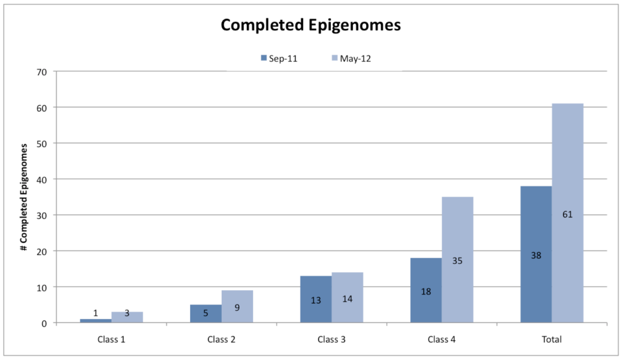

We also downloaded all available uniformly processed ENCODE DNase-seq, histone ChIP-seq, and RNA-seq data from the ENCODE portal on 3/2/2018. We counted once if a biosample has more than one experiment from different labs and filled the matrix shown below. Several biosamples (i.e., H1-hESC) have deeper coverage (vertical lines) where most biosamples have limited histone ChIP-seq assayed.

*Figure S 5-10. DNase-seq, histone ChIP-seq, and RNA-seq data availability matrix for all ENCODE biosamples*

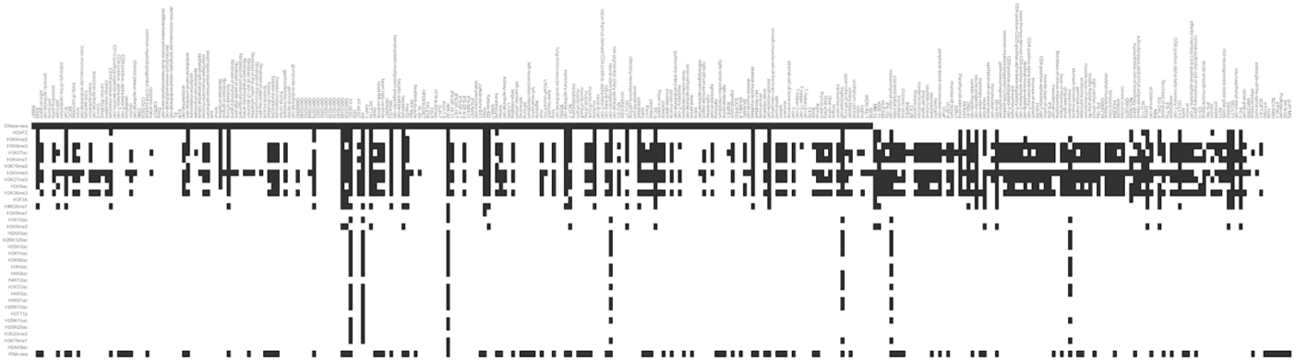

Thirdly, due to the heterogeneity of cancer, the best set of features does not necessarily come from one cell type. For example, we extracted related histones modification features from both tissue (Breast epithelium) and cell line (MCF-7) from ENCODE and correlate it with mutation counts per 1mb bin. We found that the best correlated feature for each histone may come from either cell type (Table S 5-1).

#### 5.3.3 Comparison of data from cell lines and tissue

##### 5.3.3.1 (TL, ‖) Single feature comparison

We calculated the Pearson correlation of the breast cancer mutations count per Mbp versus various histone modification features in tissue and cell line. Cell line data provides comparable (and sometimes better) correlation with mutation counts.

*Table S 5-1. Pearson correlation of BRCA mutations and various histone marks*

| Histone Mark | Tissue | Cell | Cell+Tissue |
|---|---|---|---|
| H3K4me1 | -0.479 | -0.433 | 0.499 |
| H3K4me3 | -0.481 | -0.434 | 0.484 |
| H3K9me3 | 0.424 | 0.437 | 0.515 |
| H3K27me3 | 0.372 | 0.411 | 0.491 |
| H3K36me3 | -0.535 | -0.396 | 0.544 |

We have listed the detailed features we used in the following tables.

*Table S 5-2. ENCODE MCF7 Covariates*

| Cell Type | Histone Mark |
|---|---|

| | |
|---|---|
| MCF-7 | H2AFZ |
| MCF-7 | H3F3A |
| MCF-7 | H3K27ac |
| MCF-7 | H3K27ac |
| MCF-7 | H3K27me3 |
| MCF-7 | H3K27me3 |
| MCF-7 | H3K4me1 |
| MCF-7 | H3K4me2 |
| MCF-7 | H3K4me3 |
| MCF-7 | H3K4me3 |
| MCF-7 | H3K9ac |
| MCF-7 | H3K9me2 |
| MCF-7 | H3K9me3 |
| MCF-7 | H3K9me3 |
| MCF-7 | H4K20me1 |

*Table S 5-3. ENCODE Breast Tissue Covariates*

| Cell Type | Histone Mark |
|---|---|
| Breast epithelium | H3K27ac |
| Breast epithelium | H3K27me3 |
| Breast epithelium | H3K36me3 |
| Breast epithelium | H3K4me1 |
| Breast epithelium | H3K4me3 |
| Breast epithelium | H3K9me3 |
| Breast fibroblast | H3K27ac |
| Breast fibroblast | H3K27ac |
| Breast fibroblast | H3K27me3 |
| Breast fibroblast | H3K27me3 |
| Breast fibroblast | H3K36me3 |
| Breast fibroblast | H3K36me3 |
| Breast fibroblast | H3K4me1 |
| Breast fibroblast | H3K4me1 |
| Breast fibroblast | H3K4me3 |

*Table S 5-4. ROADMAP Breast Covariates*

| Cell Type | Histone Mark |
|---|---|
| Breast myoepithelial | H3K4me1 |
| Breast myoepithelial | H3K4me3 |
| Breast myoepithelial | H3K9me3 |
| Breast myoepithelial | H3K27me3 |

Breast myoepithelial H3k36me3

###### 5.3.3.2 (TL, ‖) Joint BMR modeling comparison

We estimated the BMR using negative binomial regression similar to previous methods[54,66]. Specifically, we calculated the total number of mutation counts per 1mb bin by summering up all patient mutations, and then regressed it against four set of features: 1) features only from MCF-7; 2) features only from breast tissue in ENCODE (Epithelial Fibroblast); 3) features from both MCF-7 and breast tissue 4) features from the Roadmap breast tissue (Breast Myoepithelial).

We preformed forward selection to select the best features to use for BMR prediction. With the same number of features used, the combination of tissue and cell line data is noticeably better than data from any single source.

*Figure S 5-11. Comparison of joint modeling of BRCA mutations using various features*

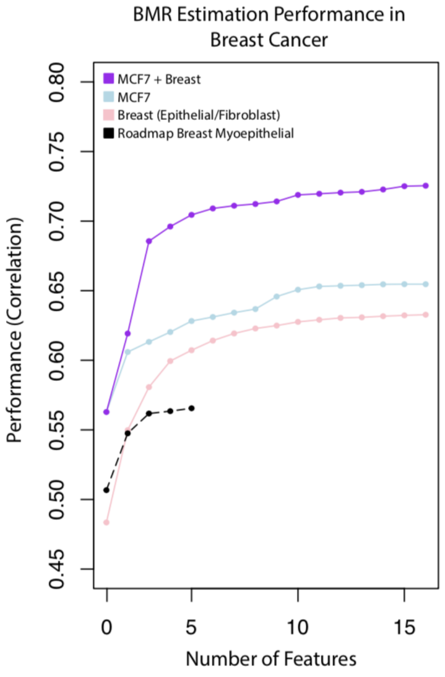

##### 5.3.4 (TL, ‖) Model selection and cross-validation for BMR estimation

We put together all the somatic variants of all patients with the same cancer type and count the total number of mutations within 1mb bins. We used all negative binomial regression to predict the mutation counts using features from ENCODE. To demonstrate the contribution of the cumulative effects from many features, we used negative binomial regression to predict the BMRs similar to Martincorena *et al*[54] and Nik-Zainal *et al*[66]. We used forward selection on the 2066 features to select the best combination of features for performance comparison.

### 6 more details about “The value of the extended gene for variant interpretation”

#### 6.1 Differential gene expression analysis

##### 6.1.1 (TL, ‖) Data used for the analysis

We downloaded the expression data from cancer cohort studies as well as the mutation profiles of each patient. We have merged somatic variant data from TCGA and previous publications and selected 95, 198, and 327 samples from TCGA for CLL, breast cancer, and liver cancer in our analysis..

##### 6.1.2 (TL, ‖) P-value calculation based the extended gene annotation

For each cancer type, we loop through each gene performing a differential expression by mutation status analysis. For each gene, we consider the extended gene annotation associated with the gene. Samples demonstrating a mutation in the extended gene definition are separated from those samples with no mutation in the extended gene annotation. A Wilcoxon test is performed between the expression values of mutated samples versus those of non-mutated samples. As a comparison, we also perform this analysis on different annotation sets, where the control group is the set of samples with no mutation, and the test group is the set of samples showing at least 1 mutation in the annotation. We perform the Wilcoxon test on the two distributions of expression for the following annotations: extended gene, distal regulatory element, eCLIP, TFBS, TSS, and CDS. The log p-value of each gene and cancer type is given. Figure S 6-1 shows the result of SRSF2 in liver cancer patients

*Figure S 6-1. Example of differential expression stratification by gene-level mutational status*

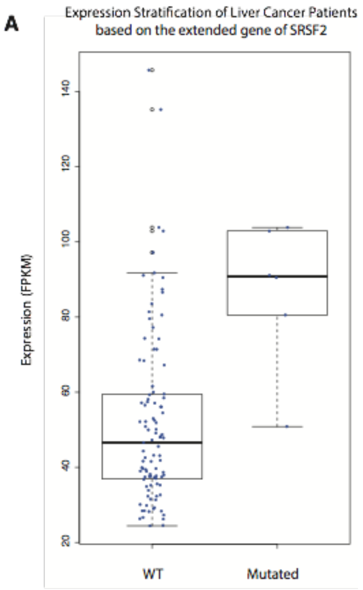

##### 6.1.3 (TL, ‖) Summary of results

In Table S 6-1, each cell represents the number of genes for which the extended gene annotation outperforms the other annotation, divided by the number of genes for which the other annotation outperforms the extended gene. One annotation is considered to outperform another if the associated p-value is more significant. A number greater than 1 suggests that the extended gene is better at finding significant differential expression based on mutation status.

*Table S 6-1. Performance summary of extended gene vs. other annotations in determining differential expression by mutation status*

| | **Extended Gene vs. Other Annotations** | | | | |
|---|---|---|---|---|---|
| Cell Type | CDS | UTR | TF | eCLIP | Enhancer |
| HepG2 | 2.12 | 2.22 | 1.83 | 3.27 | 2.89 |
| K562 | 7.22 | 14.92 | 6.11 | 15.46 | 10.5 |
| MCF-7 | 3.58 | 3.31 | 3.52 | N/A | 2.81 |

#### 6.2 (TL, ‖) Using the extended genes to calculate GWAS germline variant enrichment

We downloaded the GWAS variants from European ancestry from the GWAS Catalog https://www.ebi.ac.uk/gwas/ with keywords leukemia and breast cancer separately (version by 20180428). We removed less relevant phenotypes, such as BMR after therapy, to get a more confident set, which we called the tag SNPs.

After that, we downloaded the VCF file of the 1000 genomes Phase III data and extracted only the EUR population. We then extracted all LD SNPs with r^2 >=0.8 within 500Kb of the tag SNPs from PLINK (V1.90-beta5.3) as the start point for the enrichment analysis. We further removed the ones within known high LD regions. In the end, we had 14,401 and 4,308 variants for breast cancer and leukemia respectively for the analysis.

For the test regions, we defined the TSS regions, CDS regions, CDS+proximal (TFBS+RBP), and CDS+proximal+distal(enhancers) in MCF-7 and K562 to calculate the enrichment of GWAS variants. We defined the proximal and distal annotations in MCF-7 and K562 as the matched and unmatched annotations for breast cancer and the proximal and distal annotations in MCF-7 and K562 as the unmatched and matched annotations for leukemia. We used the VSE package in R to calculate the tag and LD SNP enrichment with default settings. We repeated the above procedure on the same annotation set focused on genes from the cancer gene consensus to test the enrichment of known cancer-associated gene sets. The raw results were provided in Table S 8-4.

#### 6.3 Using the extended genes to calculate somatic mutation burden

##### 6.3.1 (TL, ‖) Correcting the local 3mer context effect

We observed that BMR is significantly associated with local context effect in all cancer types up to several orders, which largely contributes to the mutation rate heterogeneity. Details are given in Figure S 6-2. For example, the average pooled mutation rate ranges from 1.58e04 to 2.92e03 (18.35 fold). The observed mutation has been plotted in the following radial plots for each cancer type. In general, G/C positions are more prone to mutations as compared to A/T positions, but the local context effect within G/C positions still has a strong effect. In addition, we also observed that the local context effect varies significantly across multiple cancer types. Hence, we separated the local 3mers to do BMR estimation separately in our analysis.

*Figure S 6-2. Local context severely confounds BMR in multiple cancer types*

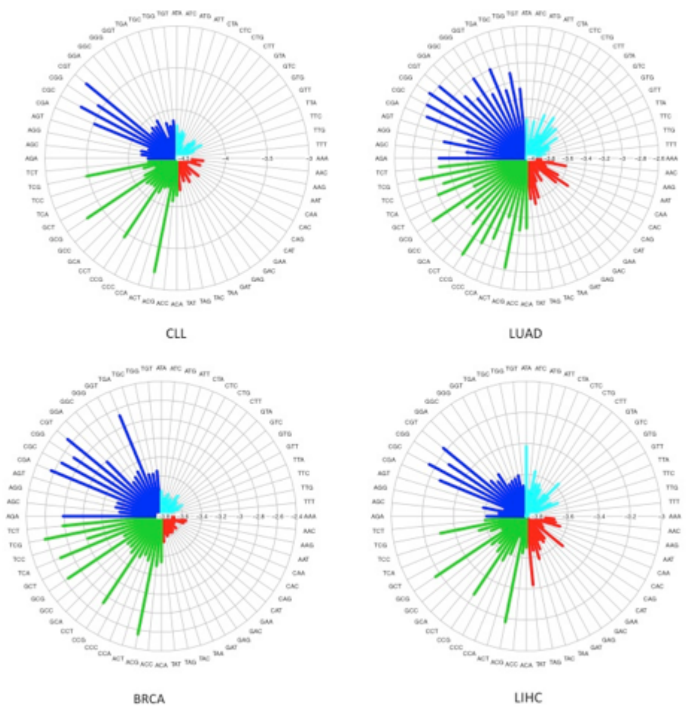

##### 6.3.2 (TL, ‖) BMR estimation through the negative binomial regression

Negative binomial regression has been used to estimate BMR in many previous papers. Here we extracted 2,066 histone modification and replication timing features from ENCODE and incorporated them into BMR estimation in a similar approach. We first did a PCA analysis on the covariate matrix. For each 3mer in each cancer types, we ordered the separate PCs according to its absolute correlation with the mutation counts per 1Mb bin. We start from the top 30 most correlated PCs and did a backward selection using Poisson regression first. In each step, we tested whether the overdispersion exists in the observed data after covariate correction using the AER package in R. If yes, we used the negative binomial regression instead. We estimated BMR for all 3mers in CLL, breast cancer, and liver cancer in the same way.

We observed a tendency for highly mutated cancers to be more likely to reject the Poisson model. For example, we plotted the overall mutation count under different 3mer context vs. the estimated overdispersion parameter (using the AER package) in R in the following figure. On one side, it is obvious that for 3mers with more variants, there is a tendency to introduce overdispersion and accept the Gamma-Poisson model (Figure S 6-3).

*Figure S 6-3. Mutation rate vs. overdispersion*

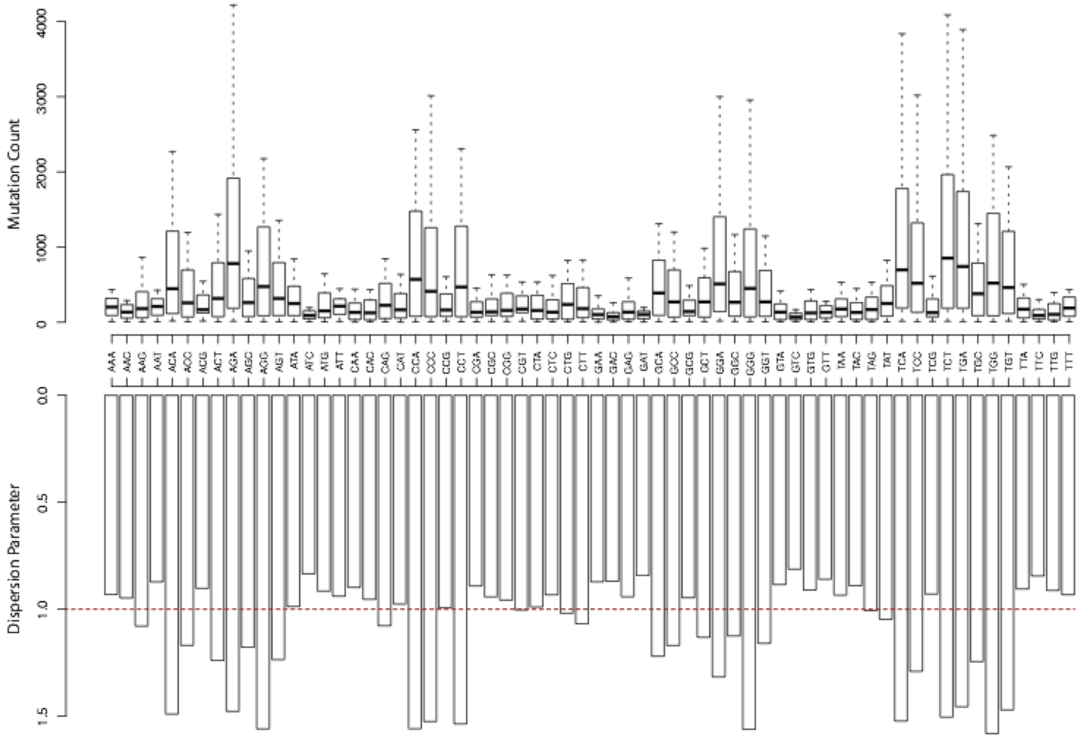

#### 6.3.3 (TL, ǁ) P-value calculation for somatic burden test

We tested the somatic burden of the TSS region (100bp before the TSS sites), CDS region, and our extended gene regions against the BMR estimated above. Specifically, for each of the discontinuous elements in an annotation, such as enhancers and gene bodies, we first find its nearest 1mb bin to extract the BMR parameters for all 3mers. Then, for each position, we count the number of mutations for each kmer in bin *i*. For any given annotation set, which might be a set of discontinuous segments in the genome, we calculated per segment mutation rate by assigning it to the nearest 1mb bins. Then for each local 3mer within a segment, we used either the trained Poisson or Negative binomial regression, depending on the result of the overdispersion test, to calculate mutation burden. We combined 3mers with the same parameter setting and used the convolution method to calculate p values for 3mers with different parameter settings. To remove bias from one single sample, we required that the somatically burdened annotation should have variants from at least two samples

##### 6.3.4 (TL, ‖) Q-Q plots of the extended genes

We provided the Q-Q plots of the p values using different annotation sets on various cancer types in Figure S 6-4.

*Figure S 6-4. Q-Q plot of the extended gene P values*

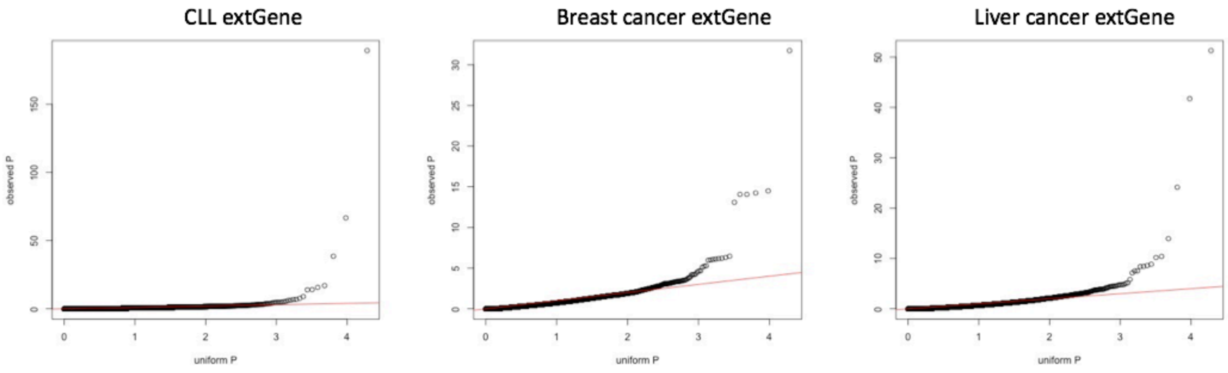

We also calculated the mutation burden on lincRNAs. We have found well-known cancer-associated lncRNAs to be burdened, such as NEAT1 in liver cancer and MALAT1 in breast cancer. Results and Q-Q plots are shown in Figure S 6-5.

*Figure S 6-5. Q-Q plots on lincRNAs*

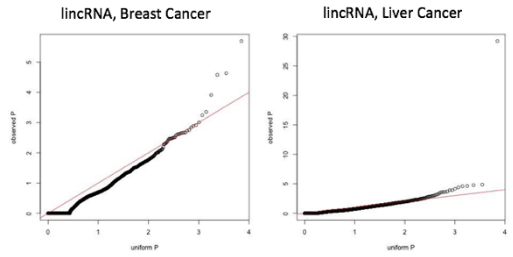

##### 6.3.5 (TL, ‖) More investigations of the highly mutated genes

Further investigations of the highly-mutated genes are needed to clarify their effect in cancer. For example, we found that there is a mutation hotspot near the first intron in BCL6 (Figure S 6-6). Off-target activity of the AID system has been previously reported at this location (see Pasqualucci et al., 1998[67] and Migliazza et al., 1995[68] for early reports of this association) and has been reported to be associated with translocation in Harris *et al,* 2001[69].

*Figure S 6-6. Mutation hotspot in BCL6*

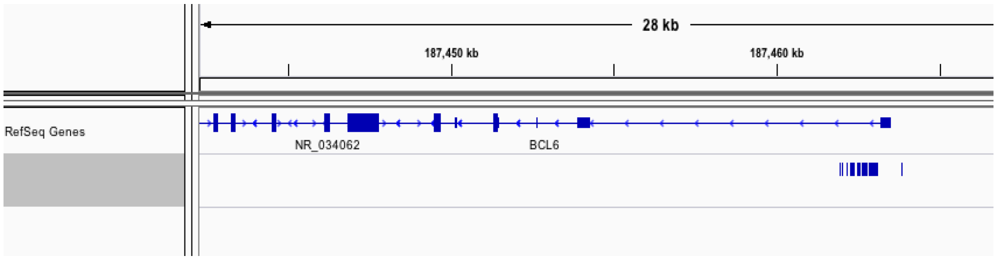

#### 6.4 (TL, Ⅱ) SV introduced oncogene activation

To give an intuitive example of the dynamics of the extended genes, we searched for the extended gene changes in tumor and normal that are caused by variants in the genomes. Specifically, we found an 130Kb heterozygous deletion in T47D cells by both optical mapping and whole genome sequencing (Figure S 6-7), about 45Kb downstream to the ERBB4 promoter, overlapping with the right-hand boundary of the TAD that encompasses ERBB4 gene. This deletion is also validated by PCR in comparison to HMEC cells, with the mutated allele 633bp and WT allele 320bp.

RNA-seq showed that ERBB4 gene expression is activated in T47D cells in compare with HMEC cells (5.188:0.002), while the gene itself does not show gain of copy (Figure S 6-8). Through 4C we observed a novel interaction in T47D cells between ERBB4 and a distal region from the next TAD, which is not present in HMEC cells. We therefore hypothesized that the heterozygous deletion disrupts the insulation of ERBB4 from distal regions and hence activates its expression on one allele. We narrowed down the disrupted TAD boundary by locating a conserved CTCF binding region (Figure S 6-9), and then tested the hypothesis by CRISPR editing with PX458 construct and a pair of sgRNA to excise the 88bp sequence on the wild-type allele in T47D cells that contains the CTCF binding motif. We validated the CRISPR editing by PCR and Sanger-sequencing (c). We check if it further enhances ERBB4 expression. Our result shows that the boundary disruption on the wild-type allele doubles ERBB4 expression, suggesting that the ERBB4 activation in T47D is at least in part due to the 130Kb deletion that disrupts its insulation.

*Figure S 6-7. Optical mapping detects the 130Kb deletion*

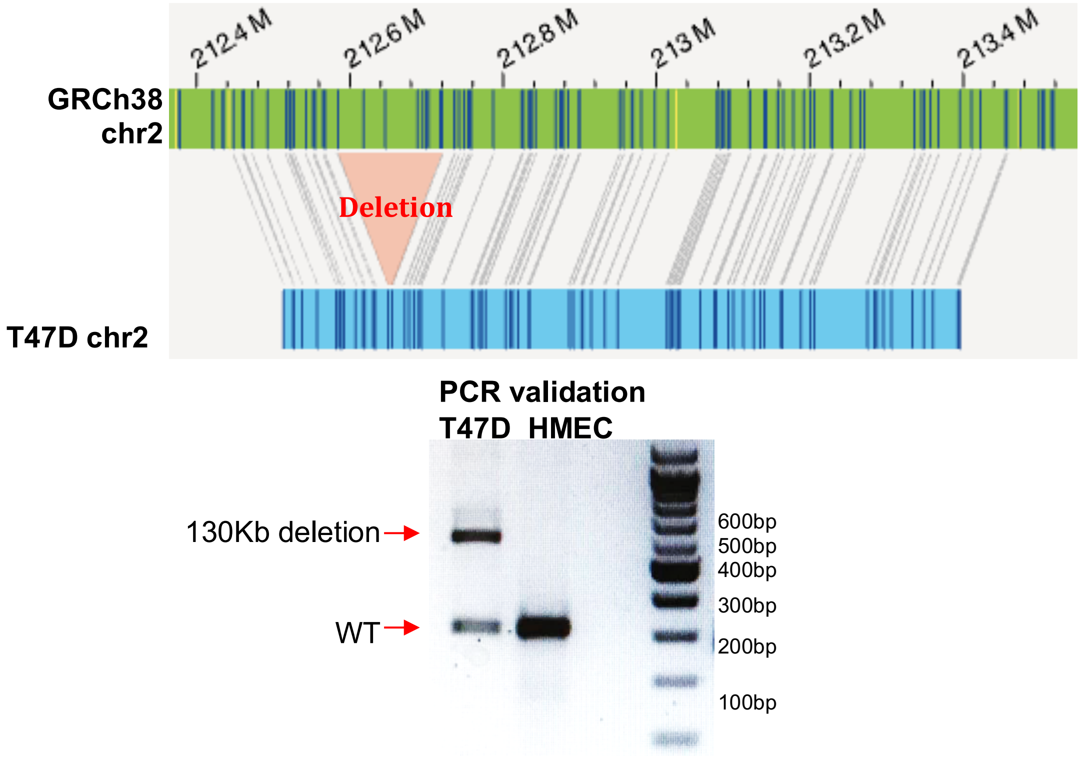

*Figure S 6-8. SV introduced oncogene activation*

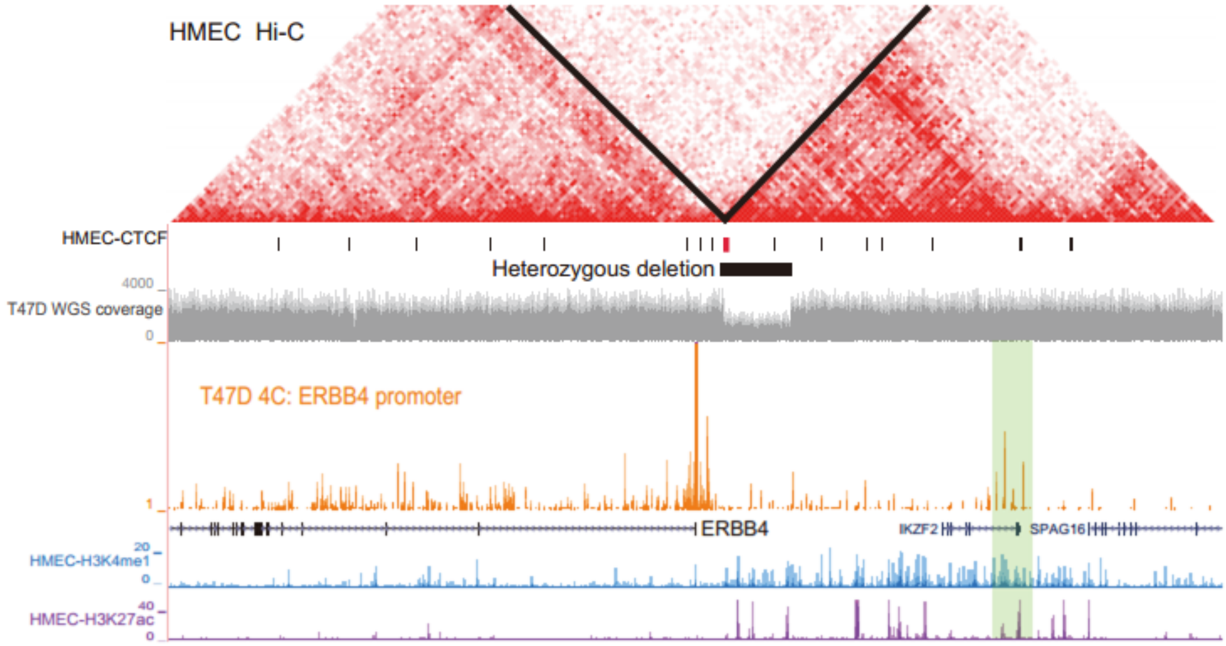

*Figure S 6-9. CTCF motif deleted*

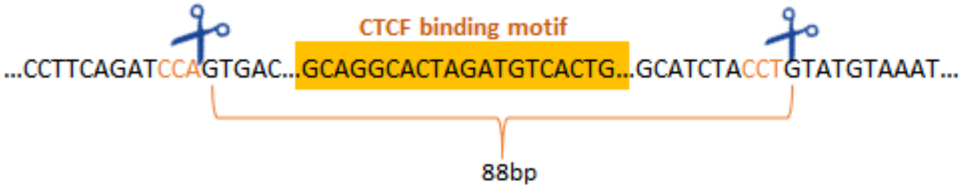

Specifically, CRISPR editing is carried out utilizing the PX458 vector. A pair of PX458 inserted constructs express the sgRNA that respectively targets the 5' and 3' end of the 86bp sequence were co-transduced to T47D cells by electroporation. T47D cells of the control group were transduced with non-targeting PX458 construct. Transduced cells were selected for GFP expression by fluorescence activated cell sorting. Selected cells were grown into single cell colonies and screened for fragment deletion by PCR. Positive colonies were expanded and ERBB4 gene expression is measured and compared by relative qPCR.

*Figure S 6-10. Vector used for the experiment*

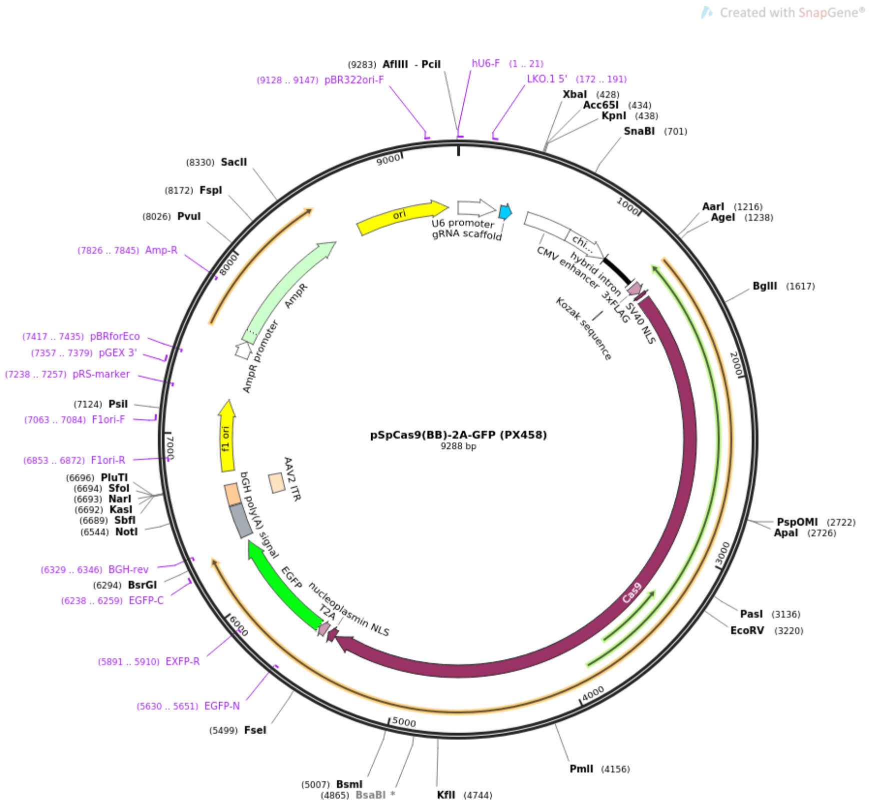

#### 6.5 (TL, ‖) Histone modification aggregation around SVs

We aggregated various ENCODE histone ChIP-seq data around SV breakpoints to see if chromatin marks are associated with SV events (Figure S 6-11). We observed an enrichment of activating H4K20me1 marks near somatic breakpoints in K562 but not in GM12978. Similar pattern was also observed with H3K27me3 mark.

*Figure S 6-11. Aggregation of histone marks around somatic SV breakpoints in K562*

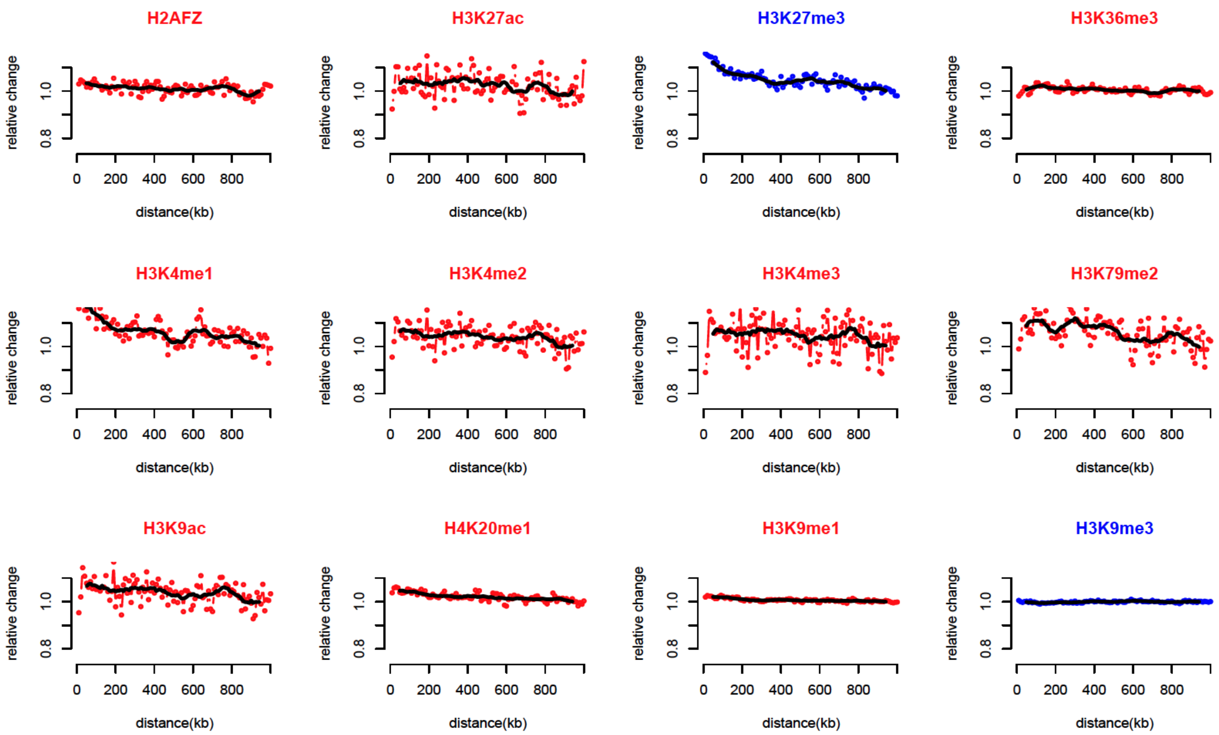

##### 6.5.1 (TL, ‖) Construction of the somatic deletion set in K562

We first merged all deletions with 50% reciprocal overlapping from two callsets: Zhou et al[62] and Dixon et al[63]. For merged SVs and SVs supported by multiple callers, we took the maximum boundaries. Then we filtered the set against 1000 Genome Project phase 3 call set to remove possible germline variants. In total, we got 3,147 deletions in our merged deletion set.

##### 6.5.2 (TL, ‖) Aggregation of histone modification markers around breakpoints

We used 10kb bins to tile 1MB upstream of 5’ breakpoints and 1MB downstream of 3’ breakpoints. Bins that overlapped with the blacklist or fell out of the chromosome boundaries were removed. Then we calculated the average normalized histone fold change signal in each bin (ENCSR000AKI and ENCSR000AKQ). We then combined all breakpoints, aligned the bins and took the mean of bins of the same distance to breakpoints. The mean is further normalized by the mean of the furthest ten bins. We fitted trend lines using cubic smooth spline with the degree of freedom of 5.

*Figure S 6-12. H4K20me1 aggregation around deletion breakpoint*

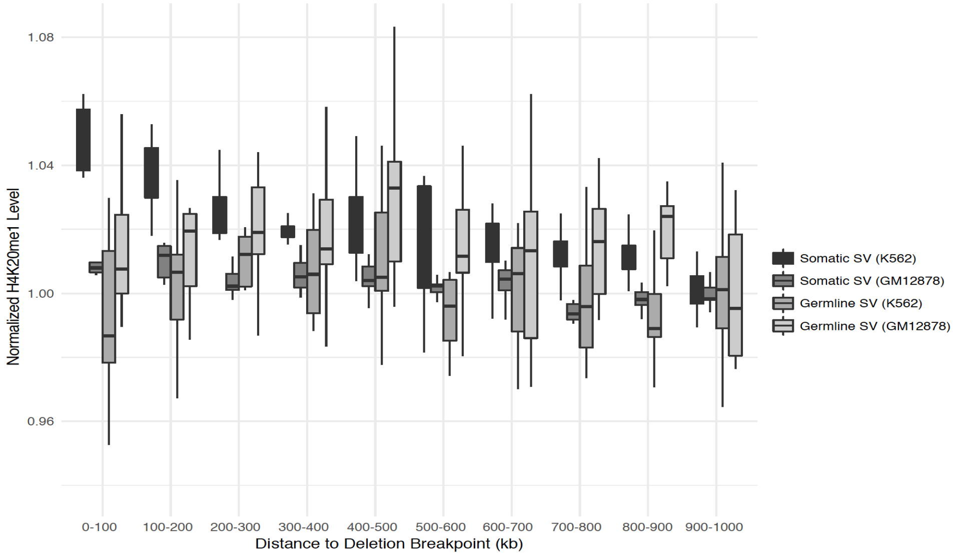

##### 6.5.3 (TL, ‖) ENCODE SV calls

The ENCODE SV data consists of many cell types and different kinds of SVs. These SVs are from the WGS calls. The summary of ENCODE SV calls can be found below in Table S 6-2.

*Table S 6-2. Summary of ENCODE SV calls*

**ENCODE SV Calls**

| Cell | Deletion | Inversion | Insertion | Duplication | Inter-Chr Transloc. | Intra-Chr |
|---|---|---|---|---|---|---|
| A549 | 3654 | 27 | 715 | 191 | 85 | 210 |
| Caki2 | 3202 | 29 | 863 | 214 | 192 | 290 |
| K562 | 3223 | 31 | 702 | 206 | 111 | 240 |
| LNCAP | 4151 | 62 | 840 | 0 | 247 | 224 |
| NA12878 | 3473 | 66 | 760 | 41 | 3 | 278 |
| NCI-H460 | 2951 | 37 | 699 | 224 | 112 | 228 |
| PANC-1 | 3165 | 48 | 887 | 239 | 199 | 313 |
| SK-N-MC | 3036 | 49 | 712 | 234 | 139 | 278 |
| T47D | 2945 | 27 | 736 | 224 | 125 | 234 |

##### 6.5.4 (TL, ‖) Elevated mutation rate around SVs

We compared the SNV/InDel density near the SV boundaries in strictly matched ENCODE cell lines and found that there are noticeably elevated SNV/InDel rates around SVs.

*Figure S 6-13. Elevated mutations rates around SVs*

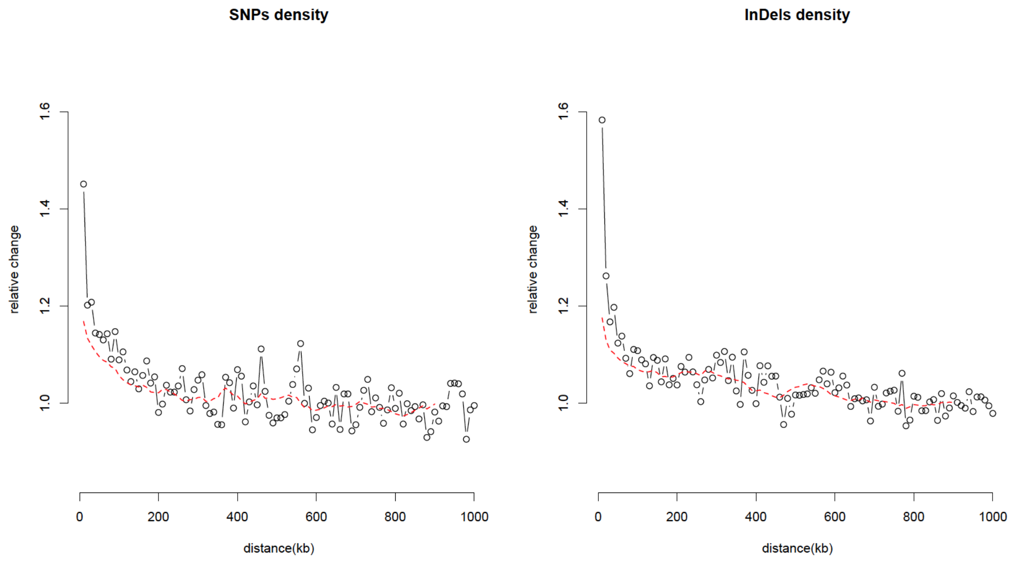

#### 6.6 (TL, ∦) Direct SV effects on gene expressions

We have shown in Figure S 6-14 several examples of SVs near promoter regions that may affect gene expression.

*Figure S 6-14. Direct SV effects on gene expressions*

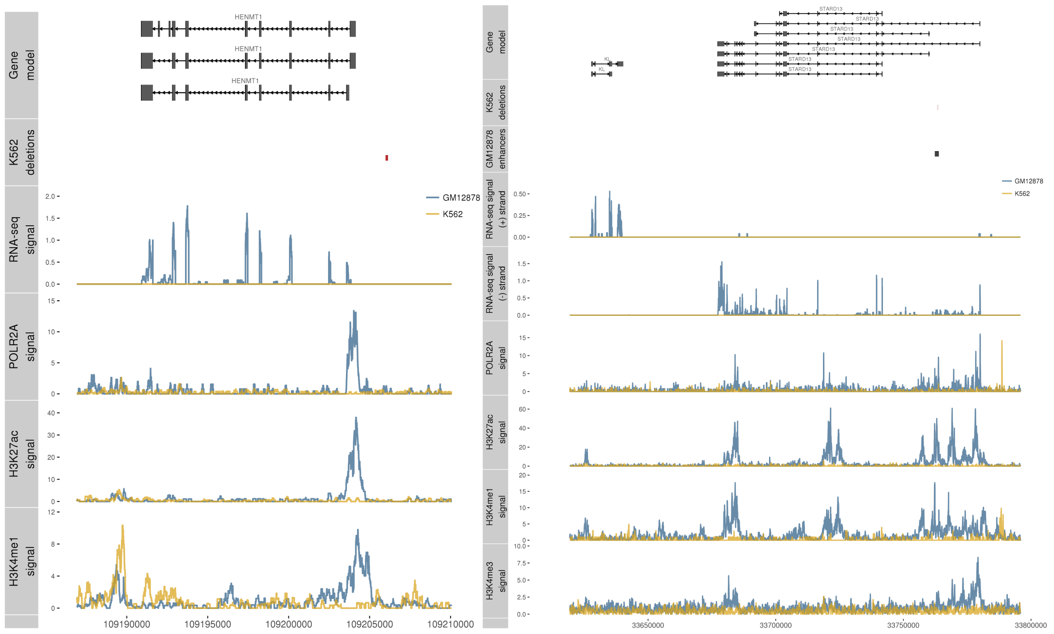

### 7 More details about “Using the ENCODEC resource to prioritize SNVs”

#### 7.1 (HL, ‖) Step-wise prioritization workflow

The description of the regulatory network and mutation recurrence analysis provides a way to prioritize key genomic features associated with cancer. Here we proposed a step-wise scheme to prioritize regulators, elements, and SNVs for small-scale validations. First, we start by searching for key regulators that frequently rewire, locate in network hubs or on top of the network hierarchy, or significantly drive expression changes in cancer. We then prioritize functional elements that associate with top regulators, undergo large regulatory and chromatin changes, or (most importantly) are highly mutated in tumors. Finally, on a nucleotide level, we can pinpoint impactful SNVs for small-scale functional characterization by their ability to disrupt or create specific binding sites, or which occur in positions of particularly high conservation or chromatin changes.

*Figure S 7-1 Step-wise prioritization workflow*

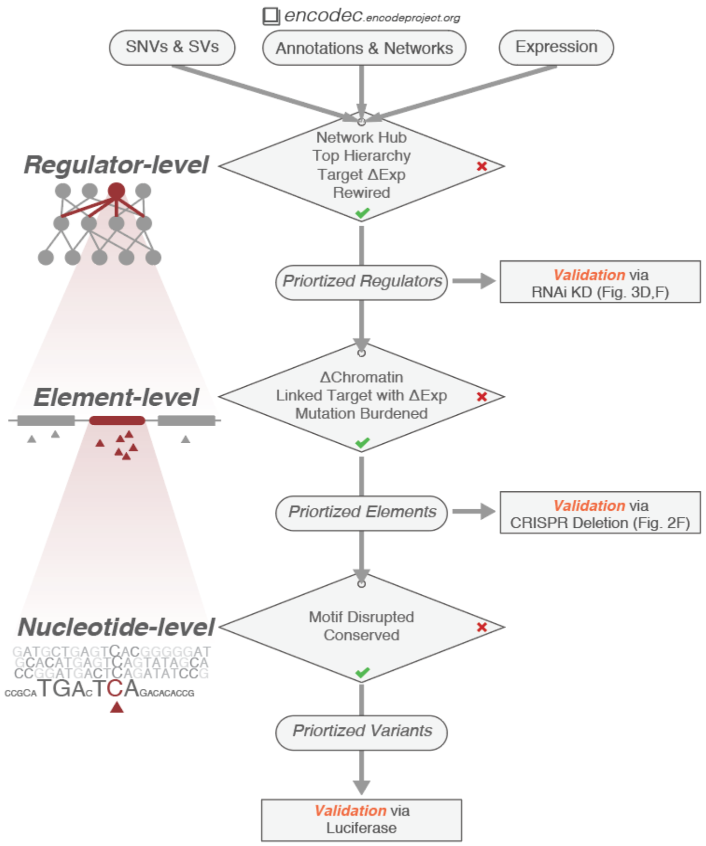

To further illustrate the difference of our methods with previous efforts, we made Table S 7-1 to show the distinct aspect of our prioritization scheme.

*Table S 7-1 Difference of the workflow with previous methods*

| Goal | ENCODEC | FunSeq | CADD | GWAVA | FATHMM-MKL | TFRank |
|---|---|---|---|---|---|---|
| Variant Pri. | ✓ | ✓ | ✓ | ✓ | ✓ | |
| Element Pri. | ✓ | | | | | ✓ |
| Regulator Pri. | ✓ | | | | | ✓ |

*Table S 7-2 Difference of the dataset used with previous methods*

| Levels | Feature | ENCODEC | FunSeq |
|---|---|---|---|
| Raw data | eCLIP | ✓ | x |
| | STARR-seq | ✓ | x |
| | Hi-C | ✓ | x |
| | ChIA-PET | ✓ | x |
| Network | Exp. Reg. power | ✓ | x |
| | Network rewiring | ✓ | x |
| | Network hub | ✓ | ✓ |
| | Network hierarchy | ✓ | x |
| | Linked gene diff. exp. | ✓ | x |
| Element | Somatic burden | ✓ | ✓ |
| | Δchromatin, exp. | ✓ | x |
| Nucleotide | conservation | ✓ | ✓ |
| | Motif disruption | ✓ | ✓ |

#### 7.2 (TL, ‖) Motif analysis using MotifTools (D-score)

At the nucleotide level, we calculated motif breaking score, called the D-score to measure disruptive-ness or deleterious-ness. Motif disruption score was calculated based on the difference between sequence specificities of reference to alternative sequence

We used the changes of PWMs introduced by a variant to quantify the motif disruptiveness effect through MotifTools (https://github.com/gersteinlab/MotifTools. Specifically, we defined the disruption score, $D_{score}$, as in equation (7-1) to represent the difference between sequence specificities to an alternative sequence.

$$D_{score} = M_{ref} - M_{alt} = -10 \times \log_{10}\left(\frac{P_{ref}}{P_{alt}}\right) \quad (7\text{-}1)$$

where $P_{ref}$ and $P_{alt}$ are the PWM scores from the reference and alternative allele. Here, the motif scores for reference and alternate sequences are given as

$$M_{ref} = -10 \times \log_{10}(P_{ref}),\ M_{alt} = -10 \times \log_{10}(P_{alt}) \quad (7\text{-}2)$$

A positive D-score denotes a variant that decreases the likelihood that a TF will bind the motif (motif-break), and negative D-score denotes a variant that increases the likelihood that a TF will bind the motif (motif-gain). For assessing D-score, uniform nucleotide background was assumed (A:C:G:T=1:1:1:1) and the p-value threshold of 1e-3 was used. For position weight matrix (PWM), JASPAR TF profiles (2016 core non-redundant vertebrates, http://jaspar.genereg.net/html/DOWNLOAD/JASPAR_CORE/pfm/nonredundant/pfm_vertebrates.txt) were used and variants that affect multiple TF binding profiles were averaged over all D-scores. More details about the tool and code can be found at https://github.com/gersteinlab/MotifTools. Somatic variants were further prioritized using their conservation score (high positive GERP score).

*Figure S 7-1. Schematic of MotifTools output*

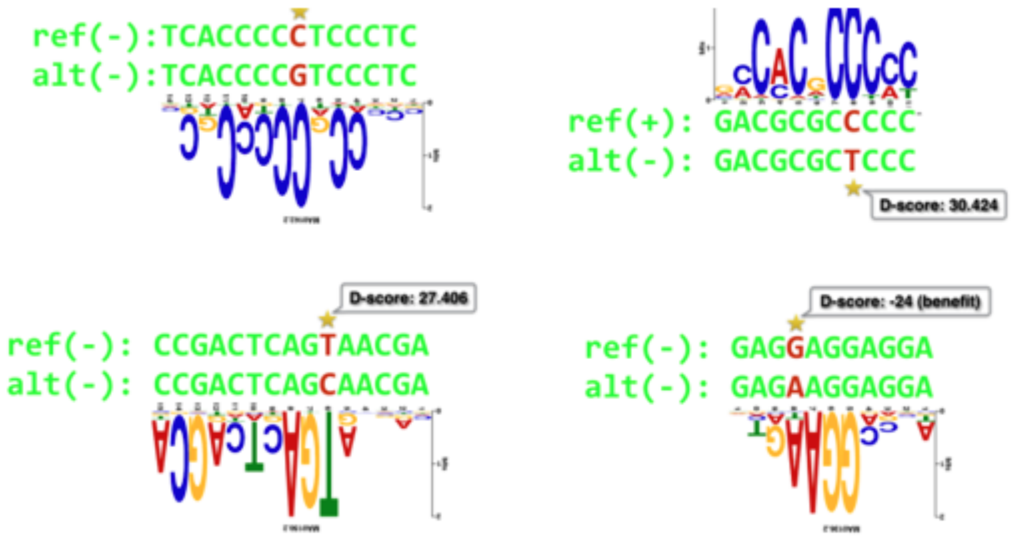

*Table S 7-3. Validated mutations in MCF-7 and luciferase assay tested region*

| Validation ID | CHR | Variant POS | REF | ALT | Element Start | Element End | Note |
|---|---|---|---|---|---|---|---|
| Failed 01 | chr16 | 85604242 | C | G | 85603992 | 85604491 | |
| Candidate 01 | chr21 | 27541982 | G | A | 27541732 | 27542231 | |
| Failed 02 | chr8 | 21541726 | A | G | 21541476 | 21541975 | |
| Candidate 02 | chr17 | 38474408 | C | G | 38474158 | 38474657 | |
| Candidate 03 | chr20 | 43971343 | G | C | 43971093 | 43971592 | |
| Candidate 04 | chr7 | 1598567 | C | T | 1598317 | 1598816 | |
| Candidate 05 | chr20 | 58563412 | C | T | 58563162 | 58563661 | Figure 6C |

| | | | | | | |
|---|---|---|---|---|---|---|
| Candidate 06 | chr7 | 150759483 | C | G | 150759233 | 150759732 |
| Candidate 07 | chr7 | 5596005 | T | G | 5595755 | 5596254 |
| Candidate 08 | chr6 | 134700462 | G | T | 134700212 | 134700711 |

#### 7.3 (TL, ‖) Details about other test regions

Using the prioritization scheme, 10 candidate regions containing disruptive variants were selected for validation in MCF-7 cell line. We present the genomic locations of candidate regions relative to histone signals and nearby genes as below.

*Figure S 7-2. Validated candidate region 1*

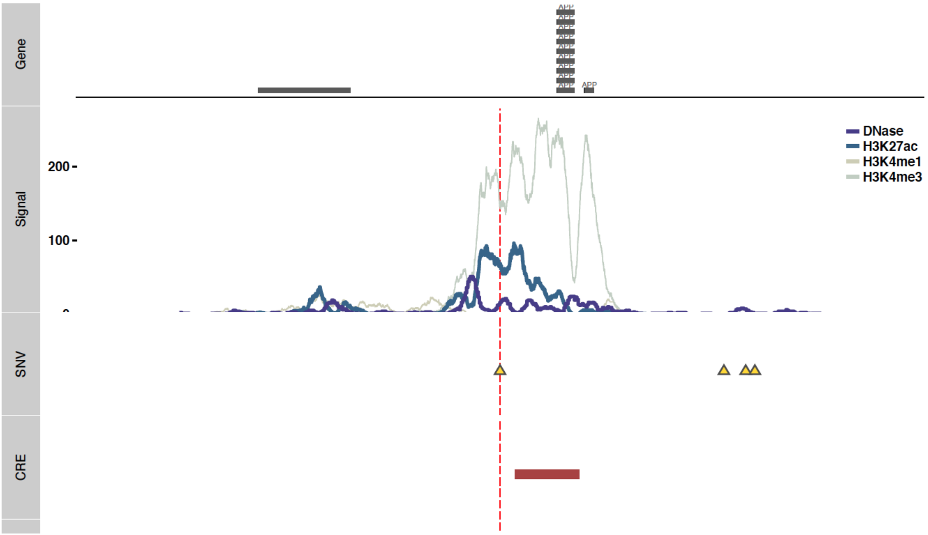

*Figure S 7-3. Validated candidate region 2*

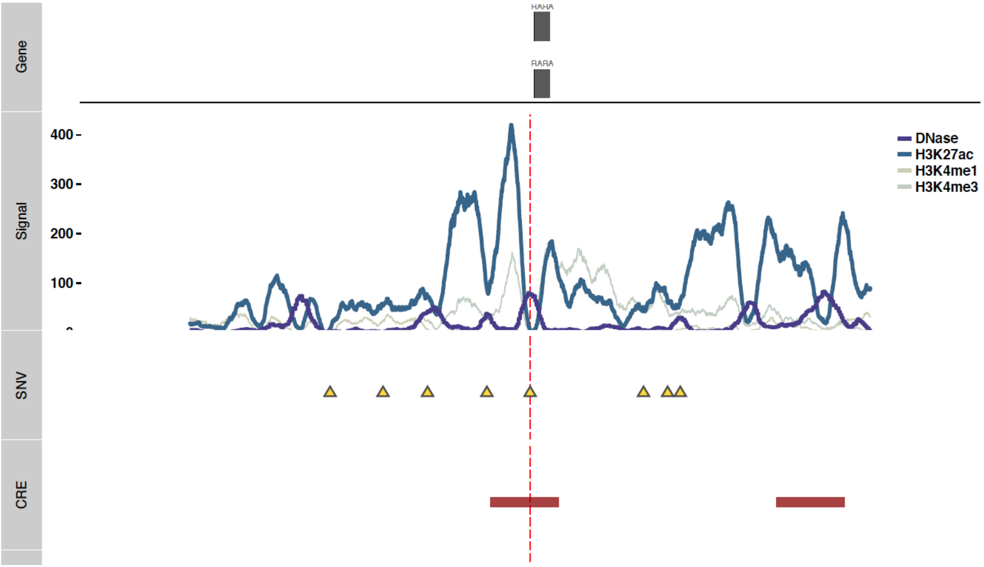

*Figure S 7-4. Validated candidate region 3*

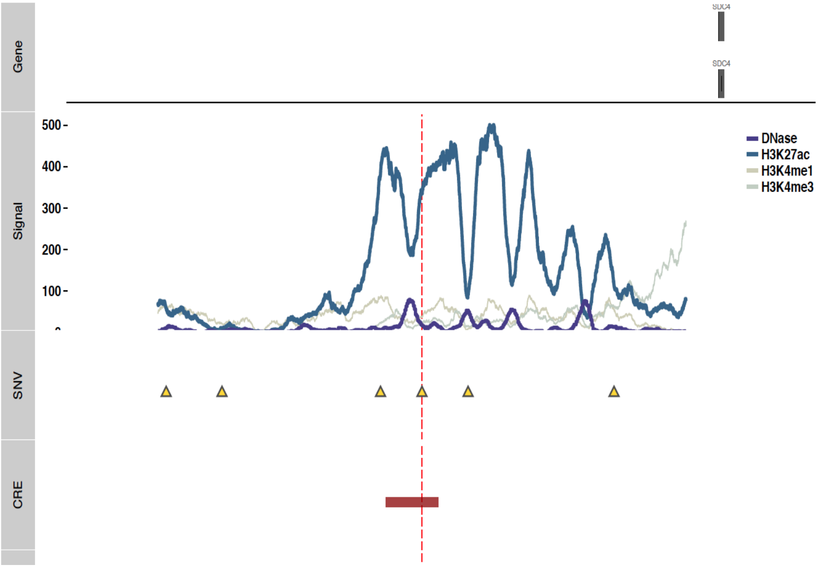

*Figure S 7-5. Validated candidate region 4*

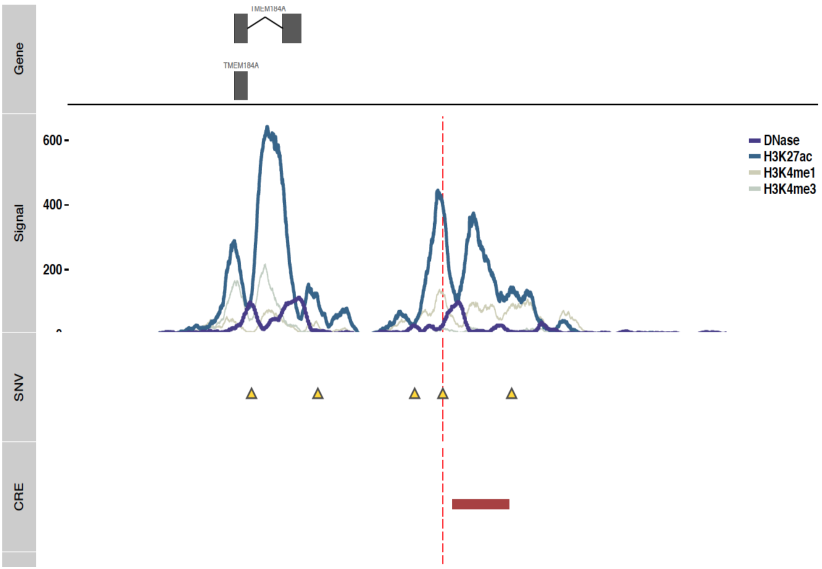

*Figure S 7-6. Validated candidate region 5*

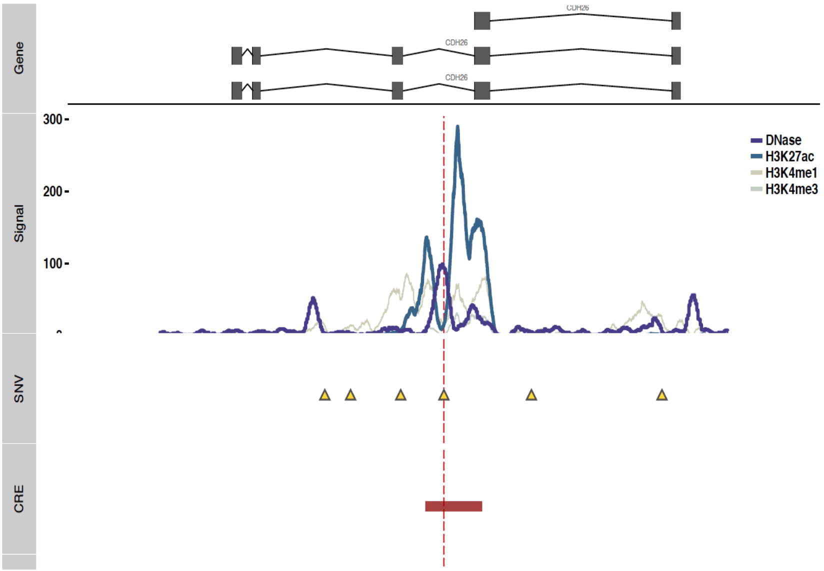

*Figure S 7-7. Validated candidate region 6*

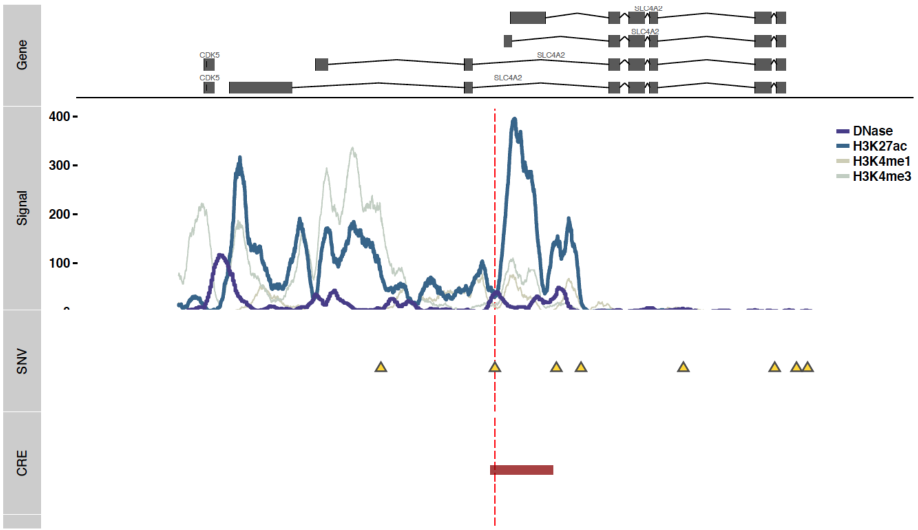

*Figure S 7-8. Validated candidate region 7*

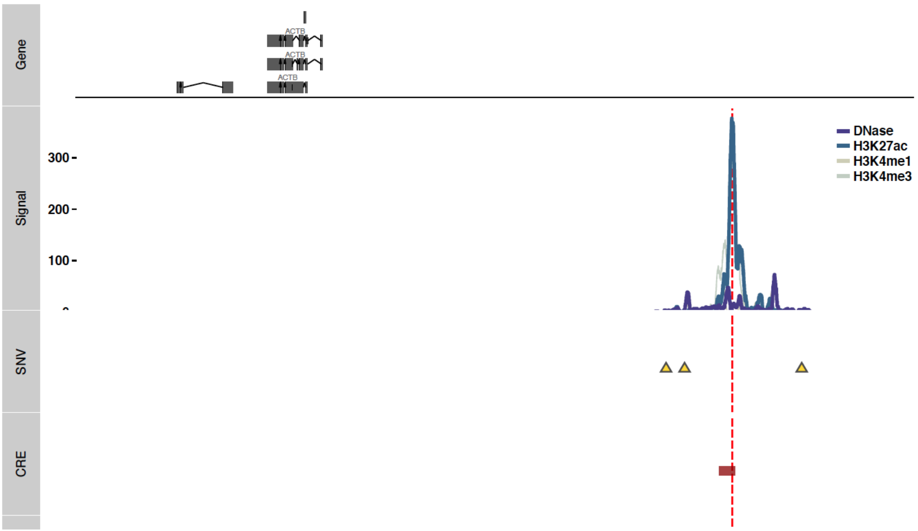

*Figure S 7-9. Validated candidate region 8*

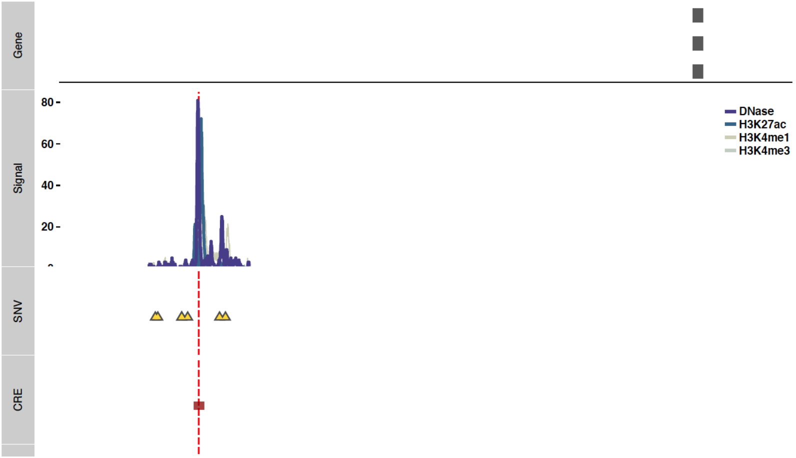

#### 7.4 (TL, Ⅱ) Experiment Details about SNV validation

Each regulatory region (both wild and mutant types) was separately synthesized. Enhancer regions were designed in such a fashion where based on the candidate SNV site, 250bp upstream and 250bp downstream were included for each enhancer region. These regions were then cloned into the pGL4.23[luc2/minP] vector (Promega, Cat# E841A). Each candidate region was placed upstream of the minP promoter to determine the effect of each putative enhancer region on luciferase expression. 100ng of each candidate construct and 100ng of Nano-luc control were co-transfected into MCF-7 cells (5,000 cells per well in DMEM media containing 10% FBS and 1% Penicillin-Streptomycin antibiotic) using the Lipofectamine 3000 reagent (Thermo Fisher, Cat# L3000001) according to the manufacturer's instructions. Cells were incubated for 48 hours before their luciferase signal was read using the Promega Nano-Glo luciferase kit (Promega, Cat# N1521) according to the manufacturer's instructions.

*Figure S 7-10. Schematic of SNV validation using Luciferase assay*

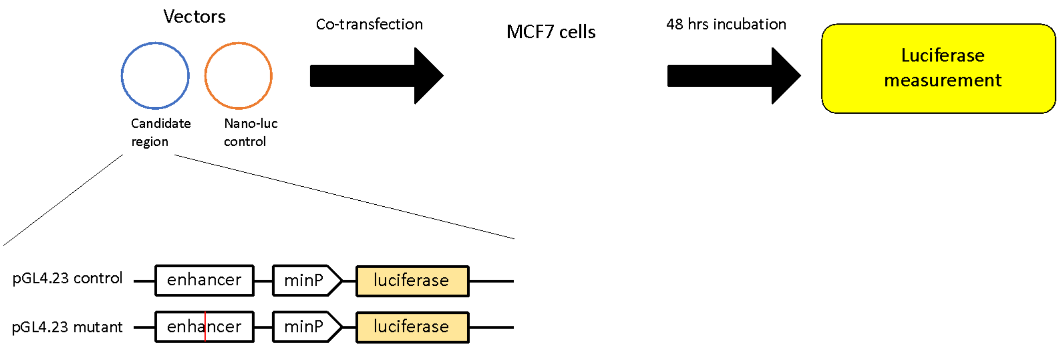

*Table S 7-4. Details of SNV replication technical replicate 1*

| | Normal Rep 1 | Normal Rep 2 | Normal Rep 3 | Mutant Rep 1 | Mutant Rep 2 | Mutant Rep 3 |
|---|---|---|---|---|---|---|
| Background | 831 | 388 | 416 | 2623 | 1296 | 1065 |
| 2 | 7698 | 5193 | 6893 | 161889 | 132344 | 179837 |
| 4 | 587863 | 778963 | 603304 | 465322 | 408546 | 460135 |
| 5 | 10281 | 16083 | 17192 | 40103 | 63770 | 48912 |
| 6 | 39090 | 20019 | 23419 | 7614 | 6760 | 4959 |
| 7 | 15039 | 18873 | 13468 | 57945 | 47666 | 59931 |
| 8 | 117702 | 115358 | 150245 | 189131 | 295907 | 247173 |
| 9 | 26775 | 30804 | 34042 | 58424 | 104433 | 27587 |
| 10 | 21705 | 22249 | 17162 | 107077 | 31005 | 76174 |
| Empty | 61423 | 87225 | 46835 | 774 | 789 | 1111 |
| Background | 562 | 1461 | 748 | 4582 | 967 | 473 |
| Background | 238 | 500 | 395 | 857 | 635 | 921 |

*Table S 7-5. Details of SNV replication technical replicate 2*

| | Mutant Rep 1 | Mutant Rep 2 | Mutant Rep 3 | Normal Rep 1 | Normal Rep 2 | Normal Rep 3 |
|---|---|---|---|---|---|---|
| Background | 11852 | 13823 | 14402 | 15111 | 13245 | 9858 |
| 2 | 1922952 | 1854116 | 1882977 | 2326518 | 1637299 | 1927383 |
| 4 | 1969924 | 1947206 | 2088052 | 1606057 | 1133593 | 1246025 |
| 5 | 1396532 | 1408962 | 1879464 | 2110566 | 1890350 | 1594218 |
| 6 | 1756884 | 1798060 | 1859447 | 1825321 | 1597249 | 1658538 |
| 7 | 1884514 | 2197614 | 2393865 | 2124074 | 1385636 | 1888050 |
| 8 | 1695866 | 1711603 | 1488882 | 2405882 | 1487463 | 1516048 |
| 9 | 1715909 | 1943040 | 1916404 | 2058790 | 1385673 | 1241105 |
| 10 | 1771446 | 1498757 | 2030086 | 1736458 | 985080 | 1237019 |
| Empty | 2575562 | 2699389 | 2494020 | 22537 | 10758 | 6625 |
| Background | 12437 | 14855 | 12235 | 7338 | 4629 | 2613 |
| Background | 3835 | 4041 | 4182 | 2990 | 1698 | 1009 |

There were 2 biological replicates and 3 technical replicates performed for each candidate region, totaling to 6 replicates. Statistical significance was tested using t-test and annotated in Figure 6B, using asterisks to communicate significance thresholds. In summary, *: 0.01<P<=0.05; **: 0.01<P <=0.001; ***: P<0.001.

*Table S 7-6. Validation p-values*

| Validation ID | P-value |
|---|---|
| Candidate 01 | 0.002555, ** |
| Candidate 02 | 0.9191, NS |

| | |
|---|---|
| Candidate 03 | 0.4828, NS |
| Candidate 04 | 0.004157, ** |
| Candidate 05 | 5.454e-05, *** |
| Candidate 06 | 0.2659, NS |
| Candidate 07 | 0.4352, NS |
| Candidate 08 | 0.005305, ** |

#### 7.5 (TL, ‖) Details about siRNA RNA-seq experiments

MCF-7 cells were seeded at an initial density of 2 x $10^5$ cells at basal media conditions (DMEM media supplemented with 10% FBS and 1% penicillin/streptomycin). After 24 hours, cells were transfected with siRNA control (Santa Cruz, Cat#: sc-37007) or siRNA targeting MYC (Santa Cruz, Cat# sc-29226) using the Viafect reagent (Promega, Cat#: E4981) in biological triplicate according to the manufacturer's instructions. Cells were harvested 72 hours later using the Trizol Reagent (Thermo Fisher, Cat#: 15596026), and RNA was isolated using the Direct-zol RNA Miniprep Plus kit (Zymo Research, Cat#: R2072). RNA-seq libraries were prepared using the TruSeq Stranded Total RNA Library Prep Kit (Illumina, Cat#: RS-122-2201). RNA-seq library quality was assessed using Bioanalyzer and Qubit analysis before it was subjected to deep sequencing on the HiSeq 4000.

We used RNA-STAR (version 2.3.0) to map the reads to the hg19 genome with Gencode v19 as annotation input for junction mapping. Default settings were used for the read mapping. After mapping, we used Cufflinks (version 2.0.2) for the gene and transcription level quantifications for both knockdown and control experiments.

### 8 Other related summary tables

*Table S 8-1. Summary of the ENCODEC resource*

| Contribution | | Description | New Method | Release | Detail details |
|---|---|---|---|---|---|
| Raw data | | Histone modification | -- | Text | 2015 ChIP-seq |
| | | Replication timing | -- | Text | 51 Repli-seq/ChIP |
| SV/SNV | | Cell line genome variations (in 9 cell lines) | -- | VCF | WGS, Bionano, Hi-C, Repli-seq |
| Annotation | | Enhancer | MatchedFilter | BED | ChIP-seq, DNase-seq |
| | | Enhancer | ESCAPE | BED | **STARR-seq** |
| | | Enhancer-gene linkage | JEME+Hi-C | BED | ChIP-seq, RNA-seq, DNase-seq, ChIA-PET, **Hi-C** |
| | | **Extended Gene** | -- | BED | eCLIP, STARR-seq, ChIP-seq, DNase -seq, ChIA-PET, Hi-C |
| Network | Proximal | **RBP-gene** (tissue-specific & universal) | -- | Text | **eCLIP** for 167 RBPs |
| | | TF-gene (tissue specific & universal) | -- | Text | ChIP-seq for 1,863 TFs |
| | | TF-gene (tissue specific, imputed) | -- | Text | 678 DNase-seq |
| | Distal | **TF-enhancer-gene** (tissue specific) | -- | Text | 486 TFs |

*Table S 8-2. List of experimental validations*

| **Assay used** | shRNA + RNA-seq | shRNA + RCR | CRISPR | Luciferase assay |
|---|---|---|---|---|
| **Validation Level** | Regulator level | Regulator crosstalk | Element level | Nucleotide level |
| **Evidence for** | MYC and SUB1 upregulate target gene expression | MYC and SUB1 bind to the promoter and 3'UTR to upregulate co-regulate several cell-cycle-related oncogenes | Oncogene ERBB4 activation through enhancer-gene linkage gain in T47D due to TAD merge | Mutations that change enhancer activity levels |
| **Associated Fig.** | Fig. 1 D & F | Fig. 1 F | Fig. 5 B | Fig. 6 B & C |
| **Associated. Supp. section** | Supplementary section 3.5 | Supplementary section 3.5 | Supplementary section 6.4 | Supplementary section 7.4 |

*Table S 8-3. Summary of discoveries*

| Level | Claims | Associated Figure |
|---|---|---|
| Network | Construction and application of RBP network in cancer | Fig. 1 A & B |
| | Discovery of previously unidentified RBP to drive cancer specific gene expression pattern and report its value to predict patient survival | Fig. 1 F |
| | Cross talk of transcriptional and post-transcriptional regulators to initiate oncogene expression | Fig. 1 E-F |
| | Network rewiring analysis using gene and gene community-based methods | Fig. 2 & 3 |
| | Stemness movement on the normal-to-tumor trajectory, and its validation through TF knockdowns | Fig. 4 |
| Annotation | Gene-centric noncoding regulome changes and effects on gene expression | Fig. 5 A-B |

We listed the other statistical supports of claims in

| | Feature | Biological Significance | Statistical Support |
|---|---|---|---|
| Figure 1 | Network Hierarchy: TF-RBP crosstalk | More TF-RBP interactions (crosstalk) at the top-level hierarchy as compared to the middle and bottom level | Wilcoxon rank sum test, p-value = 3.381e-16 (3vs1), p-value=1.221e-09 (2vs1) |
| | Network Hierarchy: target expression | TF and RBP at the top level of network hierarchy had larger correlation to drive target gene expression change | Wilcoxon rank sum test, p-value < 2.2e-16 |
| | Knockdown: MYC | Target genes of MYC affected after KD | Wilcoxon rank sum test, p-value ≤ 2.2e-16 |
| | Knockdown: SUB1 | Target genes of SUB1 affected after KD | Wilcoxon rank sum test, p-value ≤ 3.1e-25 |
| | Survival analysis: MYC | High expression of MYC results in shorter survival | Cox proportional hazards regression, p-value ≤ 0.03 |
| | Survival analysis: SUB1 | High expression of SUB1 results in shorter survival | Cox proportional hazards regression, p-value ≤ 0.007 |
| | Decay rate: SUB1 | Targets of SUB1 has longer decay rate | Wilcoxon rank sum test, two-sided, p-value = 6.96e-14 |
| Figure 2 | Network Hierarchy: target expression | TF at the top level of network hierarchy had larger correlation to drive target gene expression change | Wilcoxon rank sum test, two-sided, P=0.0002268 (3vs1), P=0.0084 (2vs1), P=0.01079 (3vs2) |
| | Network Hierarchy: percent burdened TFBS | TF at the bottom level of hierarchy had more burden in TFBS | Not Significant. See Figure S 8-1 |
| Figure 4 | Stemness | Transcriptome became more similar to normal state upon knocking down oncogenes, compared to TSGs | P-value = 0.04 |
| Figure 5 | SV Breakpoint Analysis | H4K20me1 occurs preferentially around SV breakpoints | Wilcoxon rank sum test, one-sided, somatic SV in K562: p-value = 3.265e-10, GM12878: p-value = 2.542e-08, germline SV in K562 & GM12878: NS |
| | BMR estimation using ENCODE Data | More accurate using cell and tissue data from 2068 features | Regression performance better in cell & tissue than just cell or tissue data. |
| | Extended Gene: GWAS | GWAS SNPs more easily identifiable | P values directly from VSE package. Please check details for table below |
| | Extended Gene: Expression Stratification | Stratify patients' gene expression based on mutation status, i.e. well-known cancer gene SRSF2 | Wilcoxon signed-rank test. one-sided, p-value = 0.002 |
| Figure 6 | Validation | Prioritized candidate region #5 validation via luciferase assay | Student's t-test, two-sided, p-value = 5.454e-05 |

*Table S 8-4. Detailed enrichment for GWAS analysis*

| Region | Enrichment | Adjusted P-value | cancer | gene |
|---|---|---|---|---|
| TSS | 0.577960429 | 1 | Breast Cancer | All |
| CDS | 1.924311399 | 0.502656158 | Breast Cancer | All |
| nonMatch-Prox | 3.76098321 | 0.001714928 | Breast Cancer | All |
| Match-Prox | 3.724391587 | 0.002979011 | Breast Cancer | All |
| nonMatch-ExtGene | 4.13011297 | 0.000372968 | Breast Cancer | All |
| Match-ExtGene | 4.665158032 | 3.05E-05 | Breast Cancer | All |
| TSS | 1.601269617 | 1 | Breast Cancer | CGC |
| CDS | 3.157494607 | 0.015313093 | Breast Cancer | CGC |
| nonMatch-Prox | 4.088659885 | 0.000571839 | Breast Cancer | CGC |
| Match-Prox | 3.557659194 | 0.003777868 | Breast Cancer | CGC |
| nonMatch-ExtGene | 3.952046556 | 0.000787999 | Breast Cancer | CGC |
| Match-ExtGene | 5.019338192 | 8.96E-06 | Breast Cancer | CGC |
| TSS | 1.948784655 | 0.559952222 | Leukemia | All |
| CDS | 2.814489866 | 0.048567867 | Leukemia | All |
| nonMatch-Prox | 2.866434604 | 0.045261771 | Leukemia | All |

| | | | | |
|---|---|---|---|---|
| Match-Prox | 4.103573457 | 0.000457792 | Leukemia | All |
| nonMatch-ExtGene | 3.161692366 | 0.010989852 | Leukemia | All |
| Match-ExtGene | 4.38379163 | 0.00016207 | Leukemia | All |
| TSS | 1.352070515 | 1 | Leukemia | CGC |
| CDS | 2.056541629 | 0.327800164 | Leukemia | CGC |
| nonMatch-Prox | 2.172706287 | 0.273031432 | Leukemia | CGC |
| Match-Prox | 2.706772865 | 0.054321113 | Leukemia | CGC |
| nonMatch-ExtGene | 1.482965519 | 1 | Leukemia | CGC |
| Match-ExtGene | 4.221439681 | 0.000298694 | Leukemia | CGC |

*Figure S 8-1. network hierarchy vs. fraction of burdened binding sites for TFs*

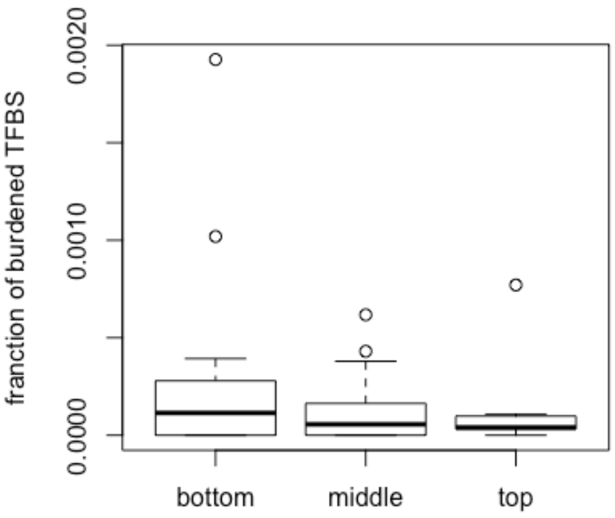

Reference

1 Greenbaum, D., Rozowsky, J., Stodden, V. & Gerstein, M. Structuring supplemental materials in support of reproducibility. *Genome Biol* **18**, 64, doi:10.1186/s13059-017-1205-3 (2017).

2 Bamford, S. *et al.* The COSMIC (Catalogue of Somatic Mutations in Cancer) database and website. *Br J Cancer* **91**, 355-358, doi:10.1038/sj.bjc.6601894 (2004).

3 Vogelstein, B. *et al.* Cancer genome landscapes. *Science* **339**, 1546-1558, doi:10.1126/science.1235122 (2013).

4 Lee, A. V., Oesterreich, S. & Davidson, N. E. MCF-7 cells--changing the course of breast cancer research and care for 45 years. *J Natl Cancer Inst* **107**, doi:10.1093/jnci/djv073 (2015).

5 Soule, H. D., Vazguez, J., Long, A., Albert, S. & Brennan, M. A human cell line from a pleural effusion derived from a breast carcinoma. *J Natl Cancer Inst* **51**, 1409-1416 (1973).

6 Keydar, I. *et al.* Establishment and characterization of a cell line of human breast carcinoma origin. *Eur J Cancer* **15**, 659-670 (1979).

7 Wosikowski, K. *et al.* Normal p53 status and function despite the development of drug resistance in human breast cancer cells. *Cell Growth Differ* **6**, 1395-1403 (1995).

8 Qu, Y. *et al.* Evaluation of MCF10A as a Reliable Model for Normal Human Mammary Epithelial Cells. *PLoS One* **10**, e0131285, doi:10.1371/journal.pone.0131285 (2015).

9 Soule, H. D. *et al.* Isolation and characterization of a spontaneously immortalized human breast epithelial cell line, MCF-10. *Cancer Res* **50**, 6075-6086 (1990).

10 Geltmeier, A. *et al.* Characterization of Dynamic Behaviour of MCF7 and MCF10A Cells in Ultrasonic Field Using Modal and Harmonic Analyses. *PLoS One* **10**, e0134999, doi:10.1371/journal.pone.0134999 (2015).

11 Thompson, E. A. *et al.* Differential response of MCF7, MDA-MB-231, and MCF 10A cells to hyperthermia, silver nanoparticles and silver nanoparticle-induced photothermal therapy. *Int J Hyperthermia* **30**, 312-323, doi:10.3109/02656736.2014.936051 (2014).

12 Tarangelo, A. & Dixon, S. J. Nanomedicine: An iron age for cancer therapy. *Nat Nanotechnol* **11**, 921-922, doi:10.1038/nnano.2016.199 (2016).

13 Savanur, M. A. *et al.* Sclerotium rolfsii lectin induces stronger inhibition of proliferation in human breast cancer cells than normal human mammary epithelial cells by induction of cell apoptosis. *PLoS One* **9**, e110107, doi:10.1371/journal.pone.0110107 (2014).

14 Bertram, C. & Hass, R. MMP-7 is involved in the aging of primary human mammary epithelial cells (HMEC). *Exp Gerontol* **43**, 209-217, doi:10.1016/j.exger.2007.11.007 (2008).

15 Dutta, S., Warshall, C., Bandyopadhyay, C., Dutta, D. & Chandran, B. Interactions between exosomes from breast cancer cells and primary mammary epithelial cells leads to generation of reactive oxygen species which induce DNA damage response, stabilization of p53 and autophagy in epithelial cells. *PLoS One* **9**, e97580, doi:10.1371/journal.pone.0097580 (2014).

16 Foster, K. A., Oster, C. G., Mayer, M. M., Avery, M. L. & Audus, K. L. Characterization of the A549 cell line as a type II pulmonary epithelial cell model for drug metabolism. *Exp Cell Res* **243**, 359-366, doi:10.1006/excr.1998.4172 (1998).

17 Nichols, W. W. *et al.* Characterization of a new human diploid cell strain, IMR-90. *Science* **196**, 60-63 (1977).

18 Fischer, K. R. *et al.* Epithelial-to-mesenchymal transition is not required for lung metastasis but contributes to chemoresistance. *Nature* **527**, 472-476, doi:10.1038/nature15748 (2015).

19 Thiery, J. P. Epithelial-mesenchymal transitions in tumour progression. *Nat Rev Cancer* **2**, 442-454, doi:10.1038/nrc822 (2002).

20 Horster, M. F., Braun, G. S. & Huber, S. M. Embryonic renal epithelia: induction, nephrogenesis, and cell differentiation. *Physiol Rev* **79**, 1157-1191 (1999).

21 Aokage, K. *et al.* Dynamic molecular changes associated with epithelial-mesenchymal transition and subsequent mesenchymal-epithelial transition in the early phase of metastatic tumor formation. *Int J Cancer* **128**, 1585-1595, doi:10.1002/ijc.25500 (2011).

22 Gregory, P. A. *et al.* The miR-200 family and miR-205 regulate epithelial to mesenchymal transition by targeting ZEB1 and SIP1. *Nat Cell Biol* **10**, 593-601, doi:10.1038/ncb1722 (2008).

23 Gibbons, D. L. *et al.* Contextual extracellular cues promote tumor cell EMT and metastasis by regulating miR-200 family expression. *Genes Dev* **23**, 2140-2151, doi:10.1101/gad.1820209 (2009).

24 Soltermann, A. *et al.* Prognostic significance of epithelial-mesenchymal and mesenchymal-epithelial transition protein expression in non-small cell lung cancer. *Clin Cancer Res* **14**, 7430-7437, doi:10.1158/1078-0432.CCR-08-0935 (2008).

25 Rho, J. K. *et al.* Epithelial to mesenchymal transition derived from repeated exposure to gefitinib determines the sensitivity to EGFR inhibitors in A549, a non-small cell lung cancer cell line. *Lung Cancer* **63**, 219-226, doi:10.1016/j.lungcan.2008.05.017 (2009).

26 Zavadil, J. & Bottinger, E. P. TGF-beta and epithelial-to-mesenchymal transitions. *Oncogene* **24**, 5764-5774, doi:10.1038/sj.onc.1208927 (2005).

27 Kang, J. H. *et al.* Aldehyde dehydrogenase is used by cancer cells for energy metabolism. *Exp Mol Med* **48**, e272, doi:10.1038/emm.2016.103 (2016).

28 Lee, J. S. *et al.* Dual targeting of glutaminase 1 and thymidylate synthase elicits death synergistically in NSCLC. *Cell Death Dis* **7**, e2511, doi:10.1038/cddis.2016.404 (2016).

29 Chuprin, A. *et al.* Cell fusion induced by ERVWE1 or measles virus causes cellular senescence. *Genes Dev* **27**, 2356-2366, doi:10.1101/gad.227512.113 (2013).

30 Li, J. *et al.* Inhibition of non-small cell lung cancer (NSCLC) growth by a novel small molecular inhibitor of EGFR. *Oncotarget* **6**, 6749-6761, doi:10.18632/oncotarget.3155 (2015).

31 Kim, J. J. *et al.* WSB1 overcomes oncogene-induced senescence by targeting ATM for degradation. *Cell Res* **27**, 274-293, doi:10.1038/cr.2016.148 (2017).

32 Mahale, J., Smagurauskaite, G., Brown, K., Thomas, A. & Howells, L. M. The role of stromal fibroblasts in lung carcinogenesis: A target for chemoprevention? *Int J Cancer* **138**, 30-44, doi:10.1002/ijc.29447 (2016).

33 Sacco, O. *et al.* Epithelial cells and fibroblasts: structural repair and remodelling in the airways. *Paediatr Respir Rev* **5 Suppl A**, S35-40 (2004).

34 Sima, J. & Gilbert, D. M. Complex correlations: replication timing and mutational landscapes during cancer and genome evolution. *Curr Opin Genet Dev* **25**, 93-100, doi:10.1016/j.gde.2013.11.022 (2014).

35 Rivera-Mulia, J. C. *et al.* Dynamic changes in replication timing and gene expression during lineage specification of human pluripotent stem cells. *Genome Res* **25**, 1091-1103, doi:10.1101/gr.187989.114 (2015).

36 Marchal, C. *et al.* Genome-wide analysis of replication timing by next-generation sequencing with E/L Repli-seq. *Nat Protoc* **13**, 819-839, doi:10.1038/nprot.2017.148 (2018).

37 Ryba, T., Battaglia, D., Pope, B. D., Hiratani, I. & Gilbert, D. M. Genome-scale analysis of replication timing: from bench to bioinformatics. *Nat Protoc* **6**, 870-895, doi:10.1038/nprot.2011.328 (2011).

38 Langmead, B. & Salzberg, S. L. Fast gapped-read alignment with Bowtie 2. *Nat Methods* **9**, 357-359, doi:10.1038/nmeth.1923 (2012).

39 Van Nostrand, E. L. *et al.* Robust transcriptome-wide discovery of RNA-binding protein binding sites with enhanced CLIP (eCLIP). *Nat Methods* **13**, 508-514, doi:10.1038/nmeth.3810 (2016).

40 Sethi, A. *et al.* A cross-organism framework for supervised enhancer prediction with epigenetic pattern recognition and targeted validation. *bioRxiv*, doi:10.1101/385237 (2018).

41 Arnold, C. D. *et al.* Genome-wide quantitative enhancer activity maps identified by STARR-seq. *Science* **339**, 1074-1077, doi:10.1126/science.1232542 (2013).

42 Cao, Q. *et al.* Reconstruction of enhancer-target networks in 935 samples of human primary cells, tissues and cell lines. *Nat Genet* **49**, 1428-1436, doi:10.1038/ng.3950 (2017).

43 Rao, S. S. *et al.* A 3D map of the human genome at kilobase resolution reveals principles of chromatin looping. *Cell* **159**, 1665-1680, doi:10.1016/j.cell.2014.11.021 (2014).

44 Barutcu, A. R. *et al.* Chromatin interaction analysis reveals changes in small chromosome and telomere clustering between epithelial and breast cancer cells. *Genome Biol* **16**, 214, doi:10.1186/s13059-015-0768-0 (2015).

45 Ay, F., Bailey, T. L. & Noble, W. S. Statistical confidence estimation for Hi-C data reveals regulatory chromatin contacts. *Genome Res* **24**, 999-1011, doi:10.1101/gr.160374.113 (2014).

46 Cheng, C., Min, R. & Gerstein, M. TIP: a probabilistic method for identifying transcription factor target genes from ChIP-seq binding profiles. *Bioinformatics* **27**, 3221-3227, doi:10.1093/bioinformatics/btr552 (2011).

47 Jiang, P., Freedman, M. L., Liu, J. S. & Liu, X. S. Inference of transcriptional regulation in cancers. *Proc Natl Acad Sci U S A* **112**, 7731-7736, doi:10.1073/pnas.1424272112 (2015).

48 Siepel, A. *et al.* Evolutionarily conserved elements in vertebrate, insect, worm, and yeast genomes. *Genome Res* **15**, 1034-1050, doi:10.1101/gr.3715005 (2005).

49 Ernst, J. & Kellis, M. ChromHMM: automating chromatin-state discovery and characterization. *Nat Methods* **9**, 215-216, doi:10.1038/nmeth.1906 (2012).

50 Han, H. *et al.* TRRUST v2: an expanded reference database of human and mouse transcriptional regulatory interactions. *Nucleic Acids Res* **46**, D380-D386, doi:10.1093/nar/gkx1013 (2018).

51 Milo, R. *et al.* Network motifs: simple building blocks of complex networks. *Science* **298**, 824-827, doi:10.1126/science.298.5594.824 (2002).

52 Neph, S. *et al.* Circuitry and dynamics of human transcription factor regulatory networks. *Cell* **150**, 1274-1286, doi:10.1016/j.cell.2012.04.040 (2012).

53 Lawrence, M. S. *et al.* Mutational heterogeneity in cancer and the search for new cancer-associated genes. *Nature* **499**, 214-218, doi:10.1038/nature12213 (2013).

54 Martincorena, I. *et al.* Universal Patterns of Selection in Cancer and Somatic Tissues. *Cell* **171**, 1029-1041 e1021, doi:10.1016/j.cell.2017.09.042 (2017).

55 Cheng, C. *et al.* An approach for determining and measuring network hierarchy applied to comparing the phosphorylome and the regulome. *Genome Biol* **16**, 63, doi:10.1186/s13059-015-0624-2 (2015).

56 Yang, E. *et al.* Decay rates of human mRNAs: correlation with functional characteristics and sequence attributes. *Genome Res* **13**, 1863-1872, doi:10.1101/gr.1272403 (2003).

57 Zhu, M., Liu, C. C. & Cheng, C. REACTIN: regulatory activity inference of transcription factors underlying human diseases with application to breast cancer. *BMC Genomics* **14**, 504, doi:10.1186/1471-2164-14-504 (2013).

58 Pereira, B. *et al.* The somatic mutation profiles of 2,433 breast cancers refines their genomic and transcriptomic landscapes. *Nat Commun* **7**, 11479, doi:10.1038/ncomms11479 (2016).

59 David M. Blei, A. Y. N., Michael I. Jordan. Latent Dirichlet Allocation. *The Journal of Machine Learning Research* **3**, 993-1022 (2003).

60 Blei, D. M., Ng, A. Y. & Jordan, M. I. Latent dirichlet allocation. *J. Mach. Learn. Res.* **3**, 993-1022 (2003).

61 Guo, Y. & Gifford, D. K. Modular combinatorial binding among human trans-acting factors reveals direct and indirect factor binding. *BMC Genomics* **18**, 45, doi:10.1186/s12864-016-3434-3 (2017).

62 Zhou, B. *et al.* Comprehensive, integrated and phased whole-genome analysis of the primary ENCODE cell line K562. *bioRxiv*, doi:10.1101/192344 (2018).

63 Dixon, J. *et al.* An Integrative Framework For Detecting Structural Variations In Cancer Genomes. *bioRxiv*, doi:10.1101/119651 (2017).

64 Li, H. *et al.* Reference component analysis of single-cell transcriptomes elucidates cellular heterogeneity in human colorectal tumors. *Nat Genet* **49**, 708-718, doi:10.1038/ng.3818 (2017).

65 Visvader, J. E. Cells of origin in cancer. *Nature* **469**, 314-322, doi:10.1038/nature09781 (2011).

66 Nik-Zainal, S. *et al.* Landscape of somatic mutations in 560 breast cancer whole-genome sequences. *Nature* **534**, 47-54, doi:10.1038/nature17676 (2016).

67 Pasqualucci, L. *et al.* BCL-6 mutations in normal germinal center B cells: evidence of somatic hypermutation acting outside Ig loci. *Proc Natl Acad Sci U S A* **95**, 11816-11821 (1998).

68 Migliazza, A. *et al.* Frequent somatic hypermutation of the 5' noncoding region of the BCL6 gene in B-cell lymphoma. *Proc Natl Acad Sci U S A* **92**, 12520-12524 (1995).

69 Harris, N. L. *et al.* New approaches to lymphoma diagnosis. *Hematology Am Soc Hematol Educ Program*, 194-220 (2001).
